# Hidden but Expanding Sampling Bias Distorts Prokaryotic Genome Collections

**DOI:** 10.64898/2026.09.29.755372

**Authors:** Hannah Götsch, Franz Baumdicker

**Affiliations:** Big Data Analytics in Bioinformatics, Institute for Bioinformatics, Justus-Liebig-University Gießen, Germany

## Abstract

Public genome collections underpin our understanding of bacterial diversity and the distribution of traits such as antibiotic resistance, and increasingly serve as training data for genomic foundation models. Their growing size promises an increasingly complete view of microbial genetic variation. However, these collections combine studies with different sampling priorities, often overrepresenting subgroups of closely related strains of particular interest. The resulting bias reduces effective information, and its consequences for diversity estimates and the value of further sequencing remain insufficiently quantified. Here we show that accounting for this escalating sampling bias is essential for accurate gene frequency estimates and hints at greater genomic diversity than current data suggest. We propose to detect and reduce sampling bias based on phylogenetic relationships compared to expectations under random sampling, which, in contrast to dereplication and data reduction methods, preserves fine-scale diversity in large samples. Applied to 402 prokaryotic species in the NCBI genome database, our method *PhyloThin* indicates that the effective number of samples is 41% lower than the actual number but retains more than twice as many genomes as dereplication. Sampling bias varies substantially among species, showing that genome counts alone provide an incomplete measure of how well a species has already been characterized. *PhyloThin* thus turns biased, expanding genome collections into more representative samples for studying genome evolution and population genetic inference with implications for training genomic foundation models, pathogen surveillance, and the discovery of novel genetic functions.

---

Biodiversity is fundamental for ecosystem functioning and resilience ^1^, making its monitoring essential to maintain ecological integrity. Yet robust methods to quantify and monitor biodiversity remain a central challenge in biology. Bacteria provide a powerful model system for addressing this issue ^2^. Public bacterial genome collections are increasingly used to estimate genetic diversity and the distribution of traits. Over the last decades, the number of completely sequenced bacterial genomes in repositories such as the *National Center for Biotechnology Information* (NCBI) of the *National Library of Medicine* has increased rapidly ^3^. Consequently, various analyses become computationally challenging or even infeasible when dealing with very large datasets ^4,5^. While machine learning methods such as large language models can process such enormous amounts of data, their performance is highly sensitive to sampling bias ^6,7^, which reduces effective information and can systematically distort conclusions ^8,9^. Thus, while redundant information should be avoided to lower the computational load, as many strains as necessary should be included to preserve information. Unbiased sampling, or reweighting the strains accordingly, is necessary to answer key questions such as: For which species would additional sampling of novel strains be most valuable? Which bacterial species are already comprehensively characterized in terms of their diversity? To answer these questions, naively using all the genomes deposited in databases is insufficient, as sampling bias in favor of certain lineages can lead to systematic underestimation of true biodiversity ^10^. However, most current databases are created from a multitude of sampling and data collection processes, where hidden biases obscure estimates. Their biological coverage depends on which strains are sequenced, not simply on the number of available genomes. This has far-reaching consequences, as genomic language models rely on unbiased and broad coverage of diversity, and in addition, bacterial biodiversity and pangenome size correlate with adaptive potential ^11^, and consequently with pathogenic capacity, pandemic risk, and the prevalence of antibiotic resistance.

Sampling bias arises whenever the selected samples fail to represent the composition of and the frequencies within the entire population. In surveys, careful planning is essential to ensure a known, ideally unbiased, sampling scheme, and the same principle applies to sequencing projects, as exemplified by the “1000 Genomes” project ^12^. In prokaryotic studies, random sampling is usually rare, either intentionally or due to practical constraints ^9^. Sampling bias can emerge from multiple sources: overrepresentation of easily cultivable strains or preferential sequencing of specific environments, e.g., clinical outbreaks. Additional biases occur during database submission or filtering, for example, when only completely sequenced genomes are retained or less diverse but medically relevant strains are preferentially included. Our study reveals another less obvious mechanism that generates sampling bias: Uncoordinated collection strategies of public databases, such as NCBI, can introduce substantial sampling bias through the combination of diverse genome collections, even when the individual studies sample their target subpopulations without bias.

Sampling bias takes several forms: some strains are uncultivable and thus missing or underrepresented in sequence data (similar to the “dark matter” challenge at species level ^13^), others arise from uniform sampling across major lineages ^9^, and the most common involves overrepresentation of closely related subgroups, which is our main focus here. Oversampling can range from being concentrated in a single closely related subgroup to being more evenly distributed across the sampled clusters (Fig. 4 A). Regardless of the concentration, we also refer to oversampling as sampling bias in the following.

A variety of tools exist to subsample large genomic datasets, most of which are phylogeny-based approaches that manually or automatically reduce the number of leaves ^14–17^. These methods typically rely on user-defined parameters, such as the desired number of remaining samples, and aim to preserve genetic diversity while removing samples. Although this can also lessen sampling bias, it does not explicitly correct for it and is therefore unlikely to eliminate it automatically. ANI-based dereplication methods ^18,19^ instead cluster closely related genomes using average nucleotide identity as a sequence-similarity threshold. Both approaches do not adequately test whether relatedness is excessive for the given sample size, although closely related genomes will become more likely as random samples grow.

Hence, while dampening the effect of sampling bias, datasets reduced by ANI or existing tree pruning methods do not represent a random sample of the population, which biases statistical analyses. An adaptive tool to detect sampling bias and, if necessary, reweight or reduce the dataset to its effective size is thus missing so far. Here, we present a method that addresses this gap. Considering phylogenetic relationships between genomic samples, our method detects sampling bias in prokaryotic populations, identifies oversampled genomes to reweight them or create a less biased effective subset that better mimics random sampling, a key assumption for many statistical methods.

We used our method *PhyloThin* to reveal and correct the prevalent sampling bias across prokaryotic genome collections at NCBI. *PhyloThin* transforms the heterogeneous and partly unstructured genome collections from 402 species into 402 reduced datasets, each approximating a more systematically and independently sampled collection. Downstream predictions based on the resulting effective and the uncorrected sample sets differ substantially. Without correction, gene frequencies will be over- or underestimated, and pangenomes can appear closed when they are in fact open, highlighting that sampling bias correction is essential to assess biodiversity, gene/trait prevalence, ancestral state reconstruction ^20^, selection forces, and the unsampled functional repertoire of prokaryotic species. Crucially, sampling biases and the resulting distortions in training data can propagate unnoticed into genomic foundation models ^6,7^.

## Results

### Adaptive sampling bias detection

In contrast to fixed ANI thresholds and basic tree pruning algorithms that trim datasets to a pre-specified number, the aim of our method *PhyloThin* is to identify sampling bias in an adaptive manner by allowing more closely related samples to be retained as the number of individuals increases. We achieve this by rescaling the coalescent times in the phylogenetic tree. In this rescaled tree, sampling bias causes detectably shorter cumulative sums of recent coalescent times than expected. We evaluate how the focus on the sum of the most recent coalescent times provides some resilience against artifacts caused by population structure or selection below and in the Methods. The principle is best explained in a constant population, which we assume in the following.

Under neutrality and unbiased sampling, coalescent times are identically and independently distributed after a proper rescaling on the coalescent time scale. In the removal step, *PhyloThin* tests whether the recent rescaled coalescent times together are shorter than expected under the reference distribution. If sampling bias is detected, the shortest external branch is marked as oversampled, and the reduced tree is reevaluated. This iterative process continues until no sampling bias can be detected. This process identifies overrepresented groups of closely related strains that require reweighting. Here, we retain one representative per group, but alternative schemes can distribute weights across all genomes in the group. The concept behind *PhyloThin* is illustrated in Fig. 1 and described in the Methods section. Coalescent simulations showed that *PhyloThin* detects both concentrated and distributed sampling bias, with performance depending on sample size, the extent of sampling bias and mutation resolution. These analyses also identified conditions leading to missed sampling bias or excessive removal. For example, the larger the sample size, the lower the resolution of the phylogenetic tree towards the tips. *PhyloThin* can account for this uncertainty using the mutation rate derived from the underlying alignment and a subsampling strategy, and reports its theoretically derived limits of sampling bias detection.

**Fig. 1.**
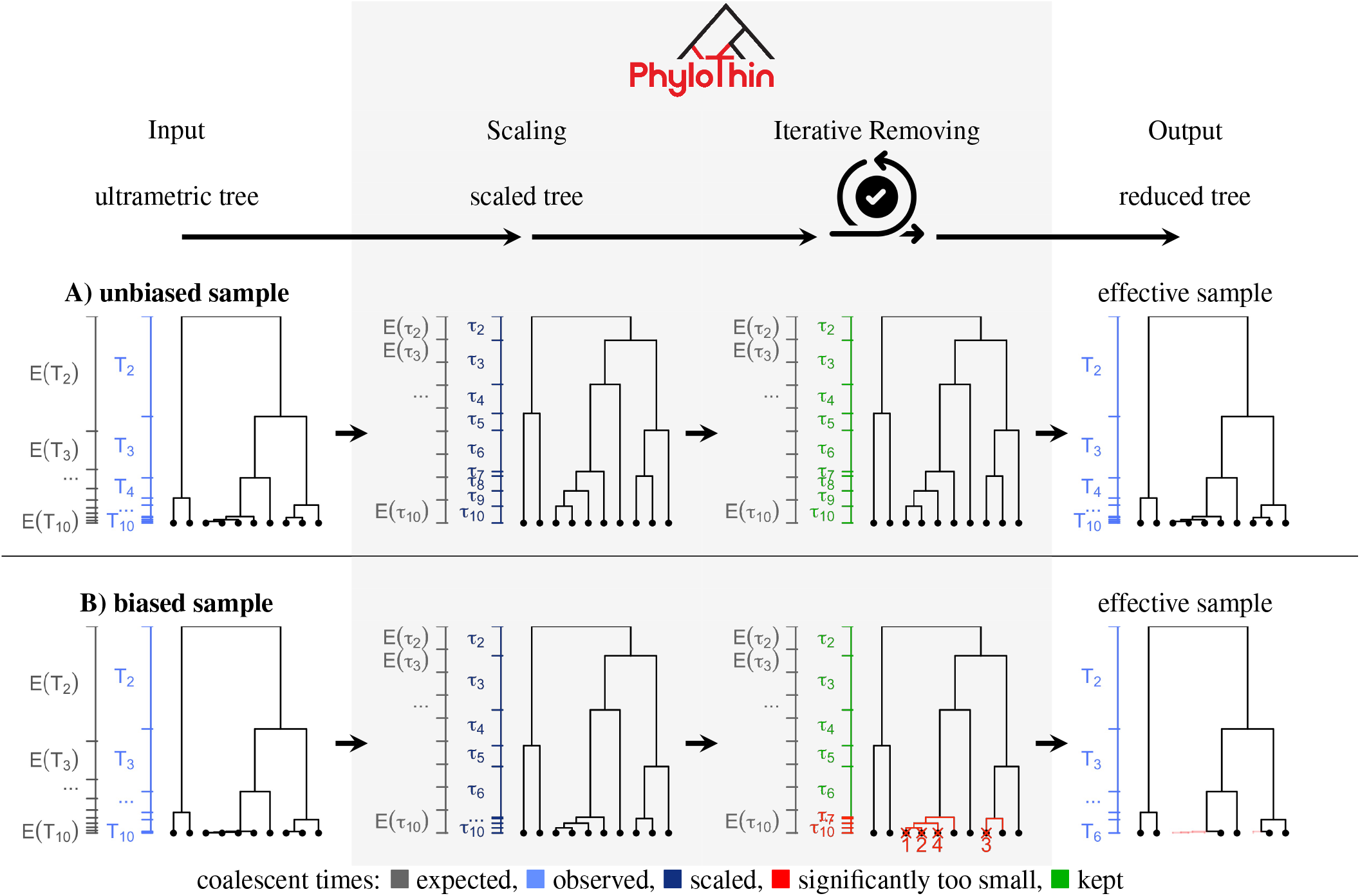
How *PhyloThin* identifies sampling bias. The first axis (gray, left side of each panel) shows the expected coalescent times of a neutral coalescent tree at each step of *PhyloThin*, representing the expectation of an unbiased sampling scheme. In the first row (A), variance is introduced to a neutral coalescent tree by drawing coalescent times from an exponential distribution. The second row (B) illustrates a biased sampling scheme, where four tips are oversampled in two groups. In the “removing” column, red crosses indicate the tips successively removed by *PhyloThin*. The real coalescent times (*T*_*i*_) and rescaled coalescent times (*τ*_*i*_) are shown on the left side of each tree (on the right side of the expected neutral coalescent times).

Given a phylogenetic tree, *PhyloThin* is fast. Runtime increases with the sample size and the amount of sampling bias. Among our datasets, analyzing sampling bias for the phylogenetic tree based on core genes of 3152 *E. coli* samples took the longest with 8 seconds on a standard laptop (15 seconds for mutation-rate-sensitive thinning). On the same dataset, activating the subsampling option without parallelization and using the recommended parameters to improve performance, increased the runtime to 36 minutes (21 minutes for subsampling and mutation-rate-sensitive thinning).

### Sampling bias in the NCBI database

The number of fully sequenced prokaryotic genomes on NCBI increased rapidly over the last years (Fig. 6 C). Both, the number of genome samples added each year (Fig. 6 A) and the proportion of genomes identified as oversampled (Fig. 6 B) increased, reflecting the impact of decreasing sequencing costs on database compositions. In particular, the proportion of oversampled genomes rose from 14% before 2008 to 25% by 2015, and has since increased steadily to 41% (Fig. 2). After applying *PhyloThin*, overall only 59% of the samples remain, reducing the 27, 070 analyzed full-genome samples to an effective number of 16, 100 genomes. However, despite the growing sampling bias, the effective number of genomes added each year continues to increase markedly (Fig. 6 A).

**Fig. 2.**
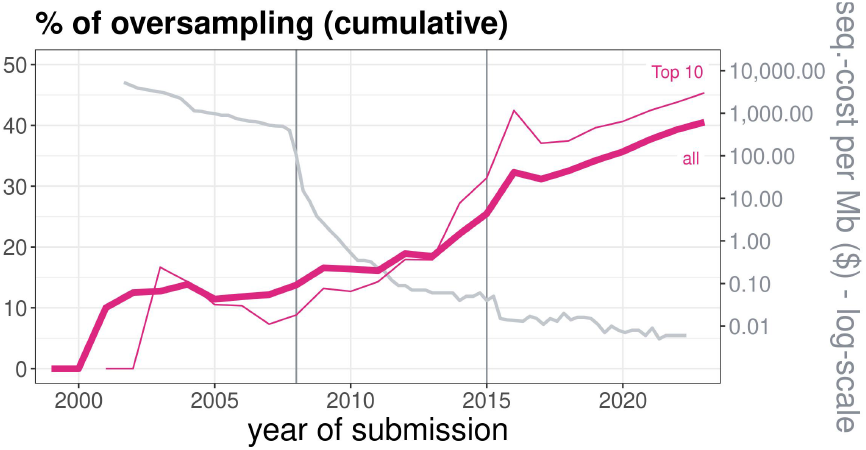
The cumulative percentage of oversampled genomes across bacterial genomes submitted to NCBI. An additional curve highlights the top ten species in the full dataset (table 1). The gray curve indicates sequencing cost per Mb ^3^. The gray vertical lines mark the two genomic revolutions: the advent of ‘second-generation’ sequencing (or NGS) in 2008, and a second strong drop in sequencing costs in 2015. We retrospectively assigned the samples detected as oversampled to their submission dates.

The extent of sampling bias varies among species (Fig. 8). Our adjustment for sampling bias reveals that the best represented species at the NCBI are not necessarily the ones with the most sequenced genomes. The ranking of the ten most abundant species shifts when considering the effective number of sampled genomes (table 1). While *Escherichia coli* remains the best represented species, despite substantial oversampling of 36.9%, *Salmonella enterica* shows with 86.6% the strongest oversampling among species represented by more than 70 genomes. *PhyloThin* retained 214 of 1, 602 *S. enterica* genomes, moving the species from third to tenth best represented species by effective sample size. This pronounced oversampling may reflect extensive genome sequencing associated with food surveillance and screening. Among the top species, *Bordetella pertussis* has the lowest oversampling rate making it the third largest effective genome set (593 genomes). *B. pertussis* also has a low diversity, which makes it an example of a well characterized species with an expected pangenome size increase of only 0.4% from the next 100 effective samples. In contrast, *S. enterica* would increase its pangenome size by 16.7%. Among species with more than 100 samples, *Vibrio cholerae* (*c*_*sb*_ = 0.68) has the highest sampling bias concentration (Fig. 4, Eq. (1)) and *Staphylococcus aureus* (*c*_*sb*_ = 0.001) has the most distributed sampling bias. Human-pathogenic species ^24^ tend to have larger sample sizes (Fig. 7). However, we detected no difference in oversampling fractions between species reported as pathogenic and non-pathogenic to humans (Fig. 8).

**Table 1.**
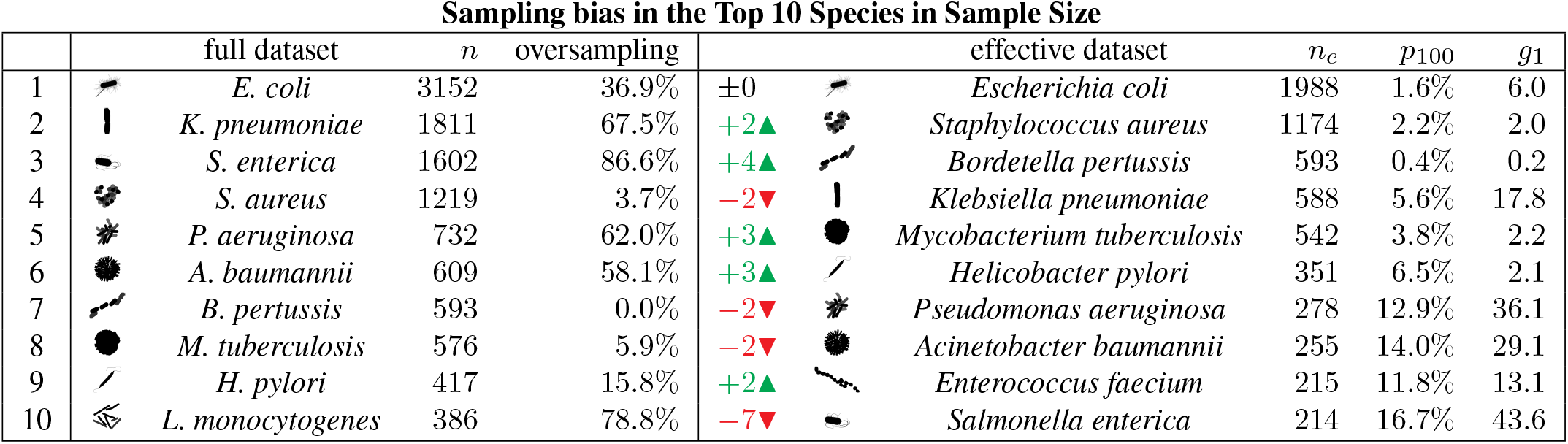
The top ten species in sample size change after sampling bias detection and correction. The table shows results for the full species-datasets (left) and effective species-datasets after thinning (right) by applying *PhyloThin* to phylogenetic trees based on core genes. For each species, the full sample size *n* in the database is given alongside the species name, and the percentage of oversampling. The metric *p*_100_ denotes the predicted increase in genes after unbiased sampling another 100 genomes and the metric *g*_1_ the predicted number of new genes expected from sequencing the next unbiased sampled genome. Silhouettes are from *PhyloPic* ^23^

|  |  | full dataset | *n* | oversampling |  |  | effective dataset | *n*<sub>*e*</sub> | *p*<sub>100</sub> | *g*<sub>1</sub> |
|---|---|---|---|---|---|---|---|---|---|---|
| 1 |  | *E. coli* | 3152 | 36.9% | ±0 |  | *Escherichia coli* | 1988 | 1.6% | 6.0 |
| 2 |  | *K. pneumoniae* | 1811 | 67.5% | +2▲ |  | *Staphylococcus aureus* | 1174 | 2.2% | 2.0 |
| 3 |  | *S. enterica* | 1602 | 86.6% | +4▲ |  | *Bordetella pertussis* | 593 | 0.4% | 0.2 |
| 4 |  | *S. aureus* | 1219 | 3.7% | −2▼ |  | *Klebsiella pneumoniae* | 588 | 5.6% | 17.8 |
| 5 |  | *P. aeruginosa* | 732 | 62.0% | +3▲ |  | *Mycobacterium tuberculosis* | 542 | 3.8% | 2.2 |
| 6 |  | *A. baumannii* | 609 | 58.1% | +3▲ |  | *Helicobacter pylori* | 351 | 6.5% | 2.1 |
| 7 |  | *B. pertussis* | 593 | 0.0% | −2▼ |  | *Pseudomonas aeruginosa* | 278 | 12.9% | 36.1 |
| 8 |  | *M. tuberculosis* | 576 | 5.9% | −2▼ |  | *Acinetobacter baumannii* | 255 | 14.0% | 29.1 |
| 9 |  | *H. pylori* | 417 | 15.8% | +2▲ |  | *Enterococcus faecium* | 215 | 11.8% | 13.1 |
| 10 |  | *L. monocytogenes* | 386 | 78.8% | −7▼ |  | *Salmonella enterica* | 214 | 16.7% | 43.6 |

Among genomes from the BioProjects included in this analysis, 46.3% were classified as oversampled, compared with 40.9% across all genomes. However, this changes significantly when *PhyloThin* is applied directly to the subtrees restricted to the BioProject-specific genomes (Fig. 5, Fig. 11 B), which resulted in a lower average proportion of 37.2% oversampling instead of 46.3%. The effect is even more pronounced for species in which more than half of the BioProject-specific samples have been removed in the full tree based analysis. Here the mean oversampling dropped from 82.2% to 49.1% in the BioProject-specific subtrees (Fig. 11 A). This suggests that sampling bias arises not only from biased sampling strategies and a focus on particular subtypes, but to a substantial extent from pooling genome samples of a multitude of independent less biased studies.

### ANI-based dereplication does not resolve oversam-pling

Comparing *PhyloThin*-results with dereplication based on fixed ANI thresholds reveals that no single fixed ANI threshold is suitable for all datasets to remove sampling bias (Fig. 3, Fig. 12 A). Here we consider two different ANI-cutoffs: 99.5%, suggested based on the observation that most of the NCBI-species show an ANI-gap between 99.2% and 99.8% 21, and, for comparison, 98%, which is the cutoff suggested if the goal of dereplication is to generate a set of genomes that are distinct when mapping short reads ^18^. As expected, the lower 98% cutoff would most of the time remove too many samples regardless of the number of samples, only 29% of all samples remain. For the 99.5% cutoff, sampling bias often remained undetected in datasets that have a low number of samples, while in large datasets with many initial samples, the same threshold became overly stringent, and would excessively remove diversity (Fig. 3, Fig. 12 B). However, despite these general patterns, dereplication using ANI varies substantially between species across the full range of sample sizes. In 41 species, *PhyloThin* retains 45.0% *±* 32.2% of the genomes that would have been removed by an ANI threshold of 99.5%; while in 332 species 86.6% *±* 24.1% of the oversampled genomes are not detected by the ANI threshold. Determining the appropriate species-specific ANI-threshold is thus a complex task that is automatically achieved when using *PhyloThin*.

**Fig. 3.**
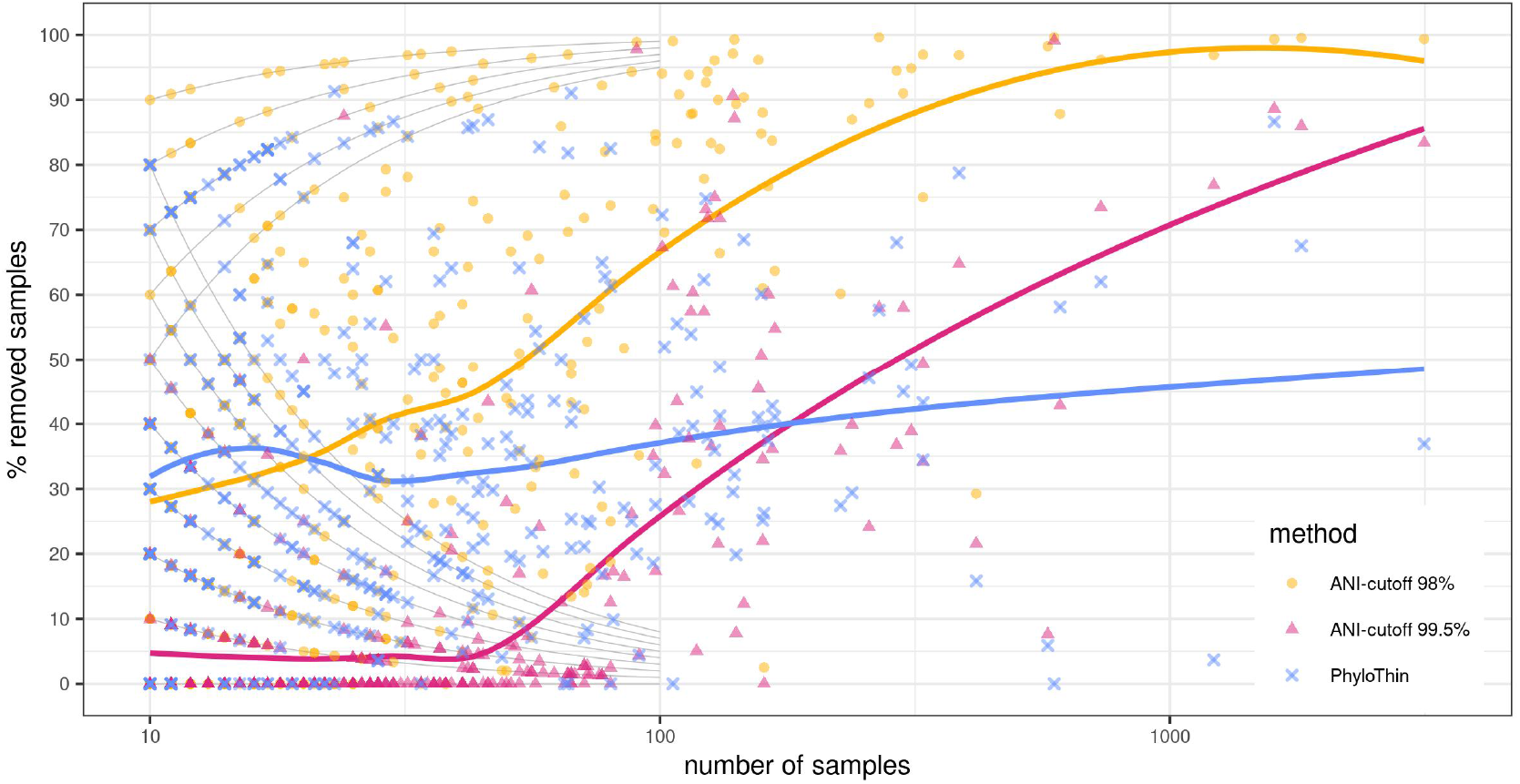
Comparison of ANI-based dereplication and *PhyloThin’s* sampling bias correction. The relationship between the proportion of removed genomes and the size of the sample set is shown. The different shapes and colors correspond to the method used to remove samples: 98% ANI-cutoff ^18^ (yellow circles), 99.5% ANI-cutoff ^21^ (pink triangles) and *PhyloThin* (blue crosses). The solid curves give the respective trend for the mean percentage for each sample size (LOESS smoothing ^22^). The gray curves connect points that correspond to the same absolute number of removed (1-8) or remaining (1-5) genomes.

### Sampling bias distorts population genetic analyses

Many population genetic analyses rely on gene or single-nucleotide polymorphism (SNP) frequencies within a population. Site and gene frequency spectra are important statistics in a range of applications, including selection detection and inference of ancestral population size ^26–28^.

Sampling bias can obscure the real distribution of SNPs and gene frequencies (Fig. 4 C) and, in the case of oversampling, often leads to a skewed distribution of rare genes and, for concentrated oversampling, also to an excess of genes at intermediate frequencies. For instance, in *Burkholderia cenocepacia* (Fig. 9a A), 54% of genomes belong to a single oversampled group (originating from NCBI-BioProject PRJEB40633^29^). As a result, the oversampled group produces an artificial peak of 525 group-specific genes at a frequency of 54%, and the frequency of those genes that are absent from the oversampled group is artificially reduced. How gene frequencies are distorted by sampling bias depends on the size of the oversampled groups. Consequently, datasets with concentrated sampling bias (Fig. 9a B) are affected differently from those with more distributed sampling bias (Fig. 9b B). In the *Burkholderia cenocepacia* example, a gene encoding an organic hydroperoxide resistance protein is absent from the oversampled group (together with 7 further genes), leading to an underestimation of its spread across the species (46% instead of 90%). In contrast, genes that encode efflux transporters and associated regulators (which can play important roles in antimicrobial resistance), for example, fall into the artificial frequency peak. Although their gene frequency in the retained effective dataset is only 10%, sampling bias inflates this value to 54%, leading to a potential misidentification as being under selection.

**Fig. 4.**
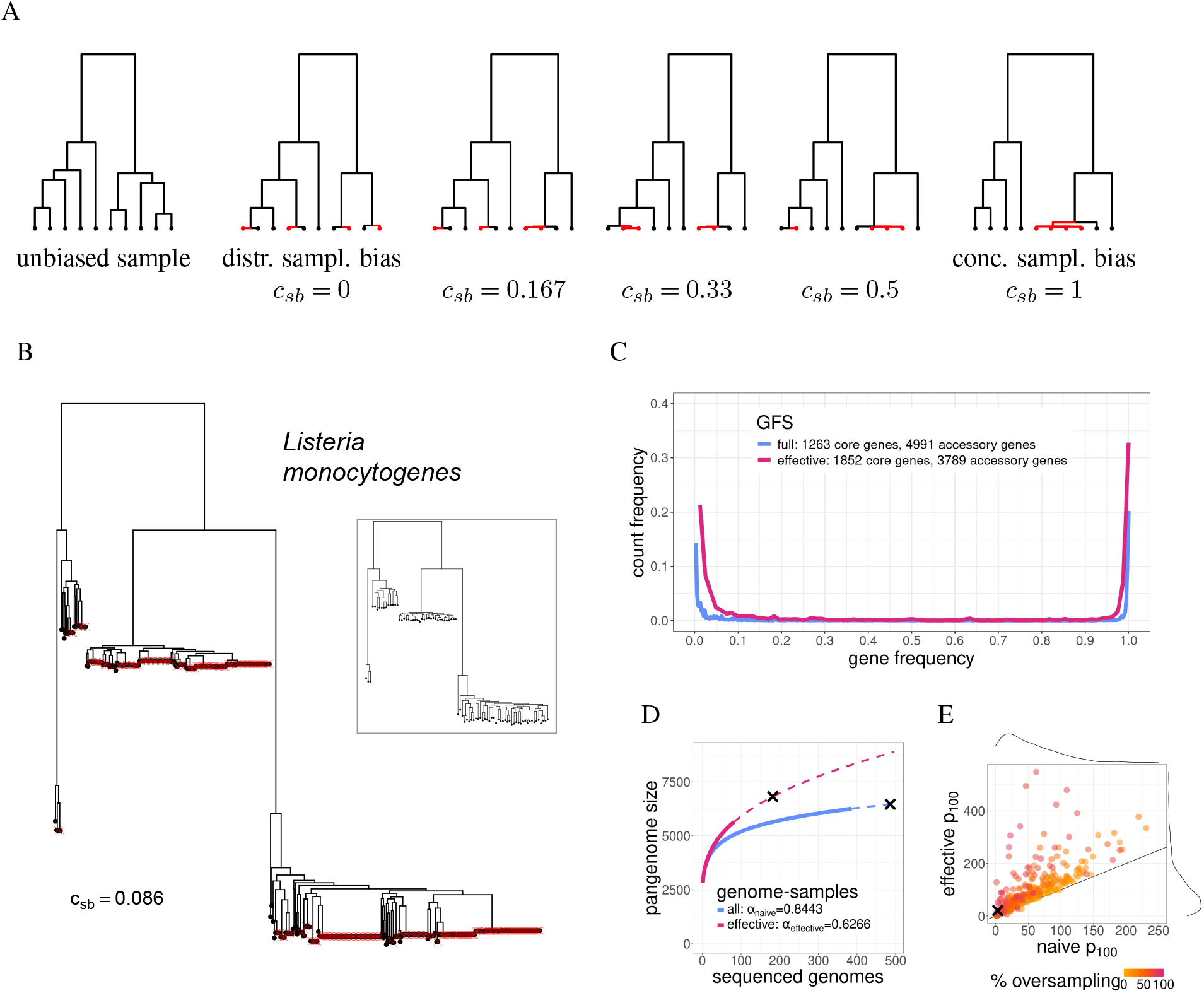
The impact of sampling bias. Panel A: **Concentrated and distributed sampling bias**. Schematic representations of inferred phylogenetic trees from unbiased sampling, and from biased sampling with concentrated and distributed sampling bias. The biased examples illustrate all possible configurations in which four of ten tips are oversampled. The oversampling concentration *c*_*sb*_ is defined in Eq. (1). Panel B: **Example sampling bias detection in *Listeria monocytogenes***. The full phylogenetic tree of *Listeria monocytogenes* based on core genes (386 tips) is displayed where the 304 removed tips are indicated as red crosses. The tree of the remaining strains after sampling bias correction is shown in the small box. Panel C and D: **Naive and effective gene frequency spectrum and pangenome accumulation curve**. The dashed curves in panel D give the predicted continuation of the pangenome accumulation curves obtained from a linear regression of the Heaps’ law on a log-log scale. The black crosses mark the predicted number of genes used to compute *p*_100_ (panel E). Comparable examples of concentrated and distributed sampling bias are shown in Fig. 9a and Fig. 9b. Panel E: **Predicted future pangenome size increase**. The percentage increase in newly detected genes after adding 100 genomes to the full dataset (naive *p*_100_) and effective dataset (effective *p*_100_), as predicted by Heaps’ law (extrapolated from the full and effective datasets), are compared for 364 species. The colors are chosen according to the percentage of oversampling. Sampling bias leads to an underestimate of the number of potentially newly detected genes (Wilcoxon signed-rank test with continuity correction: p-value < 2.2*e −* 16). The difference is larger for a high percentage of oversampling (Kendall’s rank correlation tau = *−*0.2956292, p-value < 2.2*e −* 16). The black cross represents *Listeria monocytogenes*.

### Distinguishing selective sweeps from oversampling

Identifying selection is a central task in population genetics. Selection pressures can shape phylogenetic trees such that internal branching times are shorter than expected ^30^. It is therefore essential that our correction method retains the true selection signals from a dataset, even if these resemble the effects of sampling bias. Vice versa, unless corrected by *PhyloThin*, selection detection methods can also misclassify a group of oversampled genomes as a genome-wide sweep. In simulations, selective sweeps did not affect *PhyloThin*’s results (Fig. 14b) for larger sample sizes unless they were genome-wide, very recent, and extremely strong (Fig. 14a). In a reanalysis of the isolate-genome-based phylogenetic trees corresponding to the 77 genome-wide sweep candidates reported by Yu et al. ^31^, *PhyloThin* detected 16.2% of the genomes as oversampling candidates, while retaining support for 68 of the 77 sweeps. For the remaining 9 inferred partial sweeps (11.7%), the phylogenetic trees indicated that oversampling could contribute to the sweep signal. However, sweep confirmation in Yu et al. ^31^ also incorporated metagenomic data and the small sample sizes of the analyzed trees (4 to 21) makes false positives more likely.

### Underestimating genomic diversity

Genomic diversity forms the basis for the adaptability and ecological versatility of a species. It is commonly assessed using pangenome accumulation curves, which visualize how the total gene repertoire expands as additional genomes are considered ^32^, and genomic fluidity, which describes the dissimilarity of genome pairs evaluated at the gene level ^33^. For both, accurate estimates require representative sampling to capture the true diversity of a species.

Sampling bias distorts pangenome accumulation curves (Fig. 10 A, C and E), particularly under high levels of oversampling (Fig. 10 A). The curve typically rises sharply at first, then plateaus (closed pangenome, *α >* 1) ^34^ or increases more gradually (open pangenome, *α <* 1) ^35^. When many too closely related genomes are among the samples, each genome contributes fewer novel genes than under unbiased sampling, leading to an underestimation of the diversity of the species. This produces artificially flattened pangenome accumulation curves, which, in some cases, even misleadingly suggest a closed pangenome. For example, for the human intestinal symbiont *Akkermansia muciniphila α*_naive_ = 1.11 but *α*_effective_ = 0.58, which changes its classification from closed to open.

The effect of sampling bias is stronger in datasets with concentrated sampling bias (Fig. 9a C) compared to those with more distributed bias (Fig. 9b C). Similarly, genomic fluidity is underestimated in biased datasets (Fig. 10 B, D, and F), especially for a high percentage of oversampling present in the dataset (Fig. 10 B) and in the case of concentrated sampling bias (Fig. 10 F). Applying *PhyloThin* to reduce sampling bias provides less biased diversity estimates, predicting stronger pangenome growth and higher genomic fluidity.

### Databases underestimate the capacity of prokaryotes to adapt

The extent of sequenced diversity within a species may underestimate its adaptive potential, as so far uncharacterized genes might be accessible via horizontal gene transfer and spread under selective pressures. Many rare accessory genes show signatures consistent with adaptive value and accessory-gene turnover is functionally and ecologically structured ^11,36^. Consequently, the extrapolated discovery rate of novel gene families can be viewed as a proxy for a species’ latent adaptive potential. We find that correcting for sampling bias allows more accurate forecasts of a species’ adaptive potential and helps to prioritize further sequencing efforts. Extrapolating from Heaps’ law, retrospective analysis shows that data available up to 2015 predict the retained effective gene counts of the 2023 pangenomes more accurately when corrected for sampling bias than when left uncorrected (on average 96% of genes in the pangenome are predicted vs. 87%, Fig. 13). As expected, sampling bias leads to an underestimate of newly detected genes, particularly in datasets with high levels of oversampling (Fig. 4 E). Corrected estimates for the number of expected novel genes and the fraction of so far unexplored repertoire of the pangenome vary substantially among the top 10 species. For *E. coli, S. aureus, B. pertussis* and *M. tuberculosis*, our analysis confirms that further sequencing effort will only reveal a few new genes. In contrast, despite the large genome collections already available for *S. enterica, P. aeruginosa* and *A. baumannii*, sequencing a relatively small number (*<* 100) of additional genomes could expand the known gene repertoire of each of these species by more than 10% (table 1).

The complete NCBI-dataset contains more than 2, 700, 000 genes across species (counting homologs once per species), 95% of which are retained after *PhyloThin* filtering. However, if the same number of genomes had been sampled without sampling bias, extrapolating from the corrected effective Heaps’ law functions predicts an increase of 13% in the number of sequenced genes, exceeding 3, 000, 000 in total. The diversity at the gene-presence level of prokaryotes is therefore currently underestimated.

## Discussion

Our results reveal that sampling bias is widespread and increasing in public databases (Fig. 2) such that bacterial biodiversity is systematically underestimated. In particular, merging data from various studies contributes to the often unnoticed sampling bias (Fig. 11). This effect is especially relevant in light of the current rise of genome foundation models. Removing this sampling bias is necessary to identify the effective amount of information and allows nondeceptive unbiased conclusions in approaches that rely on biological diversity, particularly for estimates of gene frequencies and adaptability of pathogens. With *PhyloThin*, we present a fully automated statistical tool to detect and correct sampling bias in prokaryotic populations while retaining fine-scale diversity for larger sample sizes. Although not considered here, the concepts of *PhyloThin* can in principle be extended beyond prokaryotes to phages, viruses, and unicellular eukaryotes, provided their recombination rates allow sufficiently resolved core genealogies. Accounting for sampling bias should be standard practice for large, as well as small, initial sample sizes, regardless of whether a reduction is necessary to effectively process the data, and enables the population genetic analysis of huge sample collections.

Bacterial genomes in public databases are not sampled randomly. Sequencing efforts are typically directed towards populations of societal, including clinical, epidemiological, or economic, interest ^9^. Human-pathogenic ^37^ species consequently tend to be represented by larger sample sizes (Fig. 7). Thus many pathogenic strains contribute to sampling bias, though we did not find a higher percentage of oversampling in total (Fig. 8). Nevertheless, particular phenotypes or lineages can be strongly enriched. For example, most complete genomes from *E. coli* are from strains pathogenic to humans, although *E. coli* is present in most warm-blooded animals ^9,38^. However, preferential representation does not necessarily result in the type of sampling bias we consider here, and the correlation between detected oversampling rates and increasing sample sizes is low (Fig. 8).

Instead, one possible explanation is that oversampling rates depend on how evenly the sampling efforts are distributed across the species’ ecological range. Generalists that are also prominent targets of genomic surveillance in our dataset have high oversampling fractions in our analysis. For *S. enterica* and *L. monocytogenes*, surveillance spans food, animal, clinical, and environmental sources, whereas genomic collections of *K. pneumoniae, P. aeruginosa*, and *A. baumannii* are strongly enriched for clinical isolates and epidemiologically important lineages. These sampling schemes repeatedly target only a subset of the species, and thereby generate too large clusters of closely related genomes. Consequently, despite a large number of sequenced genomes, much diversity can remain unsampled and underestimated, which is especially striking for *S. enterica* (table 1). However, targeted sequencing alone does not imply high oversampling: The diverse *S. aureus* is strongly represented by epidemic MRSA lineages, but shows very low oversampling in our analysis. Likewise, the rather clonally structured pathogens *B. pertussis* and *M. tuberculosis* have low oversampling fractions, although their underlying genetic diversity is quite limited. However, diversifying selection pressures of vaccines and a variety of vaccine escape variants might also influence the low oversampling rates in *B. pertussis*. Together, these observations suggest that sampling bias depends on the interaction between species-specific population structure and the way genomes are collected. In line with this explanation, the highly human-host-adapted species above are more evenly sampled than the more ubiquitous species, where sequencing efforts beyond the human context lag behind.

This dependence on sampling strategies and niche breadth also transfers to the level of the submitters of the sequences to NCBI. The *Centers for Disease Control and Prevention* (CDC) and *Food and Drug Administration* (FDA) upload the largest numbers of complete genomes in our dataset, with more than 1200 and 700 samples, respectively, of which approx. 18% and 48% are classified as oversampled by *PhyloThin*. These differences may reflect the specific surveillance aims and target species of the organizations. Sampling bias in public databases can thus arise at several stages: through targeted sampling within studies, through repeated sequencing of particular subpopulations, and through the subsequent accumulation of datasets generated for different purposes (Fig. 5 and Fig. 11).

**Fig. 5.**
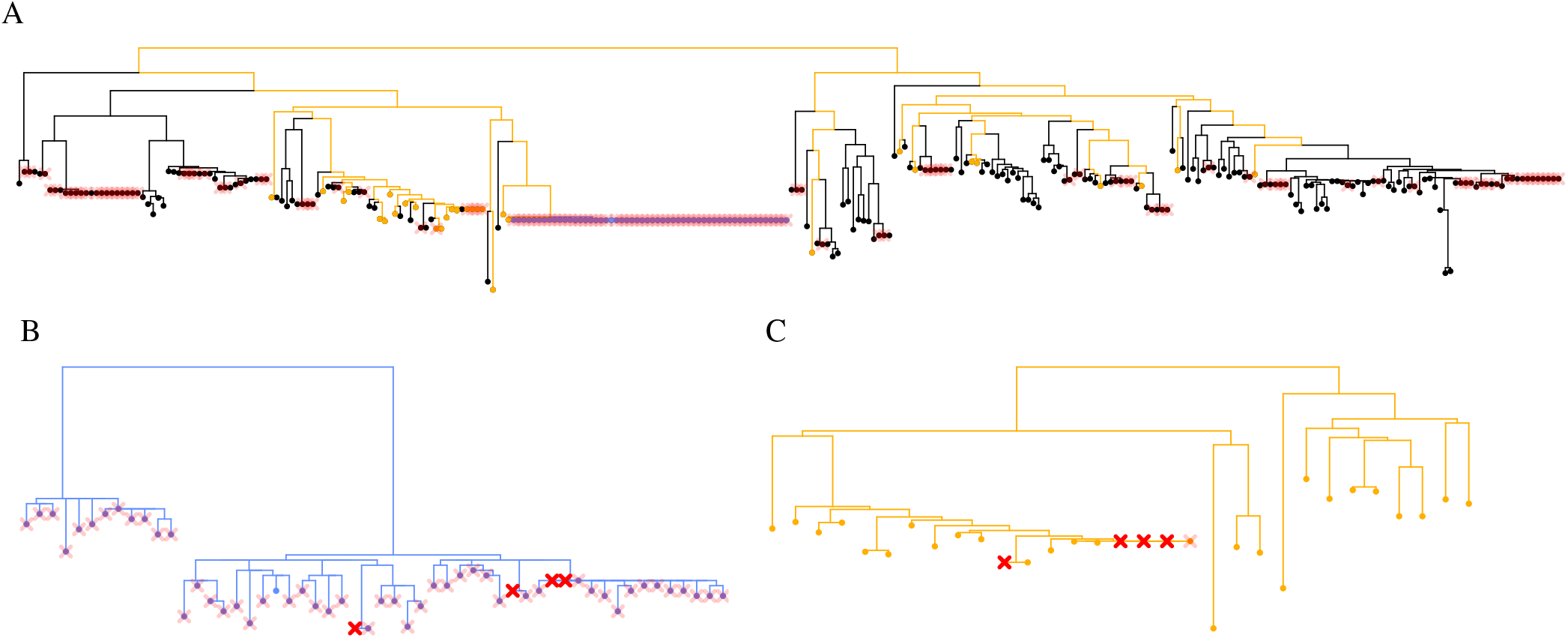
Sampling bias emerges by combining data from different less biased BioProjects. We show data for *Campylobacter jejuni* as an illustrative example. Panel A shows the full phylogenetic tree based on core genes (300 tips). Removed tips (135) are indicated as (light) red crosses (45% oversampling). The contributed genomes from BioProjects PRJNA268846 (blue, 54 tips, ^25^) and PRJNA344714 (yellow, 31 tips) and the corresponding subtrees are highlighted and shown as zoomed-in views in panels B and C, respectively. The darker red crosses in panel B and C indicate removed tips when *PhyloThin* has been applied to the subtree rather than the full tree (4 strains removed in each panel, 7.4% and 12.9% oversampling). The tips that generate additional oversampling at the level of the full dataset are indicated by light red crosses (53 in panel B and 5 in panel C) in the full tree.

Unless corrected, this sampling bias can have far-reaching consequences for downstream analyses and may propagate into genomic foundation models and other data-driven tools, including *AlphaFold* ^39^, whose performance relies on unbiased and sufficiently broad coverage of biological diversity ^7,40^. We show that sampling bias strongly affects estimates of species diversity and adaptability and cannot be adequately accounted for solely based on fixed ANI-thresholds (Fig. 3, Fig. 12). In particular, bias modifies pangenome accumulation curves (Fig. 4 D) which leads to an underestimation of pangenome diversity (Fig. 10 A and Fig. 4 E). But pangenome accumulation curves remain informative when sampling bias is appropriately considered.

Although alternative measures beyond pangenome growth can provide more robust insights ^8,33,41^, sampling bias correction is also essential to accurately infer diversity measures, such as genomic fluidity (Fig. 10 B). In fact, as sampling bias alters the gene frequency spectrum (Fig. 9a B), it propagates to all statistics derived from it. Common undesired effects that *PhyloThin* mitigates are (i) under- or overestimated spread and frequencies of genes of interest, in particular in the case of concentrated sampling bias, (ii) an apparent excess of rare variants that can provide false evidence for recent population growth, purifying selection, or genome-wide selective sweeps, as well as (iii) underestimated adaptive potential, where far more novel accessory genes could be found in further sequencing efforts than currently expected (Fig. 4 E).

Some downstream methods account for the phylogenetic relationships between samples and are therefore less sensitive to sampling bias. For example, the tool *Panstripe* ^8^ indirectly reduces the disproportionate influence of densely sampled parts of the tree when inferring gene gain and loss rates along a given phylogeny. Instead of removing individual strains and their information from downstream analyses, *PhyloThin* could be similarly used for reweighting, e.g. by rescaling the contribution of genomes by the size of identified oversampled groups they belong to. However, appropriate weights may require tool- and statistic-specific corrections. Removing sampling bias upstream provides a more general strategy across a broad range of downstream analyses and additionally reduces the computational costs.

In the last 20 years, gene sequencing costs have dropped substantially ^3^ and the number of sequenced genomes will continue to increase rapidly in the future (Fig. 6 A,C). Thus, we expect that sampling bias will increase further in the future and the need for methods to handle such a volume of data will increase ^42,43^. Our method benefits from the fact that inferring large phylogenetic trees from sequence data is already feasible, though challenges and uncertainties remain ^44,45^ and the uncertainties of tree estimation will impact our method. Once a phylogenetic tree is provided, *PhyloThin* is orders of magnitude faster than ANI-based methods. However, inferring a phylogeny from all core genes is computationally demanding, and trees based on a smaller, carefully chosen gene set may provide a practical compromise. While trees with higher resolution are recommended, *PhyloThin* accommodates any degree of diversity when using the mutation-rate-sensitive-thinning option that accounts for the tree uncertainty introduced by (low) mutation rates to prevent overestimating sampling bias. Low mutation rates might in turn not provide enough resolution to infer the shortest coalescent intervals. This may hinder the detection of sampling bias and eventually affects any given mutation rate as sample size increases. We recommend increasing the resolution by including more variable sites for tree inference whenever possible. For cases where this is not possible, *PhyloThin* offers a repeated subsampling strategy to obtain larger terminal coalescent intervals and thereby increase the resolution.

**Fig. 6.**
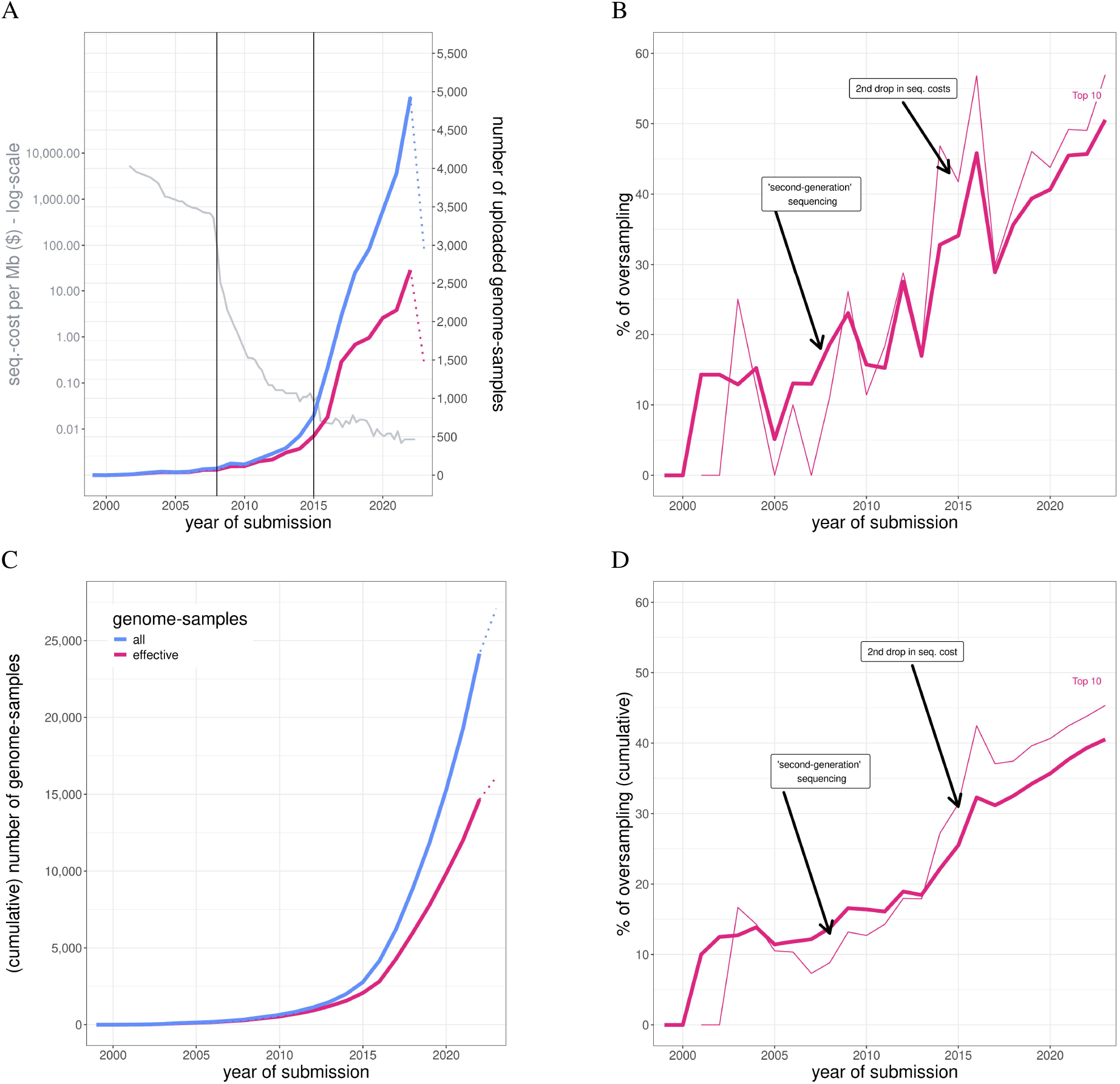
How much new data on NCBI gets sampled every year? The first row shows the development of the proportion of sampling bias in all analyzed bacteria species over time when we retrospectively assign the final oversampling classifications to submission dates. The second row shows the respective cumulative curves. The gray curve in panel A represents the sequencing cost per Mb ^3^. The black vertical lines (and arrows in panels B and D) represent the two genomic revolutions: ‘second-generation’ sequencing (or NGS) in 2008 and a second strong drop in sequencing costs in 2015. The kinks in the curve for the total number of genome samples and in the curve for the effective number of genome samples (identified through phylogeny based on the core genes) in 2023 occur because some samples from 2023 may be uploaded at a later time point and thus are not included in our dataset (panels A and C). In the second column, the percentage of oversampled genomes added each year (panel B) and the respective cumulative percentage over the years (panel D) are shown for results of the phylogenies based on core genes (settings of *PhyloThin* as described in the methods section). Additional curves are shown for the Top 10 of the full dataset (see table 1).

**Fig. 7.**
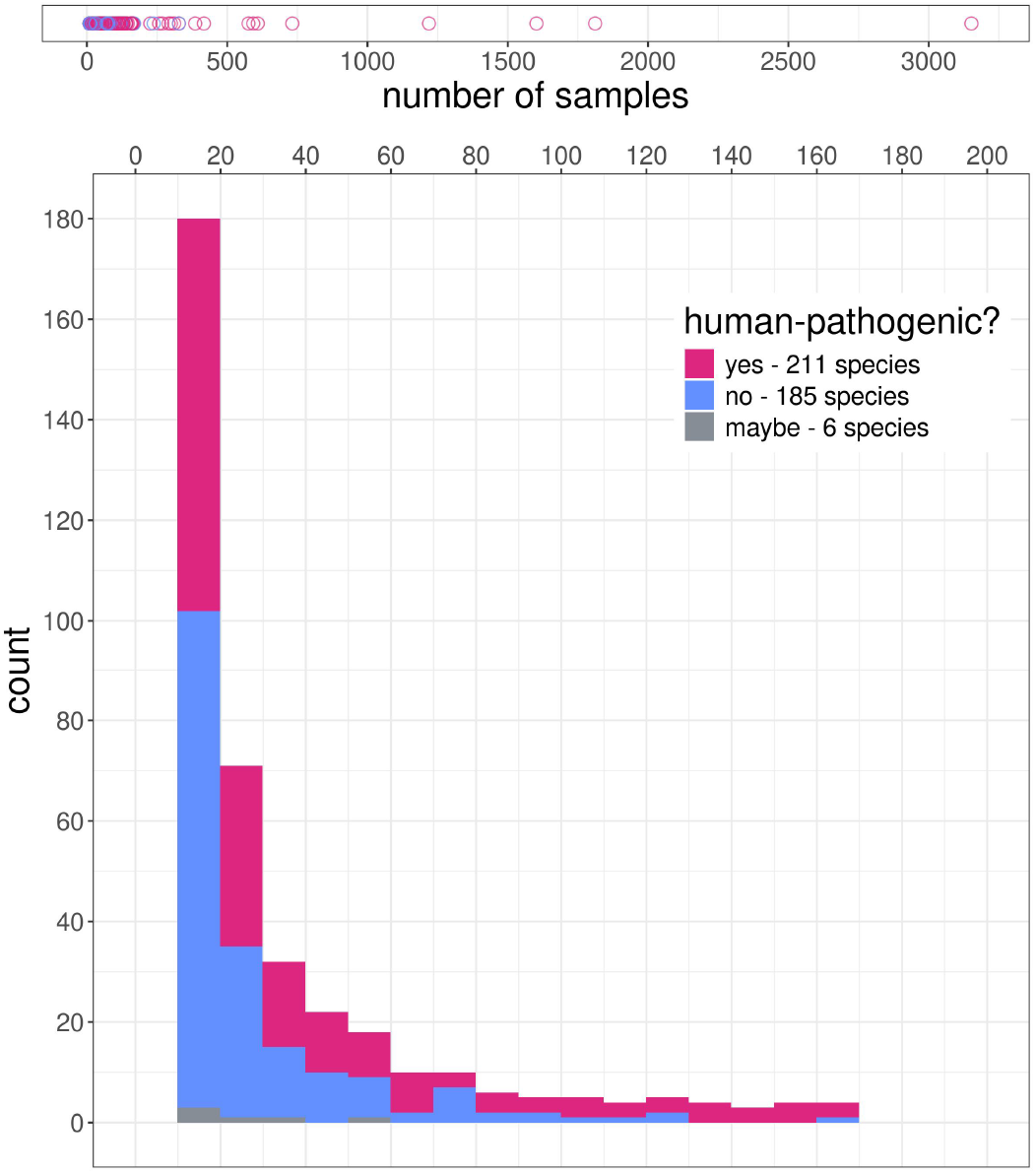
How large are the analyzed (at least 10 full-genome-entries) species sample sets? Total: 402 species, 27, 070 samples (August 4, 2023). The colors indicate the pathogenicity to humans of the species ^24^. Human-pathogenic species tend to have larger sample sizes (mean of 100.4 for pathogenic species vs. 30.9 for non-pathogenic species; Wilcoxon rank-sum test with continuity correction: p-value = 1.374*e −* 05).

**Fig. 8.**
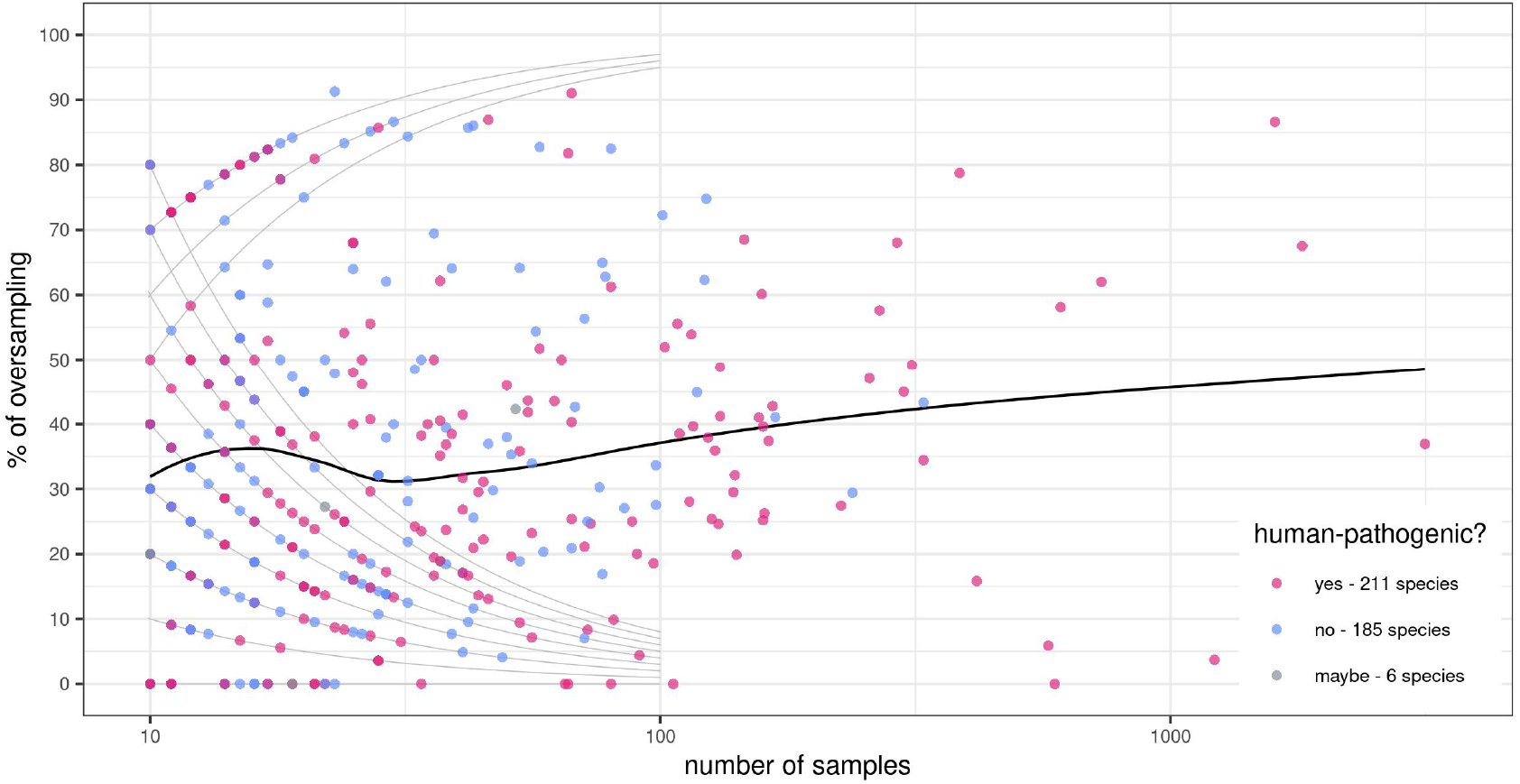
The relationship between the proportion of sampling bias and the size of the sample set. The black curve gives the trend for the mean percentage for each sample size (LOESS smoothing ^22^). The gray curves connect points that correspond to the same absolute number of oversampled or remaining genomes (L-shaped curves for 1 *−* 8 oversampled and upside-down L-shaped curves for 3 *−* 5 remaining). The species are categorized according to their pathogenicity to humans ^24^. The percentage of sampling bias is not significantly different for human-pathogenic than for human-non-pathogenic species (mean of 34.8% for pathogenic species vs. 35.2% for non-pathogenic species; Wilcoxon rank-sum test with continuity correction: p-value = 0.9835).

**Fig. 9.**
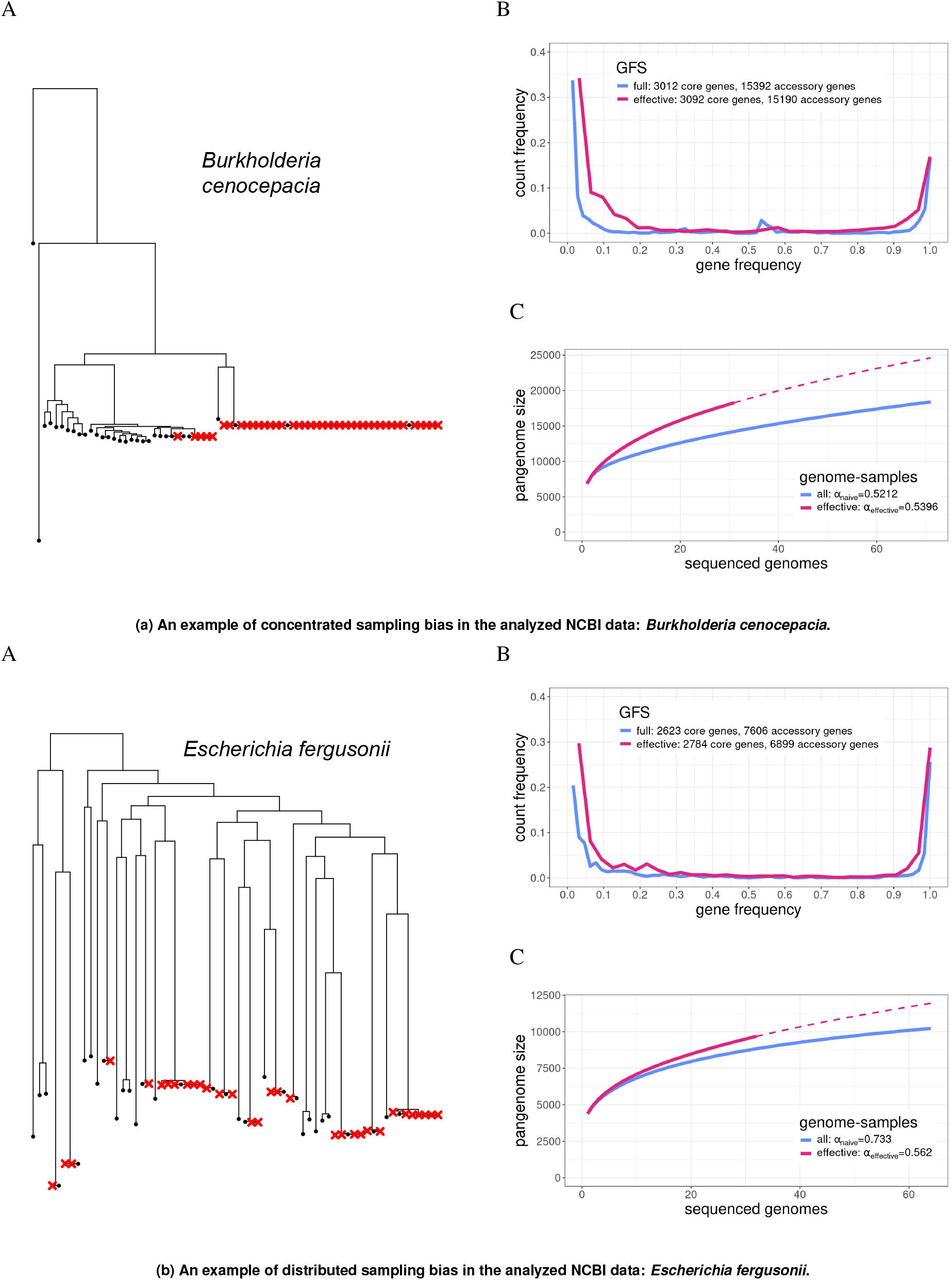
Examples of concentrated (a, *c*_*sb*_ = 0.73) and distributed (b, *c*_*sb*_ = 0.09) sampling bias in the analyzed NCBI data. In panels A, the full phylogenetic tree based on core genes (71 (a) and 64 (b) tips) is shown where removed tips (40 (a) and 32 (b)) are indicated as red crosses. In panels B and C, the full and effective GFS and pangenome accumulation curve are shown, respectively. The dashed curve in panels C gives the predicted continuation of the effective pangenome accumulation curve obtained from a linear regression of the Heaps’ law on a log-log scale.

**Fig. 10.**
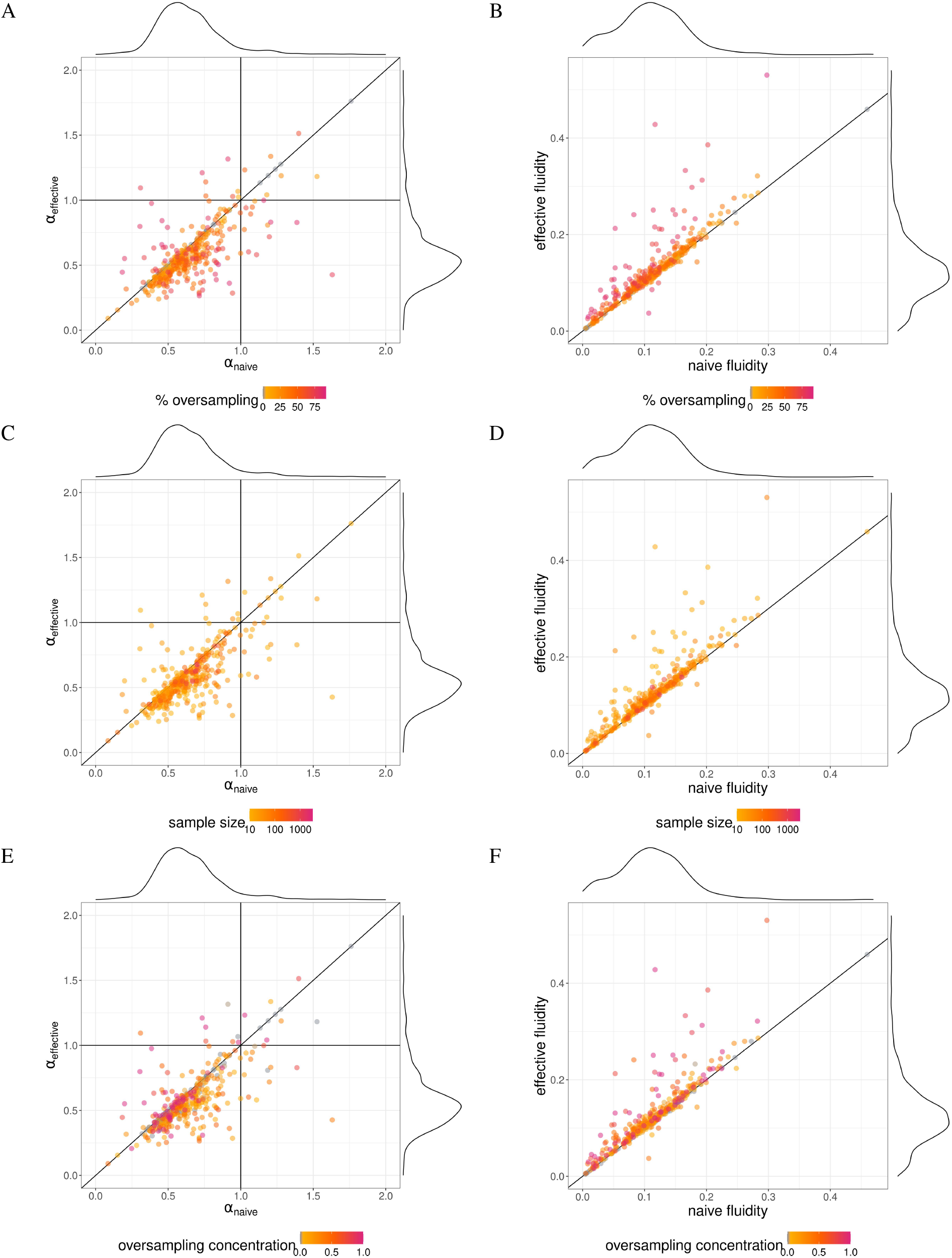
Impact of sampling bias on pangenome*α*-parameters and genomic fluidity. Panel A, C and E: The *α*-parameters of the full (*α*_naive_) and effective (*α*_effective_) dataset are compared for each species. Parameters are inferred from Heaps’ law (*g* = *cG*^*−α*^, where *g* is the number of new genes in the pangenome, *G* is the current number of genomes, and *c* and *α* are constants defined to fit the formula ^70^). Open pangenomes are characterized by *α <* 1 ^32^ and closed pangenomes by *α >* 1 ^34^. Sampling bias causes deviations in *α* (Wilcoxon signed-rank test with continuity correction: p-value < 2.2*e −* 16). The difference is larger for a higher percentage of oversampling (panel A, Kendall’s rank correlation tau = 0.2411707, p-value = 1.15*e −* 12).

As an alternative to core gene trees one might be tempted to also consider trees derived from accessory gene presence-absence states. However, one has to keep in mind that gene presence–absence trees can have expanded terminal branches compared to core gene trees ^46^, and might not be adequately described by standard coalescent theory. In addition, gene presence-absence variation is shaped by selective forces on accessory genes ^47^, as well as by incompletely sequenced genomes and plasmids. We argue that for the purpose of our tool, core-gene trees are more reliable in terms of sampling bias detection, but careful tree inference that is capable of reconstructing coalescent times is crucial.

The assumption that the phylogenetic trees follow a standard coalescent process is a simplification, as the phylogenies are shaped not only by clonal inheritance and linkage but also by population structure and selection. However, our approach is defensible for our task because sampling bias predominantly arises at recent timescales (Fig. 14a), which makes its inference robust against deviations from the coalescent at more ancient times. Unaccounted population structure would distort the overall coalescent time calibration, but a wrongly estimated scaling factor would still allow to detect the extremely short sums of terminal coalescent times. Similarly, selection is unlikely to substantially distort these recent phylogenetic patterns, as it typically changes (at least slightly) more ancient internal branching times ^30^ compared to the very recently occurring sampling bias. Also, adaptive sweeps in bacterial populations tend to involve genes rather than entire genomes ^48^. For genome-wide selective sweeps (or by analogy a recent rapidly expanding neutral subpopulation) the picture might be different, but our simulations showed that *PhyloThin* does not misclassify them as sampling bias unless the effect is strongly pronounced, very recent, and fast (Fig. 14b), a regime that requires selection coefficients higher than those typically measured ^49,50^.

*PhyloThin* enables researchers to account for oversampled lineages in prokaryotic datasets and prevent far-reaching consequences. This enables more accurate estimation of strain diversity which is crucial for pathogen control ^51^ and helps to identify those species that seem to be, but are actually not yet, sufficiently represented in the databases. Sampling bias detection thus helps to guide future sequencing efforts towards so far unexplored, potentially valuable genetic variation and strains ^52^ and improves the assessment of existing genomic resources. This is particularly relevant for genomic foundation models, for which phylogenetic dependencies among training genomes can confound the distinction between shared ancestry and functional constraint ^7,40^. Recognizing and correcting for these biases is therefore essential to fully capture the ecological and evolutionary potential of prokaryotic life. When we account for the escalating prevalence and emergence of sampling bias in the huge genomic datasets generated, we can take advantage of the sequencing efforts across the world and correctly infer bacterial diversity, genomic fluidity, and epidemic risks.

## Supporting information

Supplementary Figures NCBI species

Redme-file for Supplementary Table NCBI species

Supplementary Table NCBI species

## Methods

### PhyloThin

We present *PhyloThin* (Fig. 1), a deterministic, fully automated statistical tool written in *R* ^53^. It starts with an ultrametric phylogenetic tree, i.e., all tips are equidistant from the root. Afterwards, the coalescent times of the tree are scaled in order to allow for testing for sampling bias. Oversampled tips are then removed iteratively.

In the Kingman coalescent model on coalescent time scale, the times *T*_*i*_ - during which there were exactly *i* lineages ancestral to the sample - follow an exponential distribution, 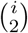 *T* ~ Exp(1) ^54^. Because phylogenetic trees are usually given in mutation units, they must first be rescaled. Weestimate a scaling factor from a subset of coalescent intervals near the root - identified using a tailored statistic - and use it to convert all branch lengths to coalescent time units. A further transformation by 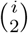 ensures that the intervals follow the expected exponential distribution, allowing us to detect unusually short external branches that signal sampling bias. *PhyloThin* iteratively removes the tip with the shortest external branch until no further signal of oversampling is detected by our method. To increase sensitivity, the method considers cumulative sums (up to some tailored threshold) of the external coalescent intervals, which follow a gamma distribution under neutrality. Deviations from this expectation reveal oversampled tips.

Phenomena such as recent fast genome-wide selective sweeps, and the neutral equivalent of recent fast population growth ^55^, may generate large groups with short external branches, which risk being disproportionately removed. The fully automated approach of *PhyloThin* to infer the number of kept samples might minimize this problem. Only strongly pronounced effects of such processes can still lead to misclassification of sampling bias and removal of the corresponding samples (Fig. 14b).

Another consideration is that long branches may result from low-quality or misaligned sequences. By transforming trees into ultrametric form, the effect of such long branches is distributed across all lineages in the relevant time frame, reducing the risk of enriching for poor-quality sequences. In some extreme cases, long branches may stop the bias-reduction procedure in *PhyloThin* prematurely. We therefore recommend that users pre-check sequence quality and inferred phylogenies ^56^ before using *PhyloThin* in their pipeline. To further minimize the risk of early termination, *PhyloThin* does not rely solely on the shortest external branch (the first coalescent interval), but instead considers sums of coalescent intervals during its iterative removal step.

The full approach of *PhyloThin* with all technical details is given in Sup. Note 1. Various tuning parameters were optimized with the help of simulations (Sup. Note 3). The user is provided with an example for the reduced tree based on an optional priority list ranking the strains for the cases where the remaining genome has to be selected among too closely related strains as well as which genome samples belong to the same oversampled group if reweighting of all the genomes in the same group is preferred. If the given phylogenetic tree is not ultrametric, *PATHd8* ^57^ is used as a first step. However, we highly recommend to use a method that uses a coalescent based process to modify the branch lengths, such as *TreeTime* ^58^ instead, to avoid an inflation of terminal branch lengths that sometimes occurs in *PATHd8*. If the dataset remains too large for a particular analysis after sampling bias detection with *PhyloThin*, (random) subsets ^59^ or tools such as *Treemmer* ^60^ can be used to further reduce its size.

### Low mutation rate adjustments

For low mutation rates, branches of length zero become more frequent, especially in large phylogenetic trees. This may violate the assumptions underlying the default settings of *PhyloThin* (external branches of length zero will always be removed in the limit of “infinite” mutation rate) and lead to an overestimation of sampling bias. We therefore recommend ensuring that a sufficient number of variable sites has been used for tree inference (e.g. by using the soft core genome) or, where this is not possible, cross-checking the results against those of the mutation-rate-sensitive version of *PhyloThin*, which explicitly accounts for an inferred mutation rate when testing the most recent coalescent times. Since mutation-rate-sensitive thinning may underestimate sampling bias and if increasing the resolution of the tree is not practical, we suggest combining mutation-rate-sensitive thinning with subsampling, which increases the chance of identifying concentrated oversampling clusters.

### Large sample size adjustments

When considering more and more samples, the expected number of differences between the most closely related individuals will drop, possibly up to the point where the mutation rate is not high enough to allow a reliable inference of the shortest coalescent times. We circumvent this issue by subsampling the strain collection multiple times. In each smaller subsample mutations translate more reliably into coalescent times. By considering the ratio of subsamples where a strain has been identified as oversampled we can then identify oversampling in the complete dataset. The sensitivity of this approach depends on how concentrated the true oversampling is and works better for more concentrated sampling bias.

### Analysis of the NCBI database - Data preprocessing

We analyzed with *PhyloThin* all bacterial species taxids from the NCBI database with at least 10 full-genome-entries (August 4, 2023). The genome-samples have been downloaded using *ncbi-genome-download* ^61^. The corresponding phylogenetic trees were then reconstructed from the core-genes of the genome-samples by *RAxML* ^62^ in the pangenome analysis tool *panX* ^63^. In addition to the core gene strain tree from *panX*, we also applied the “BIN+G” model (inferences on binary data and discrete-GAMMA among-site rate heterogeneity model) of *RAxML-NG* ^64^ on the gene presence absence matrix. Both methods use a maximum-likelihood optimality criterion to infer the phylogenetic tree.

The phylogenetic trees inferred from the core genes were made ultrametric with *TreeTime* ^58^, assigning the same sampling date to all tips to enforce ultrametricity. Starting from the 100% core genome trees (402 species), we applied *PhyloThin* with the mutation-rate-sensitive-thinning option. To assess whether the underlying alignment resolves closely related strains well, we computed *k* (Eq. (14)), the size of the smallest cluster of samples at pairwise distance zero that mutation-rate-sensitive thinning can still detect; a large *k* relative to the sample size indicates poor resolution. For species with *k >* 2 (188 species) and a ratio of *k* to sample size above 0.1 (92 species) we therefore proceeded as follows. For fewer than 100 samples (75 species) the additional subsampling step of *PhyloThin* was applied, which identified further samples as oversampled in 13 cases. For at least 100 samples (17 species) we used the tree reconstructed from the 99% soft core genome, which provides a higher resolution due to the larger alignment; recomputed *k*, applying the additional subsampling step of *PhyloThin* in the 9 species for which the ratio still exceeded 0.1.

Applying *PhyloThin*, we prioritized reference sequences and older genome samples for keeping. From the analyzed 27, 070 samples, 373 samples from 372 species taxids (*E. coli* has two reference sequences) are categorized as reference sequences.

To assess how strongly sampling bias depends on individual sequencing projects, we analyzed species and their associated BioProjects that each contributed at least 20 genomes. This yielded 140 BioProjects spanning 71 species and 154 subtrees (Fig. 11).

**Fig. 11.**
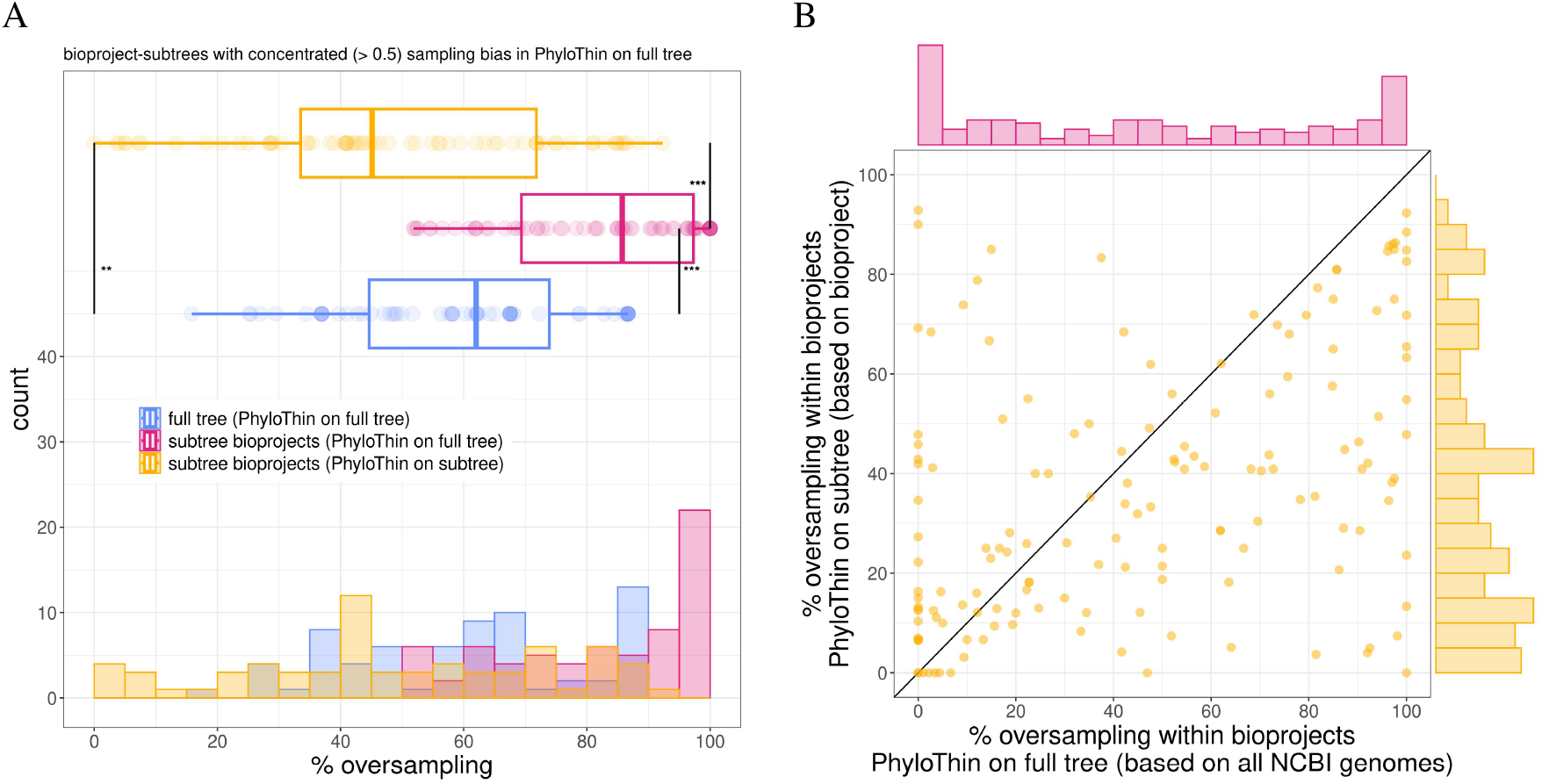
Sampling bias in BioProject-subtrees versus full species trees. We analyzed 140 BioProjects across 71 species, comprising 154 subtrees, each with at least 20 samples submitted by the same BioProject. In panel A, we compared the percentage of oversampling for BioProject-subtrees (68 subtrees) where more than 50% of the subtree has been removed by *PhyloThin* in the full tree. The percentage of oversampling in full species trees (blue, mean 60.0%), BioProject-subtrees analyzed using the full tree (pink, mean 82.2%), and BioProject-subtrees analyzed independently with *PhyloThin* (yellow, mean 49.1%) are shown. All deviations are significant (Wilcoxon signed-rank test with continuity correction for full trees and subtrees (*PhyloThin* on full tree): p-value = 4.108*e −* 11, for full trees and adjusted subtrees (*PhyloThin* on subtree): p-value = 0.003864, for subtrees (*PhyloThin* on full tree) and adjusted subtrees (*PhyloThin* on subtree): p-value = 1.372*e −* 12). In panel B, the percentage of oversampling in all BioProject-subtrees when *PhyloThin* is applied to the full tree (horizontal axis, corresponds to pink in panel A) is compared to the percentage, when *PhyloThin* has been applied to the BioProject-subtree (vertical axis, corresponds to yellow in panel A).

**Fig. 12.**
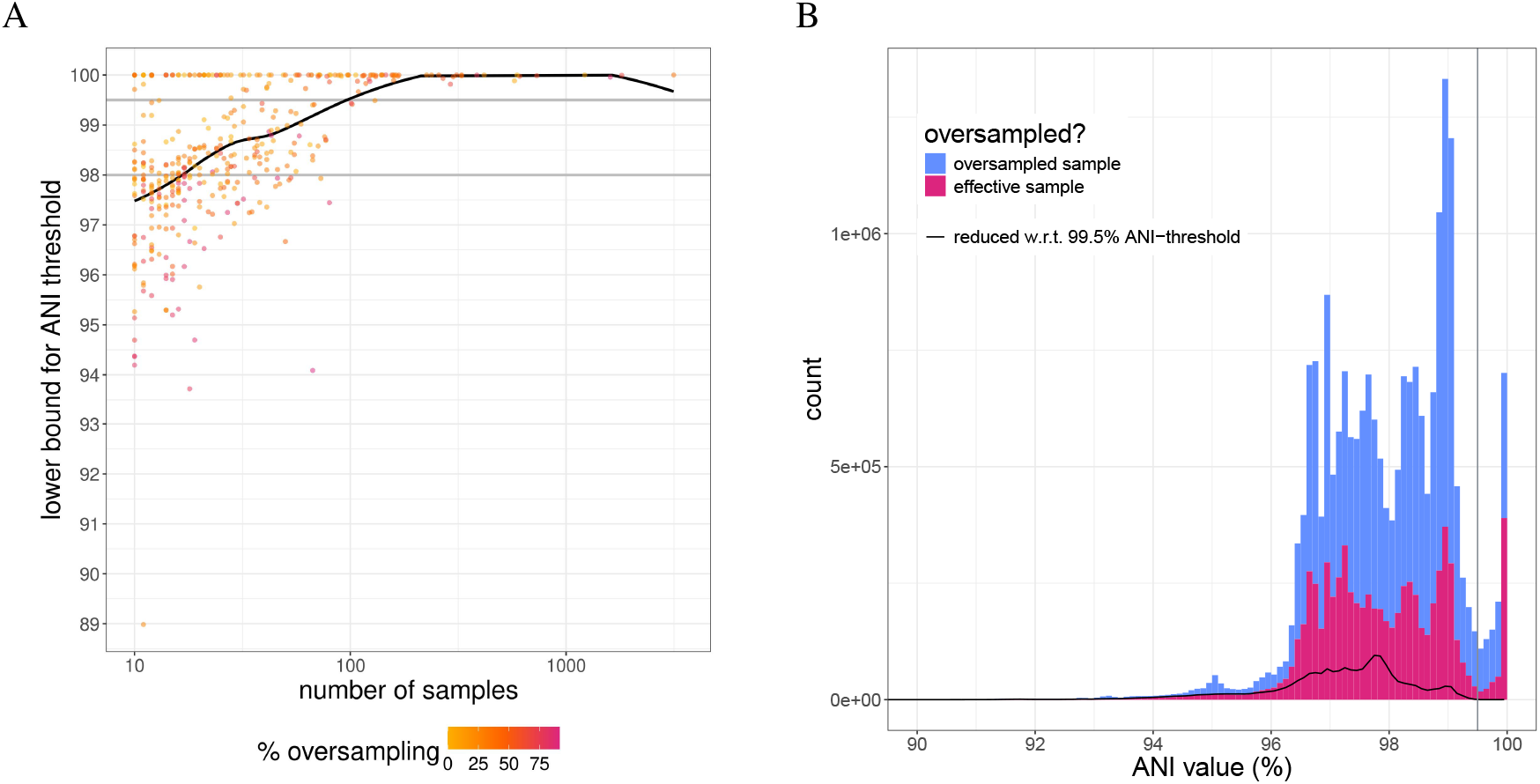
Relationship between ANI threshold and removal of sampling bias. In panel A, the lower bound of a suitable ANI threshold to remove sampling bias (the largest ANI value present in the effective dataset after applying *PhyloThin*) increases with sample size. Points are colored by the percentage of oversampling, and the black curve shows the overall trend (LOESS smoothing ^22^). Gray horizontal lines mark commonly used ANI thresholds: 98% ^18^ and 99.5% ^21^. In panel B, the histogram of (within species) ANI values for all species is shown. Pink bars represent ANI values retained after *PhyloThin*, and blue bars indicate ANI values from oversampled genomes. The black curve shows the histogram obtained using a 99.5% ANI-threshold ^21^ (gray line).

**Fig. 13.**
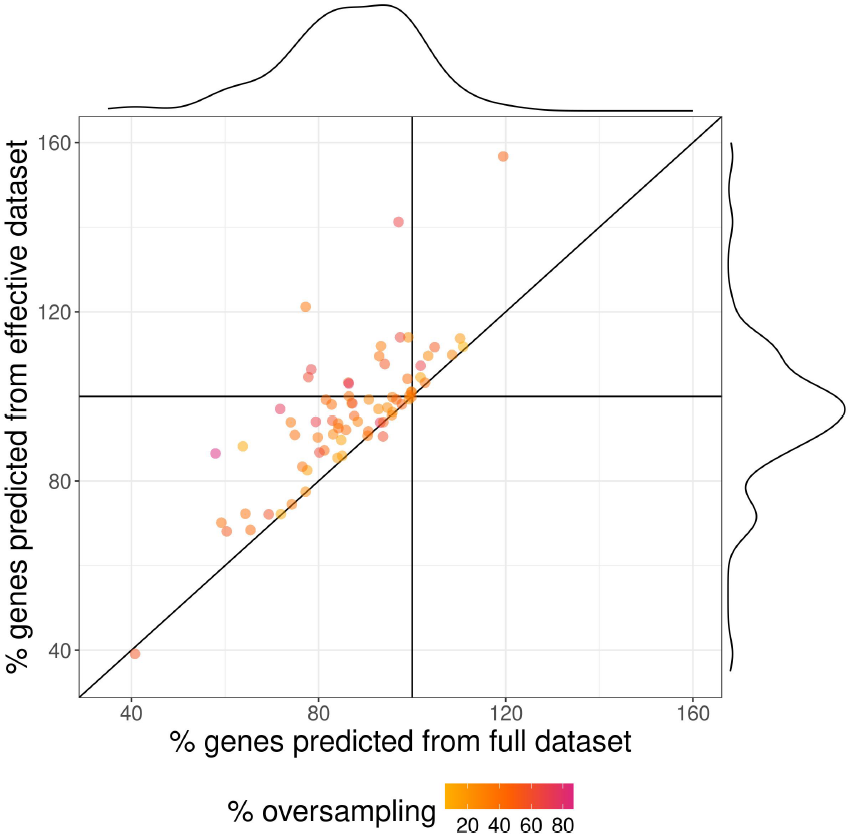
Retrospective 2015 predictions of 2023 pangenome sizes. The percentage of genes predicted by Heaps’ law (extrapolated from the full and effective datasets up to 2015) relative to the actual number of genes in the effective dataset are compared for 74 species. Sampling bias leads to an underestimate (i.e., *<* 100% on the horizontal axis) of effective pangenome sizes (mean 87.0% vs. mean 96.2%, Wilcoxon signed-rank test with continuity correction: p-value = 7.189*e −* 13) with larger difference observed in species with high percentage of oversampling (Kendall’s rank correlation tau = *−*0.2213573, p-value = 0.005331). Point color indicates the percentage of oversampling.

### Computing the average nucleotide identity

ANI-values were computed using *fastANI* ^65^. Due to computation times, *skani* ^66^ was used for some comparisons in the five largest genome sets: *E. coli, K. pneumoniae, S. enterica, S. aureus*, and *P. aeruginosa*.

### Computations from gene presence-absence matrix

The gene presence–absence matrix was generated using the corresponding helper script included in *panX* ^63^.

#### Gene frequency spectrum

The site frequency spectrum (SFS) is the number of mutated sites that occur in any *k* out of *n* samples. Similarly, the gene frequency spectrum (GFS) is the number of genes in *k* out of *n* samples ^67^. We computed the GFS directly from the gene presence–absence matrix.

#### Pangenome accumulation curve

A pangenome curve uses different permutations of the samples to estimate a growing function of the total number of genes in *k* samples ^68^. Following equation (3) in Parmigiani et al. ^69^, the expected number of distinct genes observed after sampling *k* genomes was computed as 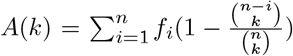, where *f*_*i*_ denotes the number of genes that occur in exactly *i* genomes (the gene frequency spectrum).

Pangenome accumulation curves approximately follow Heaps’ law, *g* = *cG*^*−α*^, where *g* is the number of newly found genes in sample number *G*, and *c* and *α* are constants estimated from the data ^70^. The parameters *c* and *α* were obtained from a linear regression of the Heaps’ law on a log-log scale. Predictions of the number of newly discovered genes upon sampling additional genomes were generated by extrapolating the fitted Heaps’ law.

#### Genomic fluidity

Genomic fluidity is defined as the ratio of unique genes to the total number of genes across randomly drawn pairs within *n* genome samples ^71^. Following equation (1) in Kislyuk et al. ^71^, fluidity was computed as 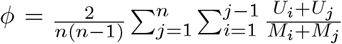, where *U*_*i*_ and *U*_*j*_ denote the number of genes unique to genomes *i* and *j*, respectively, and *M*_*i*_ and *M*_*j*_ represent their total gene counts.

### Measuring oversampling-concentration

The type of sampling bias ranges as a continuum from concentrated to distributed sampling bias (Fig. 4 A).

#### Concentrated sampling bias

A common type of intentional sampling bias is to excessively sample a single sub-population of interest. Sometimes a small number of very diverse genomes from other sub-populations are added in order to get a broad overview of the genetic diversity in the species ^72^. In its extreme form, the inferred phylogenetic tree contains only one group of highly related samples.

#### Distributed sampling bias

On the other hand, combining multiple various studies with (possible on purpose) concentrated sampling bias or unintentional sampling bias may lead to distributed sampling bias. In this case, the inferred phylogenetic tree contains several distinct groups of highly related samples distributed across the whole tree.

We suggest the following statistic *c*_*sb*_ as a possible measure on how concentrated the sampling bias in a given sampling-tree is:

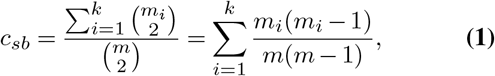

where *k ≥* 1 is the number of distinct oversampled groups, *m*_*i*_ the number of oversampled samples in group *i ∈ {*1, …, *k}* and 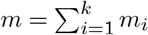 the number of all oversampled samples. Note that from each group of oversampled genomes we will maintain at least one (random) group member within the effective dataset, which does not contribute to *m*_*i*_. The oversampling-concentration *c*_*sb*_ gives the probability that two oversampled individuals belong to the same group of oversampled individuals. The statistic *c*_*sb*_∈ [0, 1], where *c*_*sb*_ = 1 corresponds to super concentrated sampling bias, i.e., the sampling-tree contains only one group of highly related samples. On the other hand, sampling bias can be distributed across the whole sampling-tree. In its extreme form the sampling-tree contains many groups of only two very closely related samples, and *m*_*i*_ = 1 for all *i*. In this case, *c*_*sb*_ = 0.

### Simulations and performance evaluation

*PhyloThin* was optimized and tested using simulation results generated under concentrated (*c*_*sb*_ = 1) and distributed (*c*_*sb*_ = 0) samplingbias scenarios, implemented in *R* ^53^. Details of the simulation setting are provided in Sup. Note 2. Simulations of partial selective sweeps were conducted using *msprime* ^73^ (Fig. 14b). Across sample sizes from 10 to 2, 000 genomes, we assessed the identification of oversampled and unbiased samples. With true coalescent branch lengths, *PhyloThin* detected oversampling under both oversampling concentrations. Adding Poisson-distributed mutations to branches showed that limited mutation resolution can lead to excessive removal, especially in large samples, unless the mutation-rate-sensitive thinning and subsampling adjustments are used. For further details on simulations, see Sup. Note 3.

**Fig. 14.**
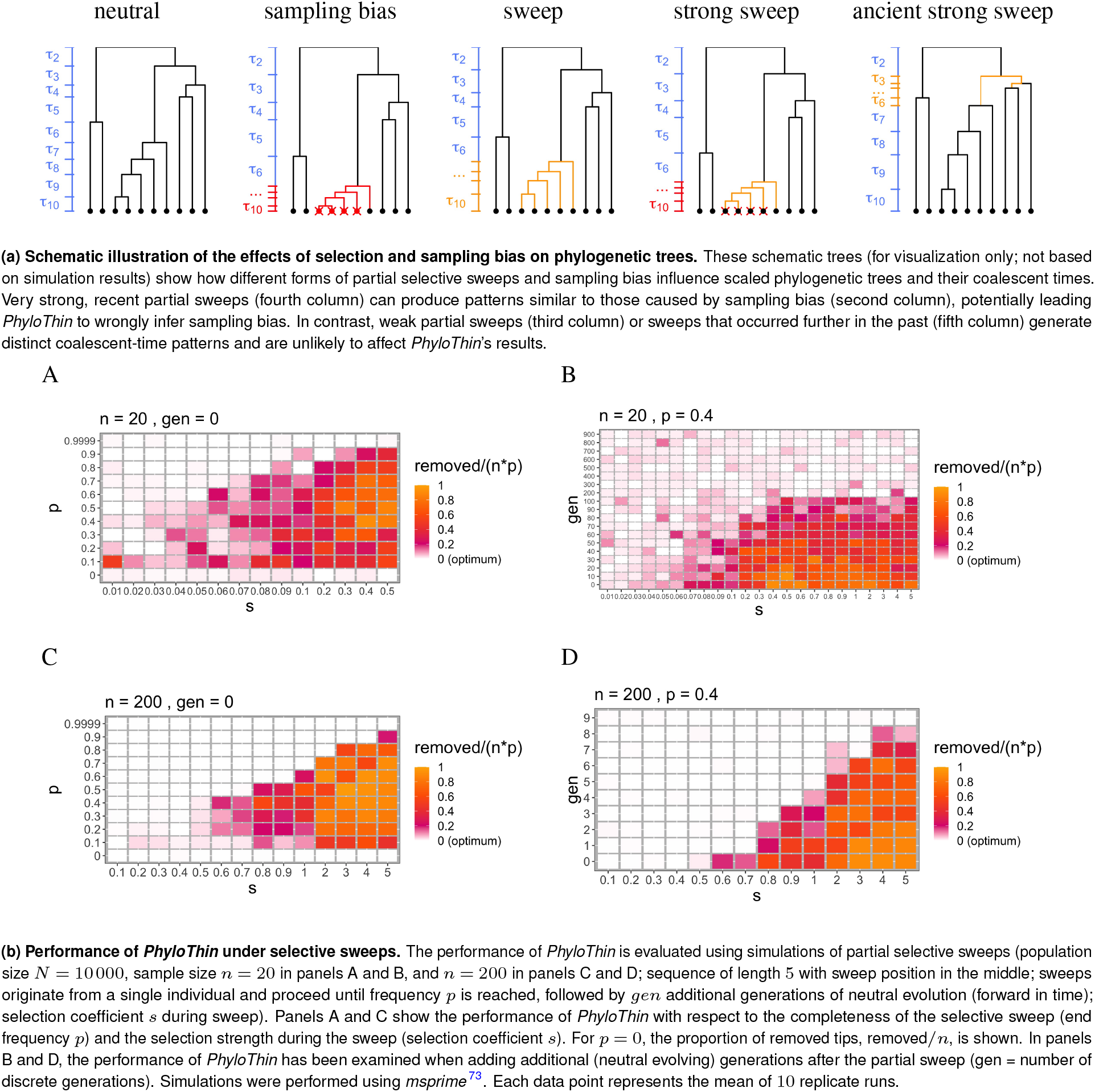
Patterns of selective sweeps and sampling bias. Selective sweeps do not affect the outcome of *PhyloThin* unless they are genome-wide, very recent, and extremely strong. For intermediate-frequency partial sweeps (end frequency *p* = 0.4), selection strengths above 0.6 where the sweep is ongoing or stopped less than 10 generations ago are necessary to confuse *PhyloThin* for samples of size 200 (Fig. 14b D). For smaller sample sizes (*n* = 20) a selection strength between 0.07 and 0.2 within the last 50 generations and above 0.2 within the last 100 generations confused *PhyloThin* in simulations (Fig. 14b B). For low-frequency and nearly complete sweeps,*PhyloThin* is more accurate (Fig. 14b A and C). Population expansions can produce patterns similar to those of genome-wide selective sweeps in nonrecombining species ^55^, and thus we expect *PhyloThin*’s results to be similarly robust against phylogenetic patterns of population growth.

### Statistics

All statistical analyses were performed in *R* ^53^ using nonparametric tests. Differences between paired datasets (for example, comparisons of full versus effective datasets to quantify sampling-bias effects) were assessed using the two-sided Wilcoxon signed-rank test with continuity correction and analogously the Wilcoxon rank-sum test for unpaired datasets (e.g., pathogenic and non-pathogenic species). Associations between differences in statistics (induced by sampling bias) and other variables (such as the percentage of oversampling) were quantified using Kendall’s rank correlation. All p-values and Kendall’s tau values are reported in the corresponding figure legends.

### Use of AI tools

Large language models were used to assist with language editing and code debugging. All AI-assisted outputs were critically reviewed and verified by the authors, who take full responsibility for the final manuscript and code.

### Data availability

The genomic data supporting the findings of this study are available from the *National Center for Biotechnology Information* (NCBI) of the *National Library of Medicine*. All processed data generated during the study are provided within the paper and its Supplementary Information, or are available from the authors upon request.

### Code availability

Unless stated otherwise, simulations, computations, statistical analyses, and data visualization were performed using a combination of *R* ^53^ and *Bash* scripts. The source code for *PhyloThin* is openly available on GitHub at 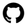 github.com/fbaumdicker/phylothin All further scripts used for the analyses in this study are available from the authors upon request.

## ACKNOWLEDGEMENTS

We thank Conrad Roth for providing us with precomputed ANI values ^21^ for a subset of our data and Xiaoqian Annie Yu for providing their phylogenetic tree-data for genome-wide sweeps ^31^. We thank Daria Evseeva, Juan Bosco Gracia Alvira, Eduardo Rocha, and Detlef Weigel for discussions.

## Additional information

Supplementary information is available for this paper:

- Sup. Note 1: *PhyloThin* - Theoretical Derivation and Implementation
- Sup. Note 2: Simulation settings
- Sup. Note 3: Simulations
- Sup. Note 4: NCBI data
- Sup. Note 5: Supplementary figures
- Supplementary Figures - NCBI species
- Supplementary Table - NCBI_species.csv

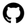 github.com/fbaumdicker/phylothin Correspondence and requests for materials should be addressed to F.B.

## Extended data

