## Supplementary Notes 1 to 5 for "Hidden but Expanding Sampling Bias Distorts Prokaryotic Genome Collections"

#### Supplementary Note 1: *PhyloThin* - Theoretical Derivation and Implementation

*PhyloThin* is based on the coalescent intervals of an ultrametric tree. Backward in time, the coalescent interval $t_i$ ($i = 1,...,m$, where $m$ denotes the number of nodes of the tree) is the time between the $i$-th and $(i+1)$-th coalescent event. From now on, we assume $n = m+1$, i.e., no multiple and simultaneous mergers (they may be considered as coalescent intervals of length zero). Before testing the dataset for oversampled genomes, the ultrametric phylogenetic tree needs to be scaled accordingly. Fig. 1 shows the concept of *PhyloThin* on different example-trees.

**Theoretical Background.** Coalescent times of a sample of size $n$ under the Kingman coalescent model are independent of the topology of the tree. Moreover, under the coalescent time scale (scaled by the population size $N$ and limit $N \to \infty$) the times $T_i$ - during which there were exactly $i$ lineages ancestral to the sample - follow an exponential distribution,i.e., $T_i \sim \text{Exp}\left(\frac{i(i-1)}{2}\right)$ [74]. Therefore, $\tau_i := \binom{i}{2} T_i \sim \text{Exp}(1)$ iid for all $i = 2,...,n$. Hence, coalescence times (under neutrality) are after this transformation identically distributed and independent of parameters. Now we can use these scaled and transformed random variables to identify unusually short times, which result from sampling bias. However a single time is exponentially distributed, so the test would not be sensitive enough to identify too short coalescent times. Thus we consider the $l$-th cumulative sum of the scaled coalescent time of the $l+1$ most closely related individuals (most closely related in this context could also mean two or more groups of closely related individuals),

$$c_l = \sum_{i=n-l}^{n} \tau_i, \tag{2}$$

which follows a gamma distribution with shape parameter $l+1$ and rate 1 for all $l = 0,...,n-2$. This is the crucial result *PhyloThin* is using to detect oversampled genomes.

***Scale the ultrametric phylogenetic tree.*** Usually phylogenetic trees and thus their coalescent intervals are given in units of their mutation rate. In order to use the results from coalescent theory given above, in a first step, the coalescent intervals $t_i$, $i = 1,...,m$, need to be scaled to coalescent time scale. Under neutrality and without any sampling bias, $t_i$ follows approximately an exponential distribution with parameter $\binom{m-i+2}{2}/s$, where $s$ is the scaling parameter we are interested in. For absolute coalescent times and in the limit $N \to \infty$, the scaling parameter $s$ can be seen as an approximation for the total population size $N$. Given an index set $I = \{i_1,...,i_\kappa\} \subseteq \{2,...,m+1\}$ of size $\kappa$, the scaling parameter $s$ can be estimated by taking the average over the corresponding $\kappa$ (transformed) coalescent intervals,

$$s = \frac{\sum_{i \in I} \binom{i}{2} t_{m-i+2}}{\kappa}. \tag{3}$$

For $i = 2,...,m+1$, the coalescent times on coalescent time scale can then be approximated by

$$T_i = \frac{t_{m-i+2}}{s}. \tag{4}$$

In a second step, a transformation of the scaled coalescent times $T_i$ ensures that the coalescent times are identically distributed. In this step, each coalescent time $T_i$ is multiplied by $\binom{i}{2}$, i.e.,

$$\tau_i = \binom{i}{2} T_i \tag{5}$$

for $i = 2,...,m+1$ such that $\tau_i$ is exponentially distributed with mean 1 under the assumption of neutrality and no sampling bias.

***Remove closely related samples.*** As explained above, we examine sums of scaled and transformed coalescent times to identify oversampled individuals, i.e., to find too short external branches. The following pseudo-algorithm illustrates the iterative testing and removal of oversampled samples in *PhyloThin*.

**Require:** $l_{\max}$ threshold for testing, ▷ Eq. (13)
$\{\tau_{l_{\max}}, ..., \tau_n\}$ transformed coalescent times of the scaled ultrametric tree

$j \leftarrow n$
$\tilde{\tau}_j \leftarrow \tau_n$
$\mathcal{T} \leftarrow \{\tau_{l_{\max}}, ..., \tau_{j-1}, \tilde{\tau}_j\}$
$c_l \leftarrow \sum_{i=n-l}^{n} \tau_i$ for $l < n - l_{\max}$ ▷ Eq. (2)
**while** $j > l_{\max}$ **do**
  **if** one of the $c_l$ too short (for some $l < j - l_{\max}$, the cumulative probability $\mathrm{pgamma}(c_l) < \alpha$) **then** ▷ ($\star$)
    one shortest external branch gets removed from the tree and
    the coalescent times of the resulting tree are used from now on, i.e.,
    $\tilde{\tau}_{j-1} \leftarrow \tilde{\tau}_j + \tau_{j-1}$
    $\mathcal{T} \leftarrow \{\tau_{l_{\max}}, ..., \tau_{j-2}, \tilde{\tau}_{j-1}\}$
    $j \leftarrow j - 1$
    $c_l \leftarrow \tilde{\tau}_j + \sum_{i=j-l}^{j-1} \tau_i$ for $l < j - l_{\max}$
  **else**
    Stop
  **end if**
**end while**

**Implementation.** *PhyloThin* is written in *R*[75] and uses the *ape*[76], *phangorn*[77] and *dplyr*[78] libraries. Before *PhyloThin* can be used, a ultrametric phylogenetic tree needs to be constructed from the $n$ genome samples. This tree can be reconstructed by using, e.g., *panX*[79] and *TreeTime*[80] or *PATHd8*[81]. A schematic documentation on how *PhyloThin* works is shown in Fig. 18.
As described above, the algorithm of *PhyloThin* can be divided into two parts: the scaling/transformation step and the testing/removal step. Fig. 15 shows the full scaling step on a simulation example. The crucial component in the scaling step is to obtain the index set $I$ to estimate the scaling parameter $s$ which is needed to scale the tree to coalescent time scale.

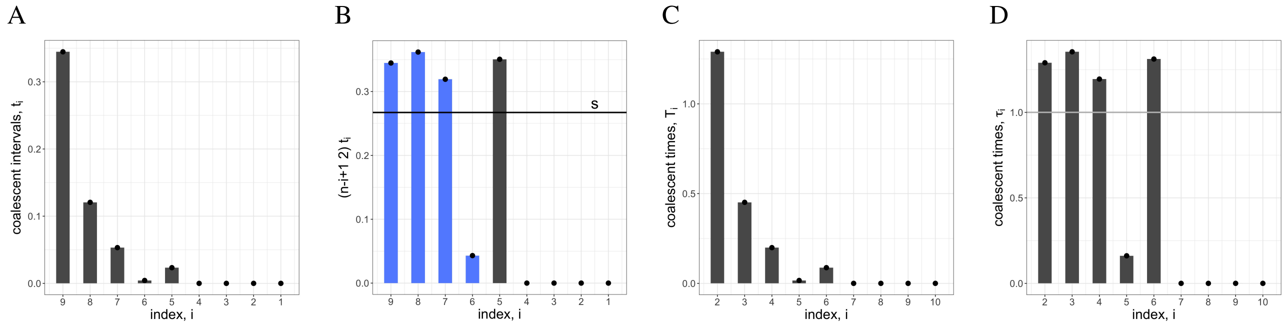


**Fig. 15.** In panel A, the coalescent intervals $t_i$ are shown for a simulation example for concentrated sampling bias with $n = 10$ samples from which a group of $np = 4$ samples is oversampled (same simulation example as in Fig. 18). In panel B, the coalescent intervals $\binom{n-i+1}{2} t_i$ are rescaled to find the scaling factor $s$, Eq. S (3). The blue bars show the coalescent intervals which have been used to obtain the scaling factor $s$ indicated by the horizontal black line. In panel C, the coalescent times $T_i$, Eq. S (4), are on coalescent time scale. In panel D, the rescaled coalescent times $\tau_i$, Eq. S (5), are shown. Under neutrality, $\tau_i$ is drawn from an exponential distribution with mean one.

***Obtain the index set*** $I$ ***to scale the tree.*** The scaling parameter $s$ can be estimated by averaging over (transformed) coalescent intervals, Eq. (3). However, if all coalescent intervals would be used, sampling bias would also influence the scaling. As long as only very few individuals are oversampled, this does not have a strong impact. But if for example $80\%$ (which is not unrealistic as we see in data, Sup. Note 4) of the individuals are oversampled, this would obscure the scaling. Hence, careful consideration must be given to selecting the index set $I$ in Eq. (3).
Under neutrality, the (transformed) coalescent intervals $\binom{i}{2} t_{m-i+2}$ follow approximately an exponential distribution with mean $s$. Deviations from this distribution can have various reasons such as selection, spatial population structure, demographic changes etc. Also sampling bias causes shorter coalescent intervals for larger $i$ (this is what we are looking for in the last step for removing samples). To avoid the signature of sampling bias in the estimation of the scaling parameter $s$, only coalescent intervals closer to the root are considered, i.e., $I \subseteq \{2, ..., k\}$. Therefore, we need a first rough estimation on how many $(k-1)$ coalescent intervals are not affected by sampling bias.

When comparing coalescent trees with and without sampling bias, the last coalescent times of the tree actually combine to the same one coalescent interval in the corresponding reduced coalescent tree without sampling bias. In other words, if $(k-1)$ coalescent intervals are not affected by sampling bias, the sum $\binom{k+1}{2}\sum_{j=k+1}^{n} t_{m-j+2}$ corresponds to the first (transformed) coalescent interval in the reduced neutral coalescent tree without sampling bias and therefore follows also approximately an exponential distribution with mean $s$. Keeping this in mind, we can construct random variables with the same mean $s$. Because $s$ is the scaling parameter that is still unknown (the main goal of the scaling step is to estimate $s$), we eliminate it by clever division. Therefore, we define

$$f_i = \frac{\sum_{j=2}^{i}\binom{j}{2}t_{m-j+2}}{i-1} \Bigg/ \frac{\sum_{j=2}^{i}\binom{j}{2}t_{m-j+2} + \binom{i+1}{2}\sum_{j=i+1}^{n} t_{m-j+2}}{i} \tag{6}$$

$$= \frac{\sum_{j=2}^{i}\tau_j}{\sum_{j=2}^{i}\tau_j + \binom{i+1}{2}\sum_{j=i+1}^{n} T_j} \cdot \frac{i}{i-1} \tag{7}$$

for $i = 2,...,m$. Importantly, $f_i$ is independent of the parameter $s$. Under neutrality, $\tau_j \sim \mathrm{Exp}(1)$. If there are $n-k-1$ genomes oversampled (and otherwise neutrality can be assumed), then $\binom{i+1}{2}\sum_{j=i+1}^{n} T_j \sim \mathrm{Exp}(1)$ for $i = k$. Therefore,

$$\mathrm{E}[f_i] = \begin{cases} < 1 & \text{for } i < k, \\ 1 & \text{for } i = k. \end{cases} \tag{8}$$

Since we assume that shorter external branches are caused by sampling bias, we define

$$k = \max\{i : f_i < \alpha_3\} \tag{9}$$

for a threshold $0 < \alpha_3 \leq 1$. In addition, we consider only coalescent intervals greater than zero and the following restrictions are set: $k \geq \min\{6, 0.1\cdot n+1\}$ and $\kappa \geq \min\{5, 0.1\cdot n\}$. Fig. 16 A shows the statistic $f_i$ and the selection of $k$ on a simulation example.

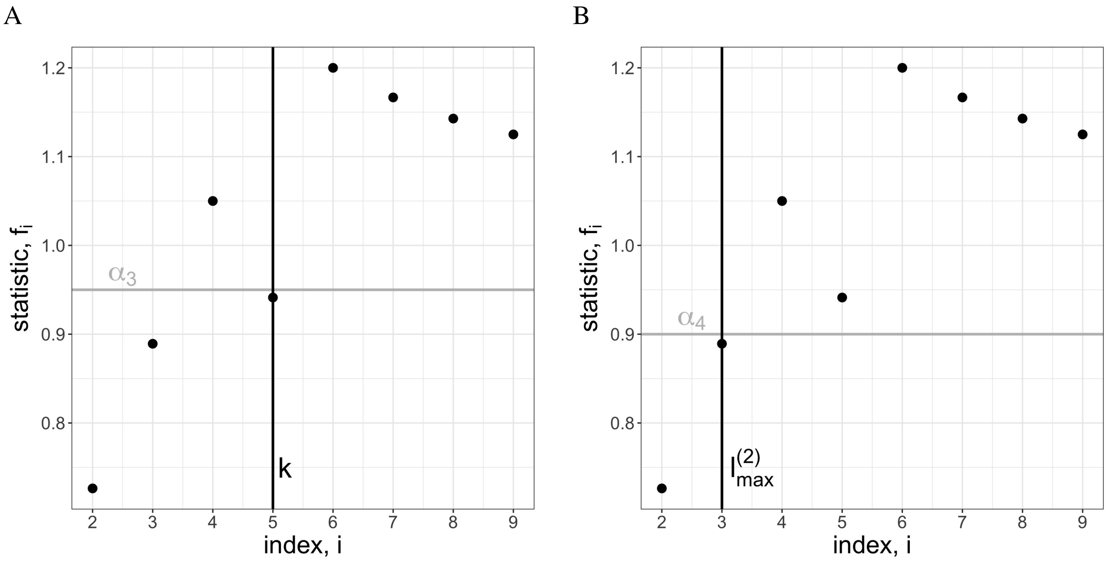


**Fig. 16.** The statistic $f_i$, Eq. S (7), is used to determine $k$ (panel A, Eq. S (9)) and $l^{(2)}_{\text{max}}$ (panel B, Eq. S (12)). Here, $f_i$ is shown for a simulation example for concentrated sampling bias with $n = 10$ samples from which a group of $np = 4$ samples is oversampled (same simulation example as in Fig. 18).

***Remove closely related samples.*** Analogous to the cumulative sums $c_l$, Eq. (2), under neutrality, the cumulative sums

$$\tilde{c}_{n-l} = \sum_{i=2}^{n-l} \tau_i \tag{10}$$

follow approximately a gamma distribution with shape parameter $n-l-1$ and rate 1 for all $l = 0,...,n-2$. The cumulative sums $\tilde{c}_{n-l}$ starting from the root (Fig. 17 A) give a good first idea on how many samples should be removed. Since the cumulative sums $c_l$ starting from the tips (Fig. 17 B) are not biased from earlier demographic and/or selection events, they are then used for a refined testing on how many tips actually get removed.

First, a bound for testing is defined by $l_{\text{max}}$. However, the estimate $l^{(1)}_{\text{min}}$ based on $\tilde{c}_{n-l}$ (Fig. 17 C) could suffer from too many short coalescent intervals near the root and in the middle of the tree (possibly due to a low mutation rate or non neutral processes) and would suggest a too low bound in that case. Thus, an additional restriction on the bound $l_{\text{max}}$ for testing is defined through the statistic $f_i$, Eq. (7):

$$l^{(1)}_{\text{min}} = \min\{0 \le l \le n-2 : \text{pgamma}(\tilde{c}_{n-l}) > \alpha_0\}, \quad \textbf{(11)}$$

$$l^{(2)}_{\text{max}} = \max\{i : f_i < \alpha_4\}, \quad \textbf{(12)}$$

$$l_{\text{max}} = \begin{cases} l^{(2)}_{\text{max}} & \text{if } l^{(1)}_{\text{min}} = 0 \\ \max\{n - l^{(1)}_{\text{min}}, l^{(2)}_{\text{max}}\} & \text{else,} \end{cases} \quad \textbf{(13)}$$

where $0 < \alpha_0 < 1$, $0 < \alpha_4 \le 1$ and pgamma gives the cumulative probability (Fig. 17 A).

($\star$) The sample with the shortest external branch gets removed as long as for the remaining coalescent tree

- $\tau_n$ is too short for the exponential distribution, i.e., $\text{pgamma}(c_0) < \alpha_1$, or
- the cumulative sums are too short for the gamma distribution, i.e., there exists a $l < n - l_{\text{max}}$ such that $\text{pgamma}(c_l) < \alpha_2$,

where $0 < \alpha_1 \le 0.5$ and $0 \le \alpha_2 < \alpha_1$ are the significance levels (Fig. 17 D). Here, we consider the condition on which sample among the shortest external branches (usually two) gets removed. For example, the youngest sample among the shortest tips has been removed for Fig. 2 and Fig. 6. If no preference is given, one branch among the shortest external branches is removed at random (only stochastic component of *PhyloThin*).

**Tuning parameters.** The following parameters have been optimized with the help of computer simulations (for more details, see Sup. Note 3):

- $\alpha_0 = 0.5$ (Fig. 23):
  significance level to find bound for testing, Eq. (11),
- $\alpha_1 = 0.1$ (figures 24 to 30):
  significance level for removing
  (examines if the first scaled and transformed coalescent interval is too short for the exponential distribution),
- $\alpha_2 = 0.025$ (figures 24 to 30):
  significance level for removing
  (examines if the sums of scaled and transformed coalescent intervals are too short for the gamma distribution),
- $\alpha_3 = 0.95$ (Fig. 20):
  threshold to find bound for scaling, Eq. (9),
- $\alpha_4 = 0.9$ (Fig. 20):
  threshold to find bound for testing, Eq. (12).

**Mutation-rate-sensitive thinning.** For low mutation rates, branches of length zero become more and more frequent (especially in large phylogenetic trees) which may violate the assumptions of the default settings in *PhyloThin* and may lead to an overestimation of sampling bias. We therefore recommend ensuring that a sufficient number of variable sites were used to infer the phylogenetic tree. If this is not possible, we suggest cross-checking the results obtained with *PhyloThin* against those of the mutation-rate-sensitive version of *PhyloThin*.

When placing mutations on an ultrametric tree, each branch length $\tilde{b}$ in the phylogenetic tree is drawn from a Poisson distribution with mean $\Theta \cdot b$, where $\Theta$ denotes the mutation rate and $b$ the corresponding branch length in the ultrametric tree, i.e., $\tilde{b} \sim \text{Poi}(\Theta \cdot b)$. Assuming no further effects, the mutation-aware coalescent times $\tilde{T}_i \sim \text{Poi}(\Theta \cdot T_i)$ on coalescent time scale, where $T_i \sim \text{Exp}\left(\frac{i(i-1)}{2}\right)$. Hence, $\tilde{T}_i$ follows a geometric distribution with mean $\frac{2\Theta}{i(i-1)}$, i.e., $\tilde{T}_i \sim \text{Geom}\left(\frac{i(i-1)}{i(i-1)+2\Theta}\right)$. The random variables $\tilde{T}_i$ can no longer be transformed to be i.i.d., but the distribution of their sum can still be computed efficiently (at least numerically and with the help of Fourier transforms).

In the mutation-rate-sensitive version of *PhyloThin*, an inferred mutation rate $\Theta = m/\sum L_i$ is incorporated into the statistical test ($\star$), where $m$ denotes the number of variable sites (SNP positions) used to infer the phylogenetic tree and $\sum L_i$ is the total branch length of the ultrametric tree on coalescent time scale. The workflow of *PhyloThin* then differs only in the distributions used in the statistical test during the removing step ($\star$). Instead of the exponential distribution ($\text{pgamma}(c_0) < \alpha_1$), the last

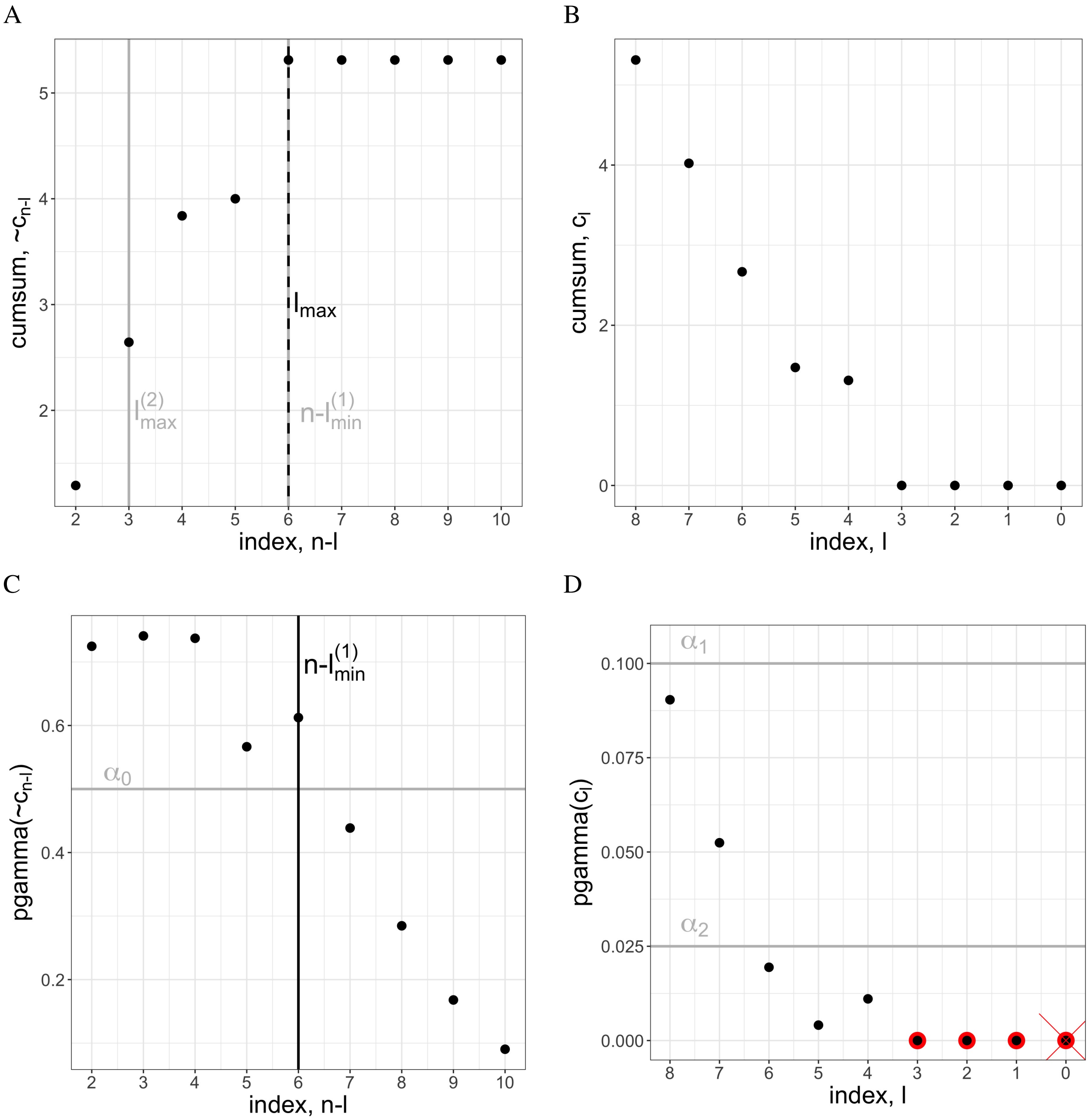


**Fig. 17.** The cumulative sums $\tilde{c}_{n-l}$ starting from the root (panel A, Eq. S (10)) are used to define $l^{(1)}_{\text{min}}$ (panel C, Eq. S (11)). The cumulative sums $c_l$ starting from the tips (panel B, Eq. S (2)) are used for testing for sampling bias. In panel D, the examined cumulative sums $c_l$ are indicated by red edgings. Because the threshold values $\alpha_1$ and $\alpha_2$ are undercut by pgamma$(c_l)$, our statistical test detects sampling bias. One of the shortest external branches gets removed, indicated by the red cross. The statistics pgamma$(c_l)$ is then recalculated for the reduced tree and the testing is repeated.
All quantities are shown for a simulation example for concentrated sampling bias with $n = 10$ samples from which a group of $np = 4$ samples is oversampled (same simulation example as in Fig. 18).

coalescent interval is tested against the geometric distribution with mean $\frac{2\Theta}{n(n-1)}$ $(\text{pgeom}(T_n) < \alpha_1)$, and instead of the gamma distribution $(\text{pgamma}(c_l) < \alpha_2$ for $l < n - l_{\text{max}})$, the cumulative sums are tested against the corresponding convolutions of geometric distributions $(\text{pgeomconv}(\sum_{i=n-l}^{n} T_i) < \alpha_2$ for $l < n - l_{\text{max}})$.
*Mutation-rate-sensitive Thinning* may underestimate sampling bias if the resolution of the phylogenetic tree is insufficient. This is the case if the expected number of differences for many recent coalescent times is actually zero when taking the mutation rate into account. At some point the number of segregating sites is simply to low such that sampling bias gets undetectable as we no longer have the resolution to estimate the correct distance between closely related individuals. The smallest cluster of highly related samples of distance zero, which can be detected, contains at least

$$k(n) = \min\{2 \le k \le n-1 : \prod_{i=n-k+2}^{n} \frac{i(i-1)}{i(i-1)+2\Theta} < \alpha_2\} + 1 \tag{14}$$

samples.

*PhyloThin* outputs a warning when the resolution might be too low ($k(n) > 0.1 \cdot n$). To mitigate this effect in this case, we suggest to increase the resolution of the phylogenetic tree by incorporating more genes in the tree-inference (e.g., soft core genome) or - if not possible/reasonable - combining *Mutation-rate-sensitive Thinning* and *Subsampling*, which increases the chance to identify at least the concentrated oversampling clusters.

**Subsampling.** As the number of samples $n$ increases, the inference of the shortest coalescent times becomes less accurate. To avoid underestimating sampling bias, we suggest repeatedly subsampling $x$ times the coalescent tree reduced by *PhyloThin* with the additional provided *R*-script. *PhyloThin* is then applied for each subsample of size $n_{\text{sub}}$. The default parameter setting is calculated as follows: Under neutrality and on coalescent time scale, the shortest external branch $\frac{b_{\text{min}}}{s}$ is expected to have length $\frac{n(n-1)}{2}$. In order for sampling bias to be detectable, $0 < \frac{b_{\text{min}}}{s} < \frac{n(n-1)}{2}$. Assuming a perfect neutral coalescent tree, $s \approx \frac{L}{2}$, where $L$ denotes the total tree length and using that on coalescent time scale the total tree length is 2 for $n \to \infty$. We approximate $\frac{2 \cdot b_{\text{min}}}{L} \lesssim \frac{n^2}{2}$. Hence, we choose

$$n_{\text{sub}} = \left\lceil \sqrt{\frac{L}{b_{\text{min}}}} \right\rceil, \tag{15}$$

where $b_{\text{min}} > 0$ gives the length of the shortest external branch (larger than zero) in the coalescent tree and $L$ the total tree length. By default, subsampling is repeated often enough that each tip is sampled 100 times on average,

$$x = \left\lceil \frac{100 \cdot n}{n_{\text{sub}}} \right\rceil. \tag{16}$$

The significance level $\alpha_5 = 0.9$ indicates the proportion on how often a sample has to be in an oversampling class so that it is classified as oversampled in the entire tree. This means that in order to detect an oversampled genome, the probability that another sample of the same cluster (of size $c$) is in the subsample has to be larger than the threshold $\alpha_5$:

$$1 - \prod_{i=1}^{n_{\text{sub}}-1} \left(1 - \frac{c-1}{n-i}\right) > \alpha_5. \tag{17}$$

In other words, only clusters that contain at least $c$ samples can be reliably detected by the subsampling method.
For *Mutation-rate-sensitive Thinning*, the default subsample size $n_{\text{sub}}$ is computed in a different way. This choice reflects a trade-off: a small value of $n_{\text{sub}}$ improves the detection of clusters for low mutation rates as short external branches can be more reliable classified as sampling bias, while a large value of $n_{\text{sub}}$ is needed to detect small clusters, since only then are enough of their samples included in the subsample. We therefore choose the default subsampling size

$$n_{\text{sub}} = \max\{2 < n_{\text{sub}} \leq n : c(n_{\text{sub}}) \geq k(n_{\text{sub}})\}, \tag{18}$$

with the monotone increasing function $k(n_{\text{sub}})$ defined in Eq. (14) and the monotone decreasing function $c(n_{\text{sub}})$ defined by rearranging Eq. (17).
Given that one sample of a cluster of size $c_k$ is contained in a subsample of size $n_{\text{sub}}$, the probability of finding $k-1$ additional samples of the same cluster in that subsample is

$$1 - \sum_{i=0}^{k-2} \frac{\binom{c_k-1}{i}\binom{n-c_k}{n_{\text{sub}}-1-i}}{\binom{n-1}{n_{\text{sub}}-1}} \tag{19}$$

(hypergeometric distribution).

#### References.


74. Kingman, J. F. C. The coalescent. *Stochastic Processes and their Applications*, 13(3):235–248, 1982. doi: 10.1016/0304-4149(82)90011-4.
75. R Core Team. *R: A Language and Environment for Statistical Computing.* R Foundation for Statistical Computing, Vienna, Austria, 2021.
76. Paradis, E and Schliep, K. ape 5.0: an environment for modern phylogenetics and evolutionary analyses in R. *Bioinformatics*, 35:526–528, 2019. doi: 10.1093/bioinformatics/bty633.
77. Schliep, K. phangorn: phylogenetic analysis in R. *Bioinformatics*, 27(4):592–593, 2011. doi: 10.1093/bioinformatics/btq706.
78. Wickham, H et al. *dplyr: A Grammar of Data Manipulation*, 2023. R package version 1.1.2.
79. Ding, W, Baumdicker, F and Neher, R. A. panX: Pan-genome analysis and exploration. *Nucleic Acids Research*, 46(1):e5–e5, 2018. doi: 10.1093/nar/gkx977.
80. Sagulenko, P, Puller, V and Neher, R. A. TreeTime: Maximum-likelihood phylodynamic analysis. *Virus Evolution*, 4(1):vex042, 2018. doi: 10.1093/ve/vex042.
81. Britton, T et al. Estimating divergence times in large phylogenetic trees. *Systematic Biology*, 56(5):741–752, 2007. doi: 10.1080/10635150701613783.

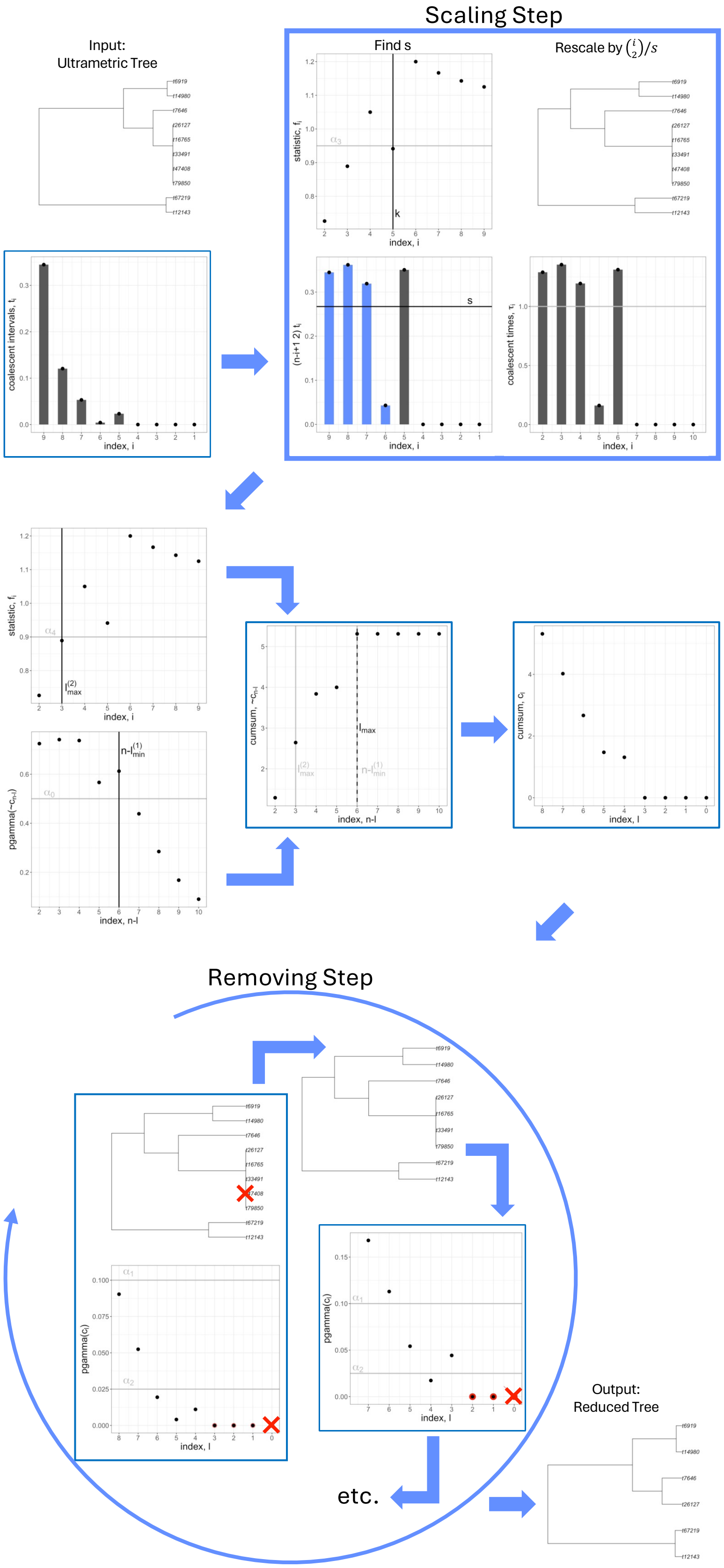


**Fig. 18. How does *PhyloThin* work?** A schematic overview on the mechanism of *PhyloThin* is shown on a simulation example for concentrated sampling bias with $n = 10$ samples from which a group of $np = 4$ samples is oversampled (for the simulation setting see Sup. Note 2). First the given ultrametric tree is scaled, then after a preprocessing step, the oversampled tips are removed iteratively and the reduced tree is output (for more details on the mechanism of *PhyloThin* see Sup. Note 1 and figures 15 to 17).

#### Supplementary Note 2: Simulation settings

We use two different strategies for simulating a ultametric tree of size $n$ with a fraction of $p$ oversampled tips (Fig. 19): Simulations with concentrated sampling bias and simulations with distributed sampling bias. In both cases, first a coalescent tree of the full population is simulated (with $N = 100\,000$ for concentrated and $N = 10\,000$ for distributed sampling bias). For optimizing the tuning parameters (in Sup. Note 3), we used simulations with concentrated sampling bias.
**Simulation parameters**: total population size $N \in \mathbb{N}$, sample size $n \in \mathbb{N}$ ($n \ll N$), fraction of sampling bias $p \in [0,1)$.

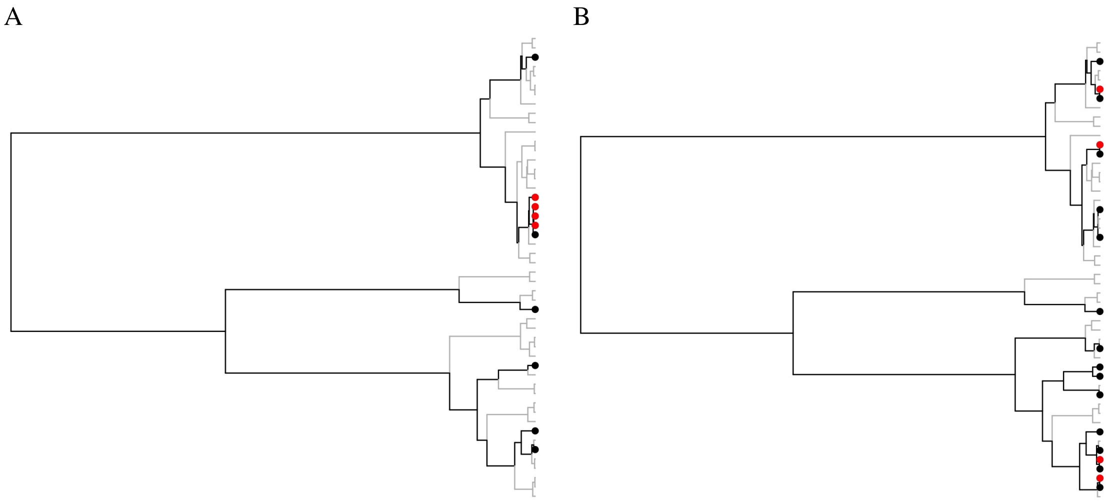


**Fig. 19.** Examples of concentrated sampling bias (panel A) and of distributed sampling bias (panel B) according to Sup. Note 2 are shown for $N = 50$, $n = 10$ and $p = 0.4$. The small population size $N = 50$ has been chosen for visualization purposes (usually $N = 100\,000$ and $N = 10\,000$ are used). The full (neutral) coalescent tree with $N = 50$ tips is shown in gray color in the background. The sampled subtree with $n = 10$ tips is shown in black color. The oversampled tips are indicated by red dots.

**Simulations with concentrated sampling bias.** The sampling-tree contains only one group of highly related samples. The subtree consisting of the $np+1$ most related tips from the total population tree represents one random sample which is highly oversampled. Then $n(1-p)-1$ samples are taken randomly from the remaining tips.
The probability that a randomly sampled tip lays in the oversampled group is

$$\text{Prob}_C[\text{oversampled}] = \frac{\text{np}+1}{\text{N}}. \quad \textbf{(20)}$$

Hence, sampling bias is only present/detectable if this probability is not too large. Another restriction on the detectability of sampling bias relates to the depth of the coalescent (sub)tree consisting of the oversampled group.

**Simulations with distributed sampling bias.** The sampling-tree contains highly related samples distributed across the whole tree. We restrict the parameter space to $p \leq 0.5$, since $p > 0.5$ leads to groups of more than one oversampled tip which makes the scenario getting closer to concentrated sampling bias. Tips with the smallest pairwise distance form the group of the oversampled samples and their highly related samples. Thus, this set of $np+g$ tips consists of $g$ groups of $\geq 2$ tips. For a large enough $N$, there are $np$ groups of two tips. At the end, the remaining $n(1-p)-g$ samples are taken randomly from the remaining tips.
The probability that a randomly sampled tip lays in a oversampled group is

$$\text{Prob}_D[\text{oversampled}] = \frac{\text{np}+\text{g}}{\text{N}}. \quad \textbf{(21)}$$

Therefore, the probability that a randomly sampled tip lays in a oversampled group is higher for distributed sampling bias compared to the respective concentrated sampling bias. However the depths of the coalescent (sub)trees of the oversampled group (in most cases two tips in each group) is much shallower for distributed sampling bias than for concentrated sampling bias. Thus, distributed sampling bias may be easier to detect with the help of *PhyloThin*.

**Adding mutations.** In order to make our simulation results closer to real data, a branch of length $b$ of the simulated coalescent tree is rescaled by the mutation rate $\Theta$ as follows: The new branch length is a realisation of a Poisson-distributed random variable with mean $\Theta \cdot b$. This method assumes that the number of mutation events on each branch follow a Poisson distribution and are visible at the time point of sampling. As the resulting tree is no longer ultrametric, we use *PATHd8*[81] before applying *PhyloThin*.

#### Supplementary Note 3: Simulations

We used computer simulations to optimize the tuning parameters and to test the performance of *PhyloThin*. For simulating a ultrametric, phylogenetic tree obtained from a biased sampling scheme, we use two different strategies for the two extreme cases of sampling bias: concentrated and distributed sampling bias. Both simulation settings are described and discussed in Sup. Note 2.

**Optimizing parameters.** *PhyloThin* consists of mainly two steps (see also Sup. Note 1): the scaling-step, where the tree gets scaled to coalescent time scale (the main step here is to obtain the index set $I$), and the removing-step, where oversampled genomes are identified. The performance of *PhyloThin* with optimized tuning parameters is shown on simulation results in Fig. 33.

***Optimizing the parameters for the removing-step.*** Simulations with concentrated sampling bias are used to optimize the tuning parameters $\alpha_0$, $\alpha_1$ and $\alpha_2$ in the removing-step. No mutation events are added,i.e., we use the true tree lengths instead of mutations (no need of $\alpha_4$). We use $I = \{2, \cdots, n(1-p)\}$ (no need of $\alpha_3$) and vary first $\alpha_0 \in [0.3, 0.7]$ (Fig. 23), and then $\alpha_1 \in (0, 0.2]$ and $\alpha_2 \in [0, \alpha_1)$ (figures 24 to 30).
In Fig. 23 a scaled version of $l^{(1)}_{\text{min}}$, Eq. (11), is examined for different sample sizes $n$ and percentages of sampling bias $p$. Since $l^{(1)}_{\text{min}}$ is a conservative estimate for the number of oversampled genomes $np$, $\alpha_0 = 0.5$ seems to be a good choice. This also makes sense theoretically as $\alpha_0$ gives a threshold on the cumulative distribution function on coalescent intervals which should follow a gamma distribution.
In figures 24 to 30 the number of removed tips, the number of false negatives and false positives are examined for different sample sizes $n$ and percentages of sampling bias $p$. In order to remove the right number of tips while keeping false positives and false negatives low, $\alpha_1 = 0.1$ and $\alpha_2 = 0.025$ seem to be a good choice.

***Optimizing the parameters for the scaling-step.*** For neutral coalescent trees with $n \to \infty$ tips, the statistic $f_{n-1}$, Eq. (7), is expected to be one. Fig. 20 shows that $f_{n-1} > 0.95$ with probability $0.9$ for $n \geq 30$. Thus, we choose $\alpha_3 = 0.95$. The parameter $\alpha_4$ (which is used in the scaling-step, see Eq. (12)) has a similar role as $\alpha_3$. However, $\alpha_4$ gives a bound for the number of coalescent intervals near the tips, whereas $\alpha_3$ gives a bound for the intervals near the root. Since the estimates should be conservative in order to account for stochastic effects (and deviations) in the data, $\alpha_4 < \alpha_3$ should hold. Hence, we set $\alpha_4 = 0.9$ which matches the $5\%$ quantile for sample sizes $n \geq 20$ in Fig. 20.

**Performance-analysis.** Simulations with concentrated sampling bias and simulations with distributed sampling bias are used for the performance analysis of *PhyloThin*. In figures 31 and 32 we test the robustness of the scaling step: We run *PhyloThin* on simulation results where the true tree lengths instead of mutations are used. We use $I = \{2, \cdots, n(1-p)\}$ as the index set for scaling but multiply the obtained scaling factor $s$, Eq. (3), by $c$, where $c \in \{0.5, 0.6, 0.7, \cdots, 2\}$. This allows us to examine the robustness of the scaling step. The scaling step is quite robust, ensuring that *PhyloThin* produces interpretable results even when the scaling factor $s$ is not precisely inferred. A higher percentages of oversampling and/or a smaller sample size increase the tolerance for deviations from the correct scaling factor.
Since phylogenetic trees are inferred from genomic data based on visible mutation events, we add mutations to our simulations and examine the performance of *PhyloThin* with varying mutation rate $\Theta \in \{1, 10, 100, 1.000, 10.000, 100.000\}$ (figures 34 and 35). The performance is getting worse for smaller mutation rates $\Theta$. We suggest that a mutation rate of $\Theta \geq 100$ is needed in order to use *PhyloThin*. But depending on the percentage of oversampling and the sample size also a lower mutation rate gives acceptable results. However, we recommend comparing results with the mutation-rate-sensitive-thinning option of *PhyloThin* to avoid overestimation of sampling bias. Most inferred mutation rates of the analyzed species on NCBI range from $1{,}000$ to $100{,}000$ (Fig. 21).

***Sampling bias vs. genome-wide partial selective sweeps.*** During a selective sweep, a new beneficial mutation can increases in frequency fast. Especially in prokaryotic genomes - where recombination is limited - this can lead to remarkable signatures in the phylogenetic trees: star-like coalescence. A partial and recent selective sweep can result in short external branches, similar to the pattern coming from sampling bias (Fig. 14a). As Fig. 14b indicates that the parameter range is rather small, where partial, recent sweeps and sampling bias lead to indistinguishable patterns for *PhyloThin*. However, small sample sizes (relative to the total population size) increase the problem (Fig. 14b A/B vs. C/D). Already a few generations of neutral evolution after the partial sweep produce a different pattern in the tree such that *PhyloThin* no longer classify the samples as oversampled (Fig. 14b B and D). Nevertheless, one should keep the potential problem in mind when reducing the dataset with *PhyloThin*. On the other hand, sampling bias could lead to false positive signals for a selective sweep. An analogous argumentation holds for population expansion, since the signature of population expansion in the corresponding coalescence tree is similar to a genome-wide selective sweep.

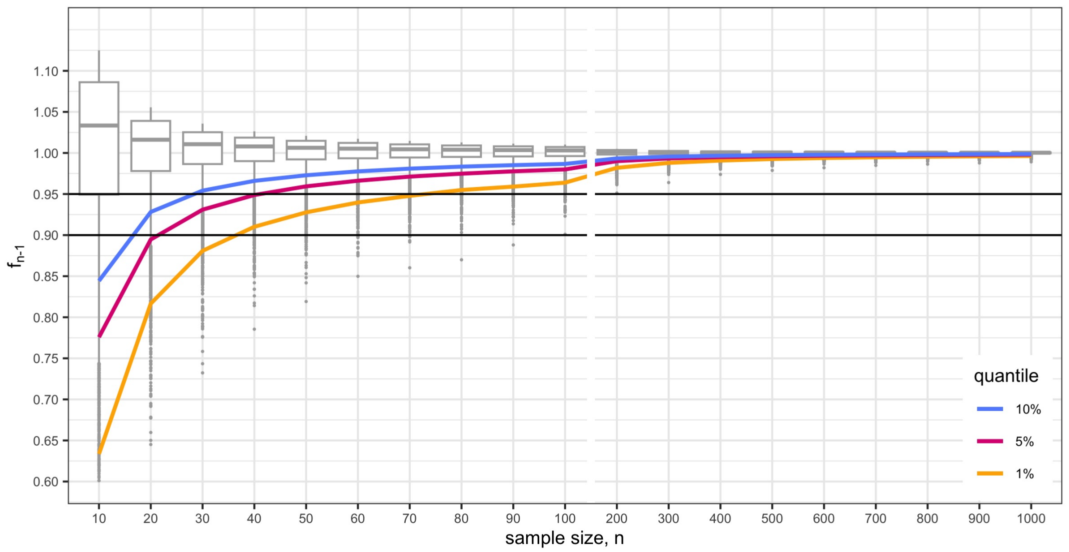


**Fig. 20.** The statistic $f_{n-1}$, Eq. (7), has been computed for neutral coalescent trees with $n$ tips. The boxplots and quantiles are based $10\,000$ replicates for each $n$. The black lines represent the thresholds $\alpha_3 = 0.95$ and $\alpha_4 = 0.9$.

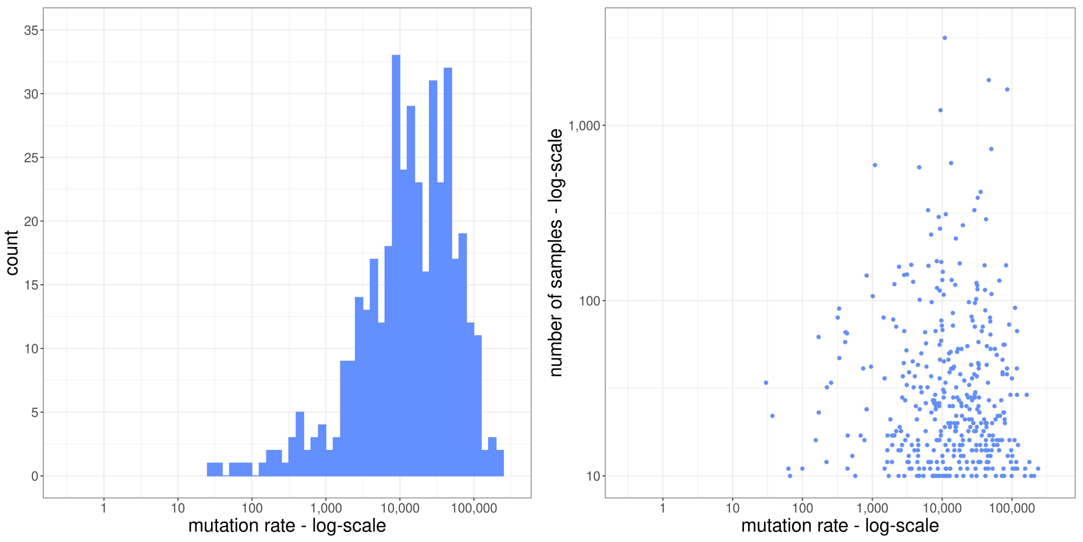


**Fig. 21.** Histogram of computed mutation rate for each analyzed species and scatter plot comparing mutation rates with sample sizes: number of SNPs in core genes divided by total branch length of the scaled ultrametric tree based on the core genes. Be aware that some axes are on logarithmic scale.

#### Supplementary Note 4: NCBI data

**Gene presence-absence.** Closely related strains sometimes carry no mutations in the core genome yet differ in the composition of their accessory genome. Therefore, as an alternative to core gene trees one might also consider trees derived from accessory gene presence-absence states. However, gene presence–absence trees have been shown to result in expanded terminal branches compared to core gene trees[82], and might not be adequately described by standard coalescent theory. In addition, gene presence-absence variation might also be caused by incompletely sequenced genomes and plasmids. Thus, one has to keep in mind that core-gene trees are better suited for detecting sampling bias and provide a higher resolution, as *PhyloThin*'s requirements (constant and frequent state transition rates, low recombination/HGT) are not always met for trees derived from gene presence–absence.

Consistent with this, phylogenies inferred from gene presence-absence in the NCBI data had longer external branches, which causes *PhyloThin* to detect less sampling bias. Oversampling counts from the two tree types were correlated, but those from presence-absence trees yielded lower estimates (Fig. 22). This discrepancy in the degree of sampling bias may thus reflect plasmid exchange and gene presence variation among strains or sequence homogenization by homologous recombination. We argue that for the purpose of our tool, core-gene trees are more likely to represent coalescent-like trees and more reliable in terms of sampling bias detection, but careful tree inference that is capable of reconstructing coalescent times is crucial. Rather than removing individual strains and their information from downstream analyses, *PhyloThin* could be used for reweighting, for example by down-weighting each genome according to the size of the oversampled group it belongs to.

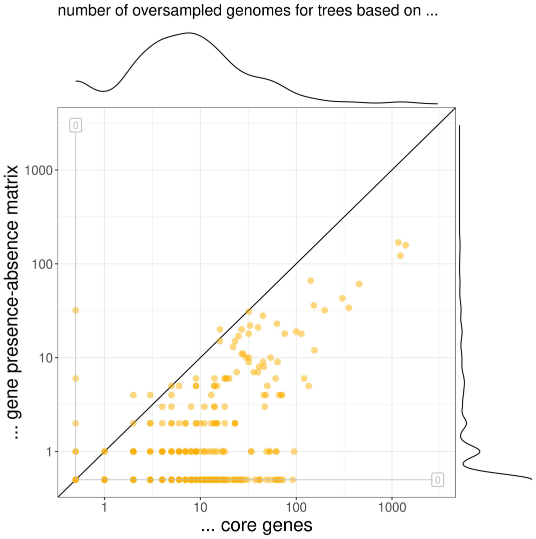


**Fig. 22. Detection of sampling bias in core genome versus full pan-genome.** Number of oversampled genomes are compared for *PhyloThin*-results based on core-gene trees and gene-presence-absence trees. Dots along the horizontal and vertical gray lines indicate species for which no oversampled genomes were detected. The number of detected oversampled genomes correlates for both type of trees (Kendall's rank correlation tau = $0.3538755$, p-value < $2.2e-16$).


***Reference.***

82. Iranzo, J et al. Gene gain and loss push prokaryotes beyond the homologous recombination barrier and accelerate genome sequence divergence. *Nature Communications*, 10(1):5376, 2019. doi: 10.1038/s41467-019-13429-2.

**Prokaryotic species from the NCBI database with at least ten full-genome-entries.** (August 4, 2023). The first column lists the species taxonomic identifier (taxid) used by NCBI, followed by the species name, the total number of genomes, and the percentage of genomes identified as oversampled in the core-gene phylogeny by *PhyloThin*. The fifth column gives the oversampling concentration ($c_{sb}$; Eq. (1)). The sixth and seventh columns report the predicted increase in genes after unbiased sampling another 100 genomes ($p_{100}$), the the predicted (core genome) number of new genes expected from sequencing the next unbiased sampled genome ($g_1$), respectively. This table extends table 1.

| taxid | species | size | oversamp. | $c_{sb}$ | $p_{100}$ | $g_1$ |
|---|---|---|---|---|---|---|
| 562 | *"Escherichia coli"* | 3152 | 36.9% | $7.0501 \cdot 10^{-3}$ | 1.56% | 6 |
| 573 | *"Klebsiella pneumoniae"* | 1811 | 67.5% | 0.0506 | 5.61% | 17.83 |
| 28901 | *"Salmonella enterica"* | 1602 | 86.6% | 0.1005 | 16.74% | 43.6 |
| 1280 | *"Staphylococcus aureus"* | 1219 | 3.7% | $1.0101 \cdot 10^{-3}$ | 2.25% | 2.03 |
| 287 | *"Pseudomonas aeruginosa"* | 732 | 62% | 0.0362 | 12.88% | 36.13 |
| 470 | *"Acinetobacter baumannii"* | 609 | 58.1% | 0.0369 | 13.96% | 29.12 |
| 520 | *"Bordetella pertussis"* | 593 | 0% | – | 0.43% | 0.17 |
| 1773 | *"Mycobacterium tuberculosis"* | 576 | 5.9% | 0.0178 | 3.8% | 2.23 |
| 210 | *"Helicobacter pylori"* | 417 | 15.8% | 0.1203 | 6.55% | 2.15 |
| 1639 | *"Listeria monocytogenes"* | 386 | 78.8% | 0.086 | 20.77% | 15.31 |
| 492670 | *"Bacillus velezensis"* | 328 | 43.3% | 0.1347 | 12.71% | 14.84 |
| 1352 | *"Enterococcus faecium"* | 328 | 34.5% | 0.0534 | 11.8% | 13.1 |
| 1423 | *"Bacillus subtilis"* | 311 | 49.2% | 0.0527 | 25.63% | 42.27 |
| 197 | *"Campylobacter jejuni"* | 300 | 45% | 0.1728 | 10.08% | 4.78 |
| 158836 | *"Enterobacter hormaechei"* | 291 | 68% | 0.0998 | 31.74% | 66.07 |
| 1314 | *"Streptococcus pyogenes"* | 269 | 57.6% | 0.0783 | 11.42% | 5.6 |
| 1351 | *"Enterococcus faecalis"* | 257 | 47.1% | 0.0241 | 19.1% | 22.43 |
| 1590 | *"Lactiplantibacillus plantarum"* | 238 | 29.4% | 0.029 | 15.68% | 17.73 |
| 1313 | *"Streptococcus pneumoniae"* | 226 | 27.4% | 0.0397 | 10.83% | 5.9 |
| 347 | *"Xanthomonas oryzae"* | 168 | 41.1% | 0.0605 | 15.33% | 12.49 |
| 1282 | *"Staphylococcus epidermidis"* | 166 | 42.8% | 0.2076 | 18.03% | 12.79 |
| 28450 | *"Burkholderia pseudomallei"* | 163 | 37.4% | 0.0514 | 27.67% | 48.03 |
| 485 | *"Neisseria gonorrhoeae"* | 160 | 26.3% | 0.0395 | 6.04% | 1.9 |
| 670 | *"Vibrio parahaemolyticus"* | 159 | 39.6% | 0.1777 | 43.1% | 92.83 |
| 1396 | *"Bacillus cereus"* | 159 | 25.2% | 0.0949 | 31.97% | 95.01 |
| 1496 | *"Clostridioides difficile"* | 158 | 60.1% | 0.0788 | 30.24% | 30.76 |
| 1719 | *"Corynebacterium pseudotuberculosis"* | 156 | 41% | 0.0764 | 1.89% | 0.62 |
| 747 | *"Pasteurella multocida"* | 146 | 68.5% | 0.1287 | 34.76% | 22.69 |
| 1311 | *"Streptococcus agalactiae"* | 141 | 19.9% | 0.1296 | 14.74% | 9.57 |
| 813 | *"Chlamydia trachomatis"* | 140 | 32.1% | 0.2303 | 5.57% | 0.7 |
| 160 | *"Treponema pallidum"* | 139 | 29.5% | 0.0805 | 28.88% | 4.6 |
| 487 | *"Neisseria meningitidis"* | 131 | 48.9% | 0.2183 | 9.76% | 4.78 |
| 1307 | *"Streptococcus suis"* | 131 | 41.2% | 0.0818 | 39.84% | 43.42 |
| 1463165 | *"Klebsiella quasipneumoniae"* | 130 | 24.6% | 0.0766 | 32.32% | 73.01 |
| 1392 | *"Bacillus anthracis"* | 128 | 35.9% | 0.1662 | 18.95% | 17.3 |
| 615 | *"Serratia marcescens"* | 126 | 25.4% | 0.4435 | 32.87% | 69.11 |
| 75985 | *"Mannheimia haemolytica"* | 124 | 37.9% | 0.074 | 11.05% | 5.47 |
| 346 | *"Xanthomonas citri"* | 123 | 74.8% | 0.2219 | 48.81% | 67.83 |
| 305 | *"Ralstonia solanacearum"* | 122 | 62.3% | 0.0705 | 48.99% | 85.97 |
| 339 | *"Xanthomonas campestris"* | 118 | 44.9% | 0.0624 | 56.17% | 80.74 |
| 446 | *"Legionella pneumophila"* | 116 | 39.7% | 0.0444 | 17.99% | 15.68 |
| 666 | *"Vibrio cholerae"* | 115 | 53.9% | 0.6758 | 58.99% | 74.85 |
| 584 | *"Proteus mirabilis"* | 114 | 28.1% | 0.0544 | 25.32% | 30.08 |
| 546 | *"Citrobacter freundii"* | 109 | 38.5% | 0.1498 | 44.15% | 105.98 |
| 623 | *"Shigella flexneri"* | 108 | 55.6% | 0.3554 | 49.8% | 80.55 |
| 29459 | *"Brucella melitensis"* | 106 | 0% | – | 1.09% | 0.44 |
| 1428 | *"Bacillus thuringiensis"* | 102 | 52% | 0.143 | 60.61% | 167.95 |
| 239935 | *"Akkermansia muciniphila"* | 101 | 72.3% | 0.0693 | 64.14% | 57.15 |
| 216816 | *"Bifidobacterium longum"* | 98 | 33.7% | 0.0833 | 25.26% | 25.51 |
| 1308 | *"Streptococcus thermophilus"* | 98 | 27.6% | 0.0997 | 25.43% | 13.11 |
| 727 | *"Haemophilus influenzae"* | 97 | 18.6% | 0.0196 | 14.48% | 7.76 |
| 40324 | *"Stenotrophomonas maltophilia"* | 91 | 4.4% | 0 | 35.67% | 70.8 |
| 519 | *"Bordetella parapertussis"* | 90 | 20% | 0.5098 | 16.93% | 8.64 |

| taxid | species | size | oversamp. | $c_{sb}$ | $p_{100}$ | $g_1$ |
|---|---|---|---|---|---|---|
| 244366 | *"Klebsiella variicola"* | 88 | 25% | 0.2078 | 48.56% | 98.29 |
| 587753 | *"Pseudomonas chlororaphis"* | 85 | 27.1% | 0.1067 | 54.58% | 121.05 |
| 1502 | *"Clostridium perfringens"* | 81 | 9.9% | 0.0357 | 35.78% | 41.59 |
| 87883 | *"Burkholderia multivorans"* | 80 | 61.3% | 0.0799 | 62.35% | 122.46 |
| 2104 | *"Mycoplasmoides pneumoniae"* | 80 | 0% | – | 4.34% | 0.47 |
| 1238 | *"Piscirickettsia salmonis"* | 80 | 82.5% | 0.1007 | 90.47% | 61.29 |
| 96345 | *"Flavobacterium psychrophilum"* | 78 | 62.8% | 0.25 | 63.26% | 32.37 |
| 36809 | *"Mycobacteroides abscessus"* | 77 | 64.9% | 0.3078 | 101.72% | 161.72 |
| 1597 | *"Lacticaseibacillus paracasei"* | 77 | 16.9% | 0.0385 | 29.76% | 28.28 |
| 1390 | *"Bacillus amyloliquefaciens"* | 76 | 30.3% | 0.0949 | 30.57% | 33.22 |
| 303 | *"Pseudomonas putida"* | 73 | 24.7% | 0.5948 | 69.64% | 248 |
| 9 | *"Buchnera aphidicola"* | 72 | 25% | 0.1634 | 27.19% | 3.05 |
| 195 | *"Campylobacter coli"* | 72 | 8.3% | 0 | 13.96% | 7.05 |
| 95486 | *"Burkholderia cenocepacia"* | 71 | 56.3% | 0.7269 | 70.35% | 200.24 |
| 28903 | *"Mycoplasmopsis bovis"* | 71 | 7% | 0.1 | 24.8% | 5.17 |
| 208962 | *"Escherichia albertii"* | 71 | 21.1% | 0.181 | 36.24% | 59.79 |
| 1358 | *"Lactococcus lactis"* | 68 | 42.6% | 0.2241 | 70.66% | 78.99 |
| 644 | *"Aeromonas hydrophila"* | 67 | 40.3% | 0.1909 | 77.66% | 125.98 |
| 283734 | *"Staphylococcus pseudintermedius"* | 67 | 25.4% | 0.1618 | 19.25% | 13.09 |
| 1491 | *"Clostridium botulinum"* | 67 | 91% | 0.6612 | 306.41% | 646.26 |
| 1134687 | *"Klebsiella michiganensis"* | 67 | 20.9% | 0.1209 | 71.81% | 185.38 |
| 35814 | *"Bordetella holmesii"* | 66 | 0% | – | 0.16% | $8.67 \cdot 10^{-2}$ |
| 263 | *"Francisella tularensis"* | 66 | 81.8% | 0.376 | 57.96% | 32.18 |
| 632 | *"Yersinia pestis"* | 65 | 0% | – | 4.3% | 2.63 |
| 564 | *"Escherichia fergusonii"* | 64 | 50% | 0.0948 | 56.83% | 86.44 |
| 235 | *"Brucella abortus"* | 62 | 43.5% | 0.2137 | 2.52% | 1.39 |
| 1613 | *"Limosilactobacillus fermentum"* | 59 | 20.3% | 0.2879 | 43.42% | 28.93 |
| 336810 | *"Candidatus Karelsulcia muelleri"* | 58 | 82.8% | 0.0133 | 13.08% | 2.18 |
| 1117645 | *"Elizabethkingia anophelis"* | 58 | 51.7% | 0.3586 | 75.58% | 98.34 |
| 2371 | *"Xylella fastidiosa"* | 57 | 54.4% | 0.2022 | 19.68% | 14.11 |
| 654 | *"Aeromonas veronii"* | 56 | 7.1% | 0 | 71.62% | 129.63 |
| 550 | *"Enterobacter cloacae"* | 56 | 23.2% | 0.1538 | 70.85% | 153.24 |
| 29519 | *"Borreliella garinii"* | 56 | 33.9% | 0.0702 | 39.16% | 5.19 |
| 582 | *"Morganella morganii"* | 55 | 41.8% | 0.1304 | 57.9% | 69.94 |
| 1717 | *"Corynebacterium diphtheriae"* | 55 | 43.6% | 0.2065 | 28.15% | 17.92 |
| 47715 | *"Lacticaseibacillus rhamnosus"* | 53 | 64.2% | 0.2727 | 35.89% | 31.86 |
| 317 | *"Pseudomonas syringae"* | 53 | 18.9% | 0.2222 | 72.47% | 154.73 |
| 1405 | *"Bacillus mycoides"* | 53 | 9.4% | 0.1 | 45.6% | 98.55 |
| 1334 | *"Streptococcus dysgalactiae"* | 53 | 35.8% | 0.1287 | 62.67% | 48.15 |
| 337 | *"Burkholderia glumae"* | 52 | 42.3% | 0.4026 | 42.77% | 72.89 |
| 548 | *"Klebsiella aerogenes"* | 51 | 19.6% | 0.1111 | 83.69% | 142.61 |
| 1685 | *"Bifidobacterium breve"* | 51 | 35.3% | 0.1373 | 45.72% | 34.05 |
| 2102 | *"Mycoplasma mycoides"* | 50 | 38% | 0.2515 | 59.68% | 17.33 |
| 173 | *"Leptospira interrogans"* | 50 | 46% | 0.3715 | 28.95% | 34.39 |
| 1653831 | *"Blattabacterium cuenoti"* | 49 | 4.1% | 0 | 13.04% | 1.28 |
| 1404 | *"Priestia megaterium"* | 47 | 29.8% | 0.2088 | 121.69% | 284.82 |
| 648 | *"Aeromonas caviae"* | 46 | 13% | 0.2 | 75.02% | 120.99 |
| 47466 | *"Borrelia miyamotoi"* | 46 | 87% | 0.45 | 82.87% | 14.78 |
| 1812935 | *"Enterobacter roggenkampii"* | 46 | 37% | 0.0735 | 105.87% | 182.9 |
| 663 | *"Vibrio alginolyticus"* | 45 | 31.1% | 0.3736 | 96.08% | 144.16 |
| 1747 | *"Cutibacterium acnes"* | 45 | 22.2% | 0.0222 | 5.75% | 3.3 |
| 358 | *"Agrobacterium tumefaciens"* | 44 | 29.5% | 0.3846 | 124.33% | 352.59 |
| 1402 | *"Bacillus licheniformis"* | 44 | 13.6% | 0.2667 | 27% | 31.22 |
| 571 | *"Klebsiella oxytoca"* | 43 | 20.9% | 0.1111 | 75.27% | 155.61 |
| 293387 | *"Bacillus altitudinis"* | 43 | 11.6% | 0.2 | 47.58% | 46.26 |
| 1599 | *"Latilactobacillus sakei"* | 43 | 25.6% | 0.0727 | 58.83% | 34.87 |
| 1402135 | *"Pseudosulfitobacter pseudonitzschiae"* | 43 | 86% | 0.7492 | 218.27% | 379.7 |
| 777 | *"Coxiella burnetii"* | 42 | 16.7% | 0.4762 | 13.77% | 5.84 |
| 34085 | *"Riemerella anatipestifer"* | 42 | 9.5% | 0.3333 | 17.79% | 12.6 |
| 28025 | *"Bifidobacterium animalis"* | 42 | 85.7% | 0.7016 | 17.41% | 15.79 |
| 624 | *"Shigella sonnei"* | 41 | 31.7% | 0.2308 | 39.27% | 38.59 |
| 294 | *"Pseudomonas fluorescens"* | 41 | 17.1% | 0.1905 | 107.68% | 348.92 |

| taxid | species | size | oversamp. | $c_{sb}$ | $p_{100}$ | $g_1$ |
|---|---|---|---|---|---|---|
| 221822 | *"Phaeobacter inhibens"* | 41 | 4.9% | 0 | 53.62% | 64.65 |
| 1767 | *"Mycobacterium intracellulare"* | 41 | 26.8% | 0.2 | 51.16% | 80.48 |
| 1764 | *"Mycobacterium avium"* | 41 | 41.5% | 0.3162 | 52.04% | 72.95 |
| 1598 | *"Limosilactobacillus reuteri"* | 41 | 17.1% | 0.1905 | 70.36% | 64.62 |
| 57706 | *"Citrobacter braakii"* | 39 | 38.5% | 0.219 | 116.38% | 157.99 |
| 48296 | *"Acinetobacter pittii"* | 39 | 7.7% | 0.3333 | 63.43% | 90.2 |
| 29575 | *"Taylorella equigenitalis"* | 39 | 64.1% | 0.5767 | 6.96% | 3.86 |
| 630 | *"Yersinia enterocolitica"* | 38 | 36.8% | 0.1429 | 54.75% | 78.97 |
| 61645 | *"Enterobacter asburiae"* | 38 | 23.7% | 0.1111 | 94.26% | 184.35 |
| 343 | *"Xanthomonas translucens"* | 38 | 18.4% | 0.1429 | 41.4% | 47.03 |
| 164546 | *"Cupriavidus taiwanensis"* | 38 | 39.5% | 0.1238 | 70.21% | 140.39 |
| 645 | *"Aeromonas salmonicida"* | 37 | 62.2% | 0.834 | 166.2% | 200.21 |
| 587 | *"Providencia rettgeri"* | 37 | 18.9% | 0.0476 | 71.13% | 113.91 |
| 55601 | *"Vibrio anguillarum"* | 37 | 18.9% | 0.2857 | 52.85% | 55.8 |
| 476 | *"Moraxella bovis"* | 37 | 40.5% | 0.1619 | 17.12% | 14.27 |
| 29486 | *"Yersinia ruckeri"* | 37 | 35.1% | 0.5769 | 59.4% | 55.56 |
| 715 | *"Actinobacillus pleuropneumoniae"* | 36 | 50% | 0.0654 | 59.95% | 33.15 |
| 28035 | *"Staphylococcus lugdunensis"* | 36 | 16.7% | 0.1333 | 34.22% | 17.48 |
| 1718 | *"Corynebacterium glutamicum"* | 36 | 69.4% | 0.2567 | 163.54% | 138.44 |
| 1406 | *"Paenibacillus polymyxa"* | 36 | 19.4% | 0.0952 | 67.72% | 122.46 |
| 837 | *"Porphyromonas gingivalis"* | 35 | 40% | 0.1758 | 75.97% | 46.11 |
| 35790 | *"Rickettsia japonica"* | 34 | 0% | – | 0.87% | 0.24 |
| 28447 | *"Clavibacter michiganensis"* | 34 | 50% | 0.1324 | 70.36% | 67.47 |
| 13373 | *"Burkholderia mallei"* | 34 | 38.2% | 0.1154 | 1.32% | 2.26 |
| 1283 | *"Staphylococcus haemolyticus"* | 34 | 23.5% | 0.1071 | 60.88% | 40.5 |
| 29518 | *"Borreliella afzelii"* | 33 | 48.5% | 0.2667 | 35.33% | 4.64 |
| 1336 | *"Streptococcus equi"* | 33 | 24.2% | 0.5357 | 38.13% | 25.06 |
| 996 | *"Flavobacterium columnare"* | 32 | 84.4% | 0.4815 | 89.47% | 72.73 |
| 83559 | *"Chlamydia suis"* | 32 | 31.3% | 0.1778 | 17.24% | 3.09 |
| 552 | *"Erwinia amylovora"* | 32 | 21.9% | 0.5238 | 12.46% | 8.12 |
| 1624 | *"Ligilactobacillus salivarius"* | 32 | 12.5% | 0 | 44.95% | 37.66 |
| 1254 | *"Pediococcus acidilactici"* | 32 | 28.1% | 0.1111 | 61.28% | 39.47 |
| 1584 | *"Lactobacillus delbrueckii"* | 31 | 6.5% | 0 | 51.74% | 29.48 |
| 731 | *"Histophilus somni"* | 30 | 86.7% | 0.4215 | 71.87% | 58.97 |
| 672 | *"Vibrio vulnificus"* | 30 | 13.3% | 0 | 100.78% | 162.61 |
| 384 | *"Rhizobium leguminosarum"* | 29 | 13.8% | 0.5 | 104.19% | 326.01 |
| 1359 | *"Lactococcus cremoris"* | 30 | 40% | 0.1212 | 82.27% | 73.2 |
| 382 | *"Sinorhizobium meliloti"* | 29 | 13.8% | 0.5 | 81.52% | 154.72 |
| 316 | *"Stutzerimonas stutzeri"* | 29 | 13.8% | 0 | 110.47% | 178.76 |
| 2588536 | *"Candidatus Methylopumilus universalis"* | 29 | 62.1% | 0.5163 | 184.78% | 53.65 |
| 196024 | *"Aeromonas dhakensis"* | 29 | 17.2% | 1 | 84.49% | 97.75 |
| 1255 | *"Pediococcus pentosaceus"* | 29 | 37.9% | 0.0909 | 79.03% | 48.01 |
| 818 | *"Bacteroides thetaiotaomicron"* | 28 | 3.6% | 1 | 64.15% | 148.43 |
| 817 | *"Bacteroides fragilis"* | 28 | 3.6% | 1 | 69.08% | 129.28 |
| 78331 | *"Mycobacterium canettii"* | 28 | 32.1% | 0.3056 | 29.37% | 26.22 |
| 730 | *"[Haemophilus] ducreyi"* | 28 | 85.7% | 0.3696 | 43.7% | 37.42 |
| 1648923 | *"Bacillus paralicheniformis"* | 28 | 32.1% | 0.1944 | 90.21% | 97.73 |
| 1639133 | *"Citrobacter portucalensis"* | 28 | 32.1% | 0.3056 | 150.64% | 240.69 |
| 1580 | *"Levilactobacillus brevis"* | 28 | 10.7% | 0 | 72.72% | 61.39 |
| 1296 | *"Mammaliicoccus sciuri"* | 28 | 14.3% | 0.5 | 72.62% | 67.28 |
| 823 | *"Parabacteroides distasonis"* | 27 | 14.8% | 0.1667 | 86.14% | 168.97 |
| 561879 | *"Bacillus safensis"* | 27 | 14.8% | 0.1667 | 68.79% | 66.82 |
| 54291 | *"Raoultella ornithinolytica"* | 27 | 7.4% | 0 | 98.2% | 157.25 |
| 380021 | *"Pseudomonas protegens"* | 27 | 18.5% | 0.2 | 47.29% | 82.58 |
| 2095 | *"Mycoplasma capricolum"* | 27 | 85.2% | 0.502 | 86.53% | 55.89 |
| 196 | *"Campylobacter fetus"* | 27 | 55.6% | 0.2762 | 57.94% | 31.84 |
| 1354 | *"Enterococcus hirae"* | 27 | 29.6% | 0.0714 | 74.44% | 57.07 |
| 1309 | *"Streptococcus mutans"* | 27 | 40.7% | 0.1818 | 58.08% | 32.4 |
| 738 | *"Glaesserella parasuis"* | 26 | 7.7% | 0 | 27.28% | 20.45 |
| 633 | *"Yersinia pseudotuberculosis"* | 26 | 46.2% | 0.1061 | 67.58% | 79.64 |
| 622 | *"Shigella dysenteriae"* | 26 | 50% | 0.3974 | 72.97% | 110.84 |
| 1744 | *"Propionibacterium freudenreichii"* | 26 | 15.4% | 0 | 42.48% | 27.29 |

| taxid | species | size | oversamp. | $c_{sb}$ | $p_{100}$ | $g_1$ |
|---|---|---|---|---|---|---|
| 714 | *"Aggregatibacter actinomycetemcomitans"* | 25 | 68% | 0.1397 | 30.32% | 32.87 |
| 57975 | *"Burkholderia thailandensis"* | 25 | 68% | 0.1471 | 226.94% | 309.17 |
| 518 | *"Bordetella bronchiseptica"* | 25 | 40% | 0.0667 | 46.34% | 52.39 |
| 47770 | *"Lactobacillus crispatus"* | 25 | 8% | 1 | 86.21% | 70.92 |
| 38313 | *"Shewanella algae"* | 25 | 16% | 0.5 | 108.42% | 101.99 |
| 274 | *"Thermus thermophilus"* | 25 | 16% | 0.5 | 46.54% | 38.23 |
| 2096 | *"Mycoplasmoides gallisepticum"* | 25 | 64% | 0.3083 | 45.83% | 14.36 |
| 1530123 | *"Acinetobacter seifertii"* | 25 | 48% | 0.697 | 107.68% | 131.44 |
| 1245 | *"Leuconostoc mesenteroides"* | 26 | 19.2% | 0 | 73.97% | 46.76 |
| 83560 | *"Chlamydia muridarum"* | 24 | 83.3% | 0.7211 | 21.54% | 9.51 |
| 83554 | *"Chlamydia psittaci"* | 24 | 54.2% | 1 | 2.63% | 1.14 |
| 718 | *"Actinobacillus equuli"* | 24 | 25% | 0.0667 | 65.8% | 38.66 |
| 61435 | *"Dehalococcoides mccartyi"* | 25 | 20% | 0 | 95% | 47.37 |
| 549 | *"Pantoea agglomerans"* | 24 | 8.3% | 0 | 134.22% | 199.37 |
| 299767 | *"Enterobacter ludwigii"* | 24 | 25% | 0.0667 | 178.45% | 211.68 |
| 29461 | *"Brucella suis"* | 24 | 16.7% | 0 | 1.38% | 1.35 |
| 85698 | *"Achromobacter xylosoxidans"* | 23 | 8.7% | 1 | 98.31% | 228.15 |
| 669 | *"Vibrio harveyi"* | 23 | 26.1% | 0.2667 | 131.34% | 185.5 |
| 56448 | *"Xanthomonas arboricola"* | 23 | 47.8% | 0.3091 | 114.99% | 149.17 |
| 299583 | *"Francisella orientalis"* | 23 | 91.3% | 1 | 3.56% | 1.33 |
| 28038 | *"Latilactobacillus curvatus"* | 23 | 0% | – | 92.37% | 64.01 |
| 511 | *"Alcaligenes faecalis"* | 22 | 13.6% | 0.3333 | 104.23% | 120.71 |
| 208224 | *"Enterobacter kobei"* | 22 | 27.3% | 0.2 | 154.14% | 236 |
| 1444770 | *"Xylella taiwanensis"* | 22 | 0% | – | 1.59% | 1.12 |
| 1408 | *"Bacillus pumilus"* | 22 | 0% | – | 53.27% | 56.29 |
| 1398 | *"Heyndrickxia coagulans"* | 22 | 50% | 0.2364 | 76.43% | 82.47 |
| 756892 | *"Acinetobacter indicus"* | 21 | 9.5% | 0 | 130.47% | 124.41 |
| 676 | *"Vibrio fluvialis"* | 21 | 0% | – | 107.69% | 123.16 |
| 65058 | *"Corynebacterium ulcerans"* | 21 | 38.1% | 0.1429 | 31.33% | 20.48 |
| 292 | *"Burkholderia cepacia"* | 21 | 14.3% | 1 | 123.12% | 365.29 |
| 2109 | *"Mycoplasmopsis synoviae"* | 21 | 33.3% | 0.2857 | 52.93% | 11.7 |
| 2098 | *"Metamycoplasma hominis"* | 21 | 0% | – | 93.41% | 19.89 |
| 2058152 | *"Klebsiella grimontii"* | 21 | 23.8% | 0 | 128.16% | 212.66 |
| 174 | *"Leptospira borgpetersenii"* | 21 | 81% | 0.3309 | 29.34% | 53.48 |
| 1303 | *"Streptococcus oralis"* | 21 | 14.3% | 0.3333 | 95.02% | 68.04 |
| 955 | *"Wolbachia pipientis"* | 20 | 45% | 0.6111 | 79.81% | 35.81 |
| 680 | *"Vibrio campbellii"* | 20 | 20% | 0 | 142.33% | 257.06 |
| 38293 | *"Photobacterium damselae"* | 20 | 15% | 0 | 148.99% | 181.98 |
| 29394 | *"Dolosigranulum pigrum"* | 20 | 10% | 0 | 35.17% | 21.21 |
| 204039 | *"Dickeya dianthicola"* | 20 | 75% | 0.6571 | 82.05% | 139.79 |
| 180957 | *"Pectobacterium brasiliense"* | 20 | 45% | 0.7778 | 135.67% | 180.88 |
| 1772 | *"Mycolicibacterium smegmatis"* | 20 | 45% | 0.4444 | 22.25% | 50.31 |
| 158 | *"Treponema denticola"* | 20 | 15% | 0 | 62.14% | 66.42 |
| 1286 | *"Staphylococcus simulans"* | 20 | 25% | 0.3 | 83.74% | 57.92 |
| 1219 | *"Prochlorococcus marinus"* | 20 | 15% | 0.3333 | 97.94% | 104.47 |
| 82996 | *"Serratia plymuthica"* | 19 | 21.1% | 0 | 87.51% | 139.82 |
| 779 | *"Ehrlichia ruminantium"* | 19 | 84.2% | 0.7667 | 52.33% | 30.97 |
| 553 | *"Pantoea ananatis"* | 19 | 0% | – | 88.07% | 118.4 |
| 40214 | *"Acinetobacter johnsonii"* | 19 | 0% | – | 162.88% | 232.46 |
| 1642 | *"Listeria innocua"* | 19 | 36.8% | 0.1429 | 94.9% | 83.31 |
| 1304 | *"Streptococcus salivarius"* | 19 | 21.1% | 0.1667 | 78.78% | 53.49 |
| 1270 | *"Micrococcus luteus"* | 19 | 26.3% | 0.3 | 123.61% | 86.47 |
| 1263550 | *"Edwardsiella piscicida"* | 19 | 47.4% | 0.3611 | 75.43% | 71.38 |
| 985002 | *"Staphylococcus argenteus"* | 18 | 77.8% | 0.7363 | 76.57% | 72.09 |
| 621 | *"Shigella boydii"* | 18 | 5.6% | 1 | 35.08% | 51.95 |
| 43767 | *"Prescottella equi"* | 18 | 11.1% | 1 | 87.89% | 102.93 |
| 33959 | *"Lactobacillus johnsonii"* | 18 | 22.2% | 0 | 135.2% | 88.21 |
| 1985254 | *"Xanthomonas phaseoli"* | 18 | 83.3% | 1 | 159.02% | 255.43 |
| 1661 | *"Trueperella pyogenes"* | 18 | 16.7% | 1 | 58.28% | 33.88 |
| 1492 | *"Clostridium butyricum"* | 18 | 27.8% | 0.2 | 56.02% | 88.06 |
| 145458 | *"Rathayibacter toxicus"* | 18 | 50% | 0.5833 | 7.27% | 6.31 |
| 1355477 | *"Bradyrhizobium diazoefficiens"* | 18 | 77.8% | 1 | 479.32% | 1,259.01 |

| taxid | species | size | oversamp. | $c_{sb}$ | $p_{100}$ | $g_1$ |
|---|---|---|---|---|---|---|
| 1328 | *"Streptococcus anginosus"* | 18 | 38.9% | 0.3333 | 140.33% | 105.55 |
| 103855 | *"Bordetella hinzii"* | 18 | 38.9% | 0.3333 | 51.57% | 66.57 |
| 783 | *"Rickettsia rickettsii"* | 17 | 82.4% | 1 | 7.17% | 10.08 |
| 729 | *"Haemophilus parainfluenzae"* | 17 | 0% | – | 97.51% | 59.92 |
| 580165 | *"Bacillus cytotoxicus"* | 17 | 52.9% | 0.4444 | 26.18% | 49.57 |
| 542 | *"Zymomonas mobilis"* | 17 | 82.4% | 1 | 159.87% | 247.9 |
| 29385 | *"Staphylococcus saprophyticus"* | 17 | 29.4% | 0.4 | 106.85% | 67.26 |
| 1905730 | *"Pectobacterium parmentieri"* | 17 | 64.7% | 0.2364 | 182.94% | 219.4 |
| 1901 | *"Streptomyces clavuligerus"* | 17 | 0% | – | 5.43% | 14.87 |
| 1821960 | *"Edwardsiella anguillarum"* | 17 | 82.4% | 0.5055 | 262.17% | 199.03 |
| 1596 | *"Lactobacillus gasseri"* | 17 | 82.4% | 0.5604 | 341.74% | 197.13 |
| 1579 | *"Lactobacillus acidophilus"* | 17 | 82.4% | 1 | 27.56% | 40.18 |
| 1176649 | *"Agrobacterium fabrum"* | 17 | 58.8% | 0.8 | 191.36% | 272.04 |
| 1089444 | *"Dickeya solani"* | 17 | 82.4% | 1 | 58.9% | 88.49 |
| 853 | *"Faecalibacterium prausnitzii"* | 16 | 12.5% | 0 | 132.44% | 178.7 |
| 80867 | *"Paracidovorax avenae"* | 16 | 0% | – | 51.22% | 89.31 |
| 728 | *"Avibacterium paragallinarum"* | 16 | 18.8% | 0.3333 | 23.92% | 23.75 |
| 67780 | *"Edwardsiella ictaluri"* | 16 | 31.3% | 0.2 | 25.96% | 27.82 |
| 588 | *"Providencia stuartii"* | 16 | 81.3% | 0.6154 | 184.3% | 338.52 |
| 545 | *"Citrobacter koseri"* | 16 | 50% | 0.1071 | 216.09% | 228.53 |
| 488447 | *"Burkholderia contaminans"* | 16 | 37.5% | 0.1333 | 126.5% | 359.34 |
| 480 | *"Moraxella catarrhalis"* | 16 | 25% | 0 | 55.27% | 24.46 |
| 47877 | *"Pseudomonas amygdali"* | 16 | 18.8% | 0.3333 | 149.37% | 260.97 |
| 44008 | *"Enterococcus cecorum"* | 16 | 12.5% | 1 | 88.48% | 77.45 |
| 43080 | *"Sulfolobus islandicus"* | 16 | 12.5% | 0 | 59.47% | 59.43 |
| 334542 | *"Rhodococcus qingshengii"* | 16 | 18.8% | 0.3333 | 163.62% | 280.93 |
| 314275 | *"Alteromonas mediterranea"* | 16 | 43.8% | 0.2857 | 259.45% | 315.79 |
| 28141 | *"Cronobacter sakazakii"* | 16 | 43.8% | 0.3333 | 120.14% | 145.03 |
| 28108 | *"Alteromonas macleodii"* | 16 | 25% | 0.5 | 183.43% | 236.22 |
| 199 | *"Campylobacter concisus"* | 16 | 18.8% | 0 | 134.89% | 83.25 |
| 171284 | *"Mycoplasmopsis cynos"* | 16 | 0% | – | 66.75% | 47.14 |
| 1290 | *"Staphylococcus hominis"* | 16 | 12.5% | 0 | 116.7% | 70.43 |
| 1063 | *"Cereibacter sphaeroides"* | 16 | 81.3% | 1 | 185.15% | 429.2 |
| 664662 | *"Borreliella bavariensis"* | 15 | 46.7% | 0.4762 | 79.87% | 24.26 |
| 554 | *"Pectobacterium carotovorum"* | 15 | 0% | – | 117.76% | 165.3 |
| 396 | *"Rhizobium phaseoli"* | 15 | 53.3% | 0.5357 | 119.51% | 224.53 |
| 29430 | *"Acinetobacter haemolyticus"* | 15 | 6.7% | 1 | 130.79% | 138.81 |
| 28116 | *"Bacteroides ovatus"* | 15 | 46.7% | 0.3333 | 160.03% | 414.54 |
| 2702 | *"Gardnerella vaginalis"* | 15 | 80% | 0.8333 | 112.68% | 100.53 |
| 2267275 | *"Bartonella krasnovii"* | 15 | 40% | 0.2667 | 33.52% | 19.64 |
| 204042 | *"Dickeya zeae"* | 15 | 13.3% | 0 | 109.12% | 134.74 |
| 2026186 | *"Bacillus paranthracis"* | 15 | 26.7% | 0.1667 | 92.69% | 156.67 |
| 1589 | *"Lactiplantibacillus pentosus"* | 15 | 53.3% | 0.5357 | 86.52% | 114.39 |
| 1587 | *"Lactobacillus helveticus"* | 15 | 46.7% | 0.4286 | 103.75% | 78.4 |
| 1574161 | *"Proteus terrae"* | 15 | 33.3% | 0.4 | 118.25% | 136.31 |
| 152480 | *"Burkholderia ambifaria"* | 15 | 53.3% | 0.5357 | 144.72% | 325.56 |
| 1421 | *"Lysinibacillus sphaericus"* | 15 | 80% | 0.8333 | 259.69% | 642.24 |
| 137591 | *"Weissella cibaria"* | 15 | 60% | 0.2222 | 155.88% | 101.36 |
| 1201031 | *"Pectobacterium aroidearum"* | 15 | 60% | 0.3333 | 141.29% | 178.17 |
| 91844 | *"Candidatus Portiera aleyrodidarum"* | 14 | 64.3% | 0.7778 | 3.15% | 0.95 |
| 83655 | *"Leclercia adecarboxylata"* | 14 | 0% | – | 128.99% | 181.77 |
| 76759 | *"Pseudomonas monteilii"* | 14 | 78.6% | 0.4364 | 548.07% | 761.04 |
| 60552 | *"Burkholderia vietnamiensis"* | 14 | 35.7% | 0.2 | 202.3% | 340.04 |
| 438 | *"Acetobacter pasteurianus"* | 14 | 78.6% | 0.8182 | 301.02% | 299.9 |
| 38323 | *"Bartonella henselae"* | 14 | 50% | 0.7143 | 10.65% | 8.38 |
| 34 | *"Myxococcus xanthus"* | 14 | 35.7% | 0.3 | 27.11% | 88.39 |
| 29494 | *"Vibrio furnissii"* | 14 | 21.4% | 0.3333 | 146.97% | 177.58 |
| 28132 | *"Prevotella melaninogenica"* | 14 | 28.6% | 1 | 154.79% | 153.71 |
| 28131 | *"Prevotella intermedia"* | 14 | 28.6% | 0.1667 | 146.81% | 112.19 |
| 260554 | *"Bacillus halotolerans"* | 14 | 0% | – | 70.48% | 80.53 |
| 2035 | *"Curtobacterium flaccumfaciens"* | 14 | 21.4% | 0 | 83.58% | 104.68 |
| 1855823 | *"Macrococcus canis"* | 14 | 14.3% | 1 | 98.9% | 86.02 |

| taxid | species | size | oversamp. | $c_{sb}$ | $p_{100}$ | $g_1$ |
|---|---|---|---|---|---|---|
| 1681 | *"Bifidobacterium bifidum"* | 14 | 35.7% | 0.1 | 79.44% | 47.47 |
| 139 | *"Borreliella burgdorferi"* | 14 | 50% | 0.3333 | 19.61% | 11.51 |
| 1363 | *"Lactococcus garvieae"* | 14 | 78.6% | 0.8182 | 391% | 315.66 |
| 1302 | *"Streptococcus gordonii"* | 14 | 42.9% | 0.2 | 105.5% | 69.29 |
| 1281 | *"Staphylococcus carnosus"* | 14 | 71.4% | 0.3556 | 62.45% | 73.79 |
| 103816 | *"Rhodococcus pyridinivorans"* | 14 | 0% | – | 152.19% | 253.52 |
| 80869 | *"Paracidovorax citrulli"* | 13 | 76.9% | 0.4 | 128.04% | 164.45 |
| 651 | *"Aeromonas media"* | 13 | 15.4% | 0 | 183.86% | 212.29 |
| 62322 | *"Shewanella baltica"* | 13 | 30.8% | 0.5 | 191.19% | 241.42 |
| 56454 | *"Xanthomonas hortorum"* | 13 | 38.5% | 0.1 | 106.99% | 164.99 |
| 43769 | *"Corynebacterium propinquum"* | 13 | 46.2% | 0.4 | 41.1% | 35.96 |
| 37734 | *"Enterococcus casseliflavus"* | 13 | 15.4% | 0 | 139.64% | 168.75 |
| 2287 | *"Saccharolobus solfataricus"* | 13 | 46.2% | 0.4667 | 19.17% | 28.27 |
| 2027919 | *"Enterobacter cloacae complex sp."* | 13 | 15.4% | 1 | 156.1% | 277.13 |
| 155322 | *"Bacillus toyonensis"* | 13 | 15.4% | 0 | 99.92% | 168.14 |
| 1310165 | *"Ralstonia pseudosolanacearum"* | 13 | 7.7% | 1 | 65.67% | 111.9 |
| 1299 | *"Deinococcus radiodurans"* | 13 | 15.4% | 0 | 1.76% | 2.18 |
| 1076 | *"Rhodopseudomonas palustris"* | 13 | 23.1% | 1 | 169.13% | 237.66 |
| 85991 | *"Chlamydia pecorum"* | 12 | 25% | 0.3333 | 5.8% | 1.79 |
| 851 | *"Fusobacterium nucleatum"* | 12 | 16.7% | 1 | 126.4% | 122.59 |
| 83558 | *"Chlamydia pneumoniae"* | 12 | 75% | 0.5 | 6.42% | 5.78 |
| 83555 | *"Chlamydia abortus"* | 12 | 50% | 1 | 4.13% | 2.88 |
| 821 | *"Phocaeicola vulgatus"* | 12 | 33.3% | 0.1667 | 245.88% | 399.15 |
| 648995 | *"Agrobacterium pusense"* | 12 | 25% | 0.3333 | 308.09% | 418.4 |
| 54736 | *"Salmonella bongori"* | 12 | 50% | 1 | 110.34% | 116.57 |
| 47883 | *"Pseudomonas synxantha"* | 12 | 8.3% | 1 | 144.87% | 268.69 |
| 450367 | *"Peribacillus frigoritolerans"* | 12 | 16.7% | 0 | 160.77% | 264.13 |
| 34021 | *"Candidatus Liberibacter asiaticus"* | 12 | 8.3% | 1 | 4.39% | 1.79 |
| 28095 | *"Burkholderia gladioli"* | 12 | 25% | 0.3333 | 147.35% | 343.71 |
| 253 | *"Chryseobacterium indologenes"* | 12 | 58.3% | 0.5238 | 273.39% | 494.65 |
| 246432 | *"Staphylococcus equorum"* | 12 | 16.7% | 0 | 123.15% | 93.14 |
| 2209 | *"Methanosarcina mazei"* | 12 | 25% | 0.3333 | 122.25% | 119.87 |
| 2151 | *"Mesoplasma florum"* | 12 | 33.3% | 0.1667 | 167.63% | 42.45 |
| 213 | *"Helicobacter cinaedi"* | 12 | 75% | 1 | 494.71% | 268.29 |
| 201 | *"Campylobacter lari"* | 12 | 25% | 0.3333 | 117.88% | 53.68 |
| 1748967 | *"Citrobacter cronae"* | 12 | 33.3% | 1 | 97.93% | 88.02 |
| 1694 | *"Bifidobacterium pseudolongum"* | 12 | 16.7% | 0 | 103.08% | 51.33 |
| 1680 | *"Bifidobacterium adolescentis"* | 12 | 25% | 0.3333 | 226.49% | 142.82 |
| 1509 | *"Clostridium sporogenes"* | 12 | 8.3% | 1 | 107.26% | 151.89 |
| 1499973 | *"Escherichia marmotae"* | 12 | 16.7% | 1 | 130.27% | 178.03 |
| 1292 | *"Staphylococcus warneri"* | 12 | 75% | 1 | 88.26% | 88.48 |
| 126385 | *"Providencia alcalifaciens"* | 12 | 75% | 1 | 423.9% | 387.38 |
| 106654 | *"Acinetobacter nosocomialis"* | 12 | 16.7% | 0 | 156.81% | 172.92 |
| 803 | *"Bartonella quintana"* | 11 | 45.5% | 0.3 | 15.49% | 8.22 |
| 782 | *"Rickettsia prowazekii"* | 11 | 0% | – | 0.15% | $9.76 \cdot 10^{-2}$ |
| 636 | *"Edwardsiella tarda"* | 11 | 72.7% | 0.5714 | 225.25% | 228.79 |
| 59814 | *"Pantoea dispersa"* | 11 | 72.7% | 1 | 213.24% | 406.07 |
| 585 | *"Proteus vulgaris"* | 11 | 9.1% | 1 | 130.89% | 182.87 |
| 54571 | *"Streptomyces venezuelae"* | 11 | 27.3% | 1 | 174.45% | 628.42 |
| 52584 | *"Brachyspira pilosicoli"* | 11 | 27.3% | 0.3333 | 98.03% | 70.36 |
| 42858 | *"Mammaliicoccus lentus"* | 11 | 9.1% | 1 | 66.11% | 48.96 |
| 40215 | *"Acinetobacter junii"* | 11 | 0% | – | 150.73% | 152.58 |
| 357276 | *"Phocaeicola dorei"* | 11 | 54.5% | 0.1333 | 312.31% | 416.2 |
| 35703 | *"Citrobacter amalonaticus"* | 11 | 36.4% | 0.1667 | 315.96% | 391.18 |
| 300181 | *"Serratia ureilytica"* | 11 | 0% | – | 86.05% | 107.46 |
| 28090 | *"Acinetobacter lwoffii"* | 11 | 72.7% | 0.75 | 250.61% | 466.74 |
| 2285 | *"Sulfolobus acidocaldarius"* | 11 | 18.2% | 1 | 3.5% | 3.6 |
| 2133 | *"Spiroplasma citri"* | 11 | 0% | – | 66.72% | 38.32 |
| 2099 | *"Mesomycoplasma hyopneumoniae"* | 11 | 27.3% | 0 | 62.28% | 19.29 |
| 1922217 | *"Candidatus Erwinia haradaeae"* | 11 | 18.2% | 0 | 23.08% | 5.09 |
| 1778540 | *"Dickeya fangzhongdai"* | 11 | 36.4% | 0.1667 | 146.9% | 165.23 |
| 1774 | *"Mycobacteroides chelonae"* | 11 | 36.4% | 0.1667 | 191.47% | 246.5 |

| taxid | species | size | oversamp. | $c_{sb}$ | $p_{100}$ | $g_1$ |
|---|---|---|---|---|---|---|
| 1768 | *"Mycobacterium kansasii"* | 11 | 72.7% | 0.5714 | 33.76% | 76.89 |
| 1648 | *"Erysipelothrix rhusiopathiae"* | 11 | 72.7% | 1 | 67.99% | 41.05 |
| 140 | *"Borrelia hermsii"* | 11 | 72.7% | 0.5714 | 100.88% | 48.3 |
| 1288 | *"Staphylococcus xylosus"* | 11 | 0% | – | 147.12% | 89.1 |
| 1183401 | *"Agrobacterium fabacearum"* | 11 | 0% | – | 201.14% | 338.47 |
| 114186 | *"Candidatus Carsonella ruddii"* | 11 | 27.3% | 0.3333 | 9.04% | 1.1 |
| 106592 | *"Ensifer adhaerens"* | 11 | 36.4% | 0.3333 | 260.71% | 663.21 |
| 985762 | *"Staphylococcus agnetis"* | 10 | 30% | 0.3333 | 100.04% | 76.52 |
| 930166 | *"Pseudomonas brassicacearum"* | 10 | 70% | 1 | 253.09% | 530.54 |
| 881260 | *"Enterobacter bugandensis"* | 10 | 50% | 0.6 | 142.67% | 152.35 |
| 84112 | *"Eggerthella lenta"* | 10 | 0% | – | 55.28% | 72.6 |
| 81947 | *"Vagococcus lutrae"* | 10 | 20% | 0 | 106.07% | 66.58 |
| 781 | *"Rickettsia conorii"* | 10 | 70% | 0.5238 | 151.19% | 101.87 |
| 53345 | *"Enterococcus durans"* | 10 | 40% | 0.1667 | 160.35% | 158.9 |
| 408 | *"Methylorubrum extorquens"* | 10 | 30% | 0.3333 | 333.9% | 533.06 |
| 38284 | *"Corynebacterium accolens"* | 10 | 30% | 0.3333 | 110.84% | 70.9 |
| 37919 | *"Rhodococcus opacus"* | 10 | 40% | 1 | 252.8% | 839.32 |
| 36855 | *"Brucella canis"* | 10 | 0% | – | 0.48% | 0.75 |
| 364410 | *"Granulibacter bethesdensis"* | 10 | 30% | 0.3333 | 76.31% | 47.81 |
| 33964 | *"Leuconostoc citreum"* | 10 | 20% | 0 | 133.96% | 81.57 |
| 33033 | *"Parvimonas micra"* | 10 | 20% | 1 | 82.28% | 43.65 |
| 32046 | *"Synechococcus elongatus"* | 10 | 80% | 0.75 | 49.82% | 39.64 |
| 315405 | *"Streptococcus gallolyticus"* | 10 | 40% | 0.1667 | 227.38% | 182.07 |
| 29388 | *"Staphylococcus capitis"* | 10 | 80% | 0.75 | 65.69% | 51.81 |
| 28198 | *"Aliarcobacter cryaerophilus"* | 10 | 20% | 0 | 280.2% | 171.33 |
| 28182 | *"Leptospira noguchii"* | 10 | 20% | 1 | 89.44% | 121.26 |
| 28026 | *"Bifidobacterium pseudocatenulatum"* | 10 | 0% | – | 126.21% | 83.39 |
| 2527775 | *"Polynucleobacter paneuropaeus"* | 10 | 40% | 1 | 377.52% | 144.32 |
| 251701 | *"Pseudomonas syringae"* | 10 | 80% | 1 | 145.49% | 254.43 |
| 1886 | *"Streptomyces albidoflavus"* | 10 | 80% | 1 | 43.5% | 81.74 |
| 1833 | *"Rhodococcus erythropolis"* | 10 | 20% | 1 | 206.98% | 342.28 |
| 164514 | *"Blattabacterium punctulatus"* | 10 | 30% | 0 | 7.26% | 1.52 |
| 162 | *"Treponema phagedenis"* | 10 | 20% | 0 | 19.95% | 23.17 |
| 1604 | *"Lactobacillus amylovorus"* | 10 | 0% | – | 179.55% | 123.22 |
| 1520 | *"Clostridium beijerinckii"* | 10 | 30% | 0.3333 | 208.75% | 370.42 |
| 1452 | *"Bacillus atrophaeus"* | 10 | 20% | 0 | 104.12% | 120.17 |
| 1422 | *"Geobacillus stearothermophilus"* | 10 | 40% | 0.5 | 134.25% | 181.93 |
| 134821 | *"Ureaplasma parvum"* | 10 | 0% | – | 51.52% | 9.69 |
| 1160784 | *"Candidatus Nasuia deltocephalinicola"* | 10 | 0% | – | 79.02% | 3.7 |

### Supplementary Note 5: Supplementary figures

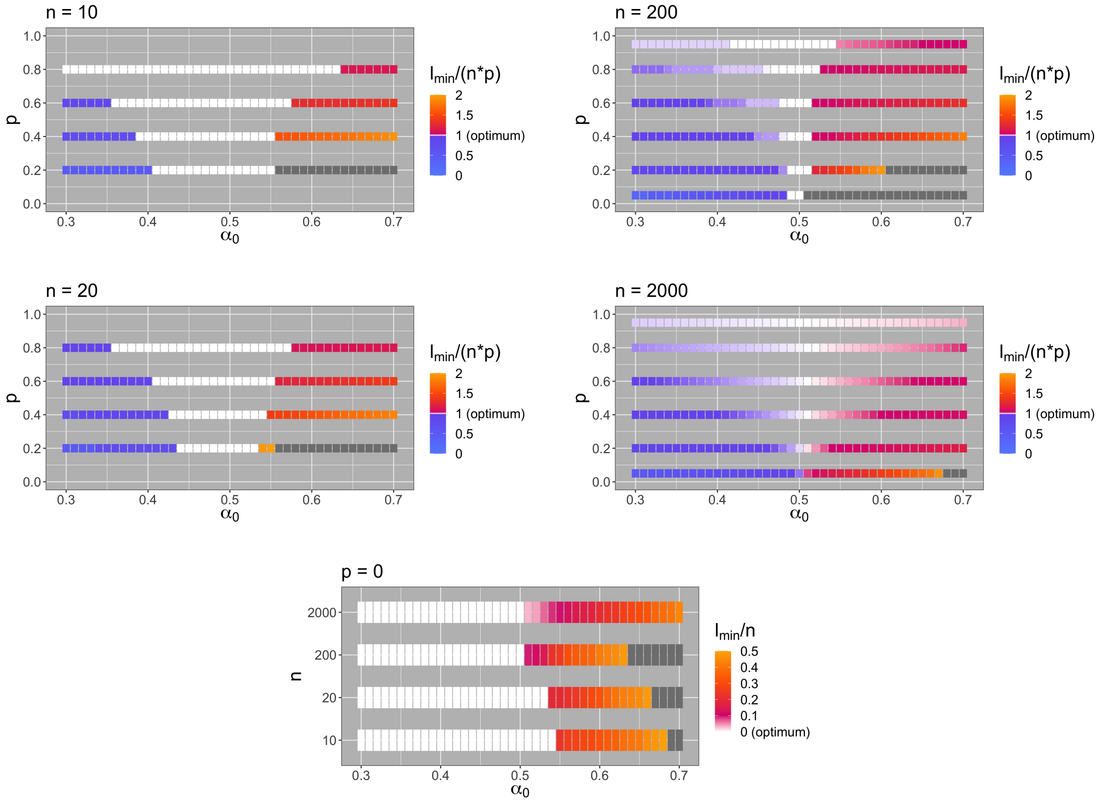


**Fig. 23.** *PhyloThin* is optimized for $\alpha_0 \in [0.3, 0.7]$ by looking at $l^{(1)}_{\text{min}}$, Eq. (11), for simulation results ($N = 100\,000$, $n \in \{10, 20, 200, 2000\}$, $p \in \{0, 0.05, 0.2, 0.4, 0.6, 0.8, 0.95\}$). The total population tree has been generated $10$ times. For each $p$ the most related subtree consisting of at least $np + 1$ tips is used to represent the oversampled group (simulation setting: concentrated sampling bias). The results shown are based on the mean of $100$ simulation-runs for each combination of parameters (i.e., the remaining random sample is different for each run).
For optimal $\alpha_0$ the following should hold: $\frac{l_{\text{min}}}{np} \gtrapprox 1$ for $p > 0$ and $\frac{l_{\text{min}}}{n} \approx 0$ for $p = 0$. The color scheme (given in the legend on the right hand side of each panel) is the same for each panel. The optimal value ($0$ or $1$) is indicated in white color. Gray colored areas correspond to higher values than given in the legend.

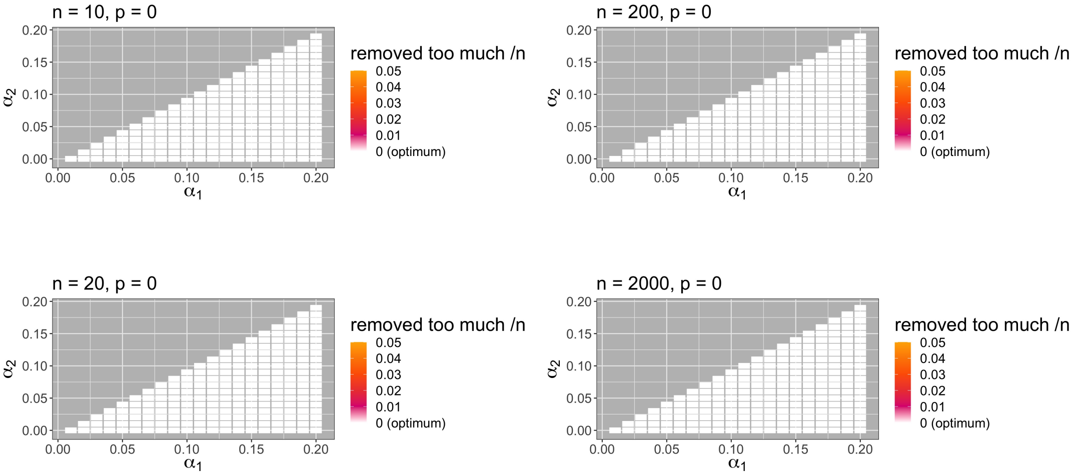


**Fig. 24.** *PhyloThin* is optimized for $\alpha_1 \in (0, 0.2]$ and $\alpha_2 \in [0, \alpha_1)$ by looking at the number of false positives for simulation results ($N = 100\,000$, $n \in \{10, 20, 200, 2000\}$, $p = 0$). The simulation setting is the same as described in the caption of Fig. 23.
For optimal $\alpha_1$ and $\alpha_2$, $\frac{\#\text{removed too much}}{n} \approx 0$ for $p = 0$. The color scheme (given in the legend on the right hand side of each panel) is the same for each column. The optimal value ($0$) is indicated in white color. Gray colored areas correspond to higher values than given in the legend.

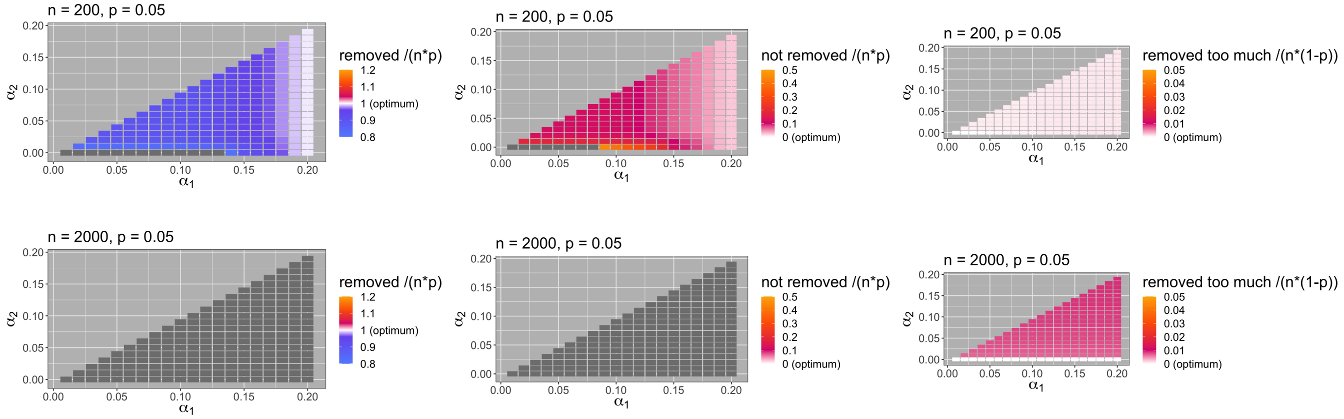


**Fig. 25.** *PhyloThin* is optimized for $\alpha_1 \in (0, 0.2]$ and $\alpha_2 \in [0, \alpha_1)$ by looking at the number of removed tips, the number of false negatives and false positives for simulation results ($N = 100\,000$, $n \in \{200, 2000\}$, $p = 0.05$). The simulation setting is the same as described in the caption of Fig. 23.
For optimal $\alpha_1$ and $\alpha_2$ the following should hold: $\frac{\#\text{removed}}{np} \approx 1$, $\frac{\#\text{not removed}}{np} \approx 0$, $\frac{\#\text{removed too much}}{n(1-p)} \approx 0$ for $p > 0$. The color scheme (given in the legend on the right hand side of each panel) is the same for each column. The optimal value ($0$ or $1$) is indicated in white color. Gray colored areas correspond to higher values than given in the legend.

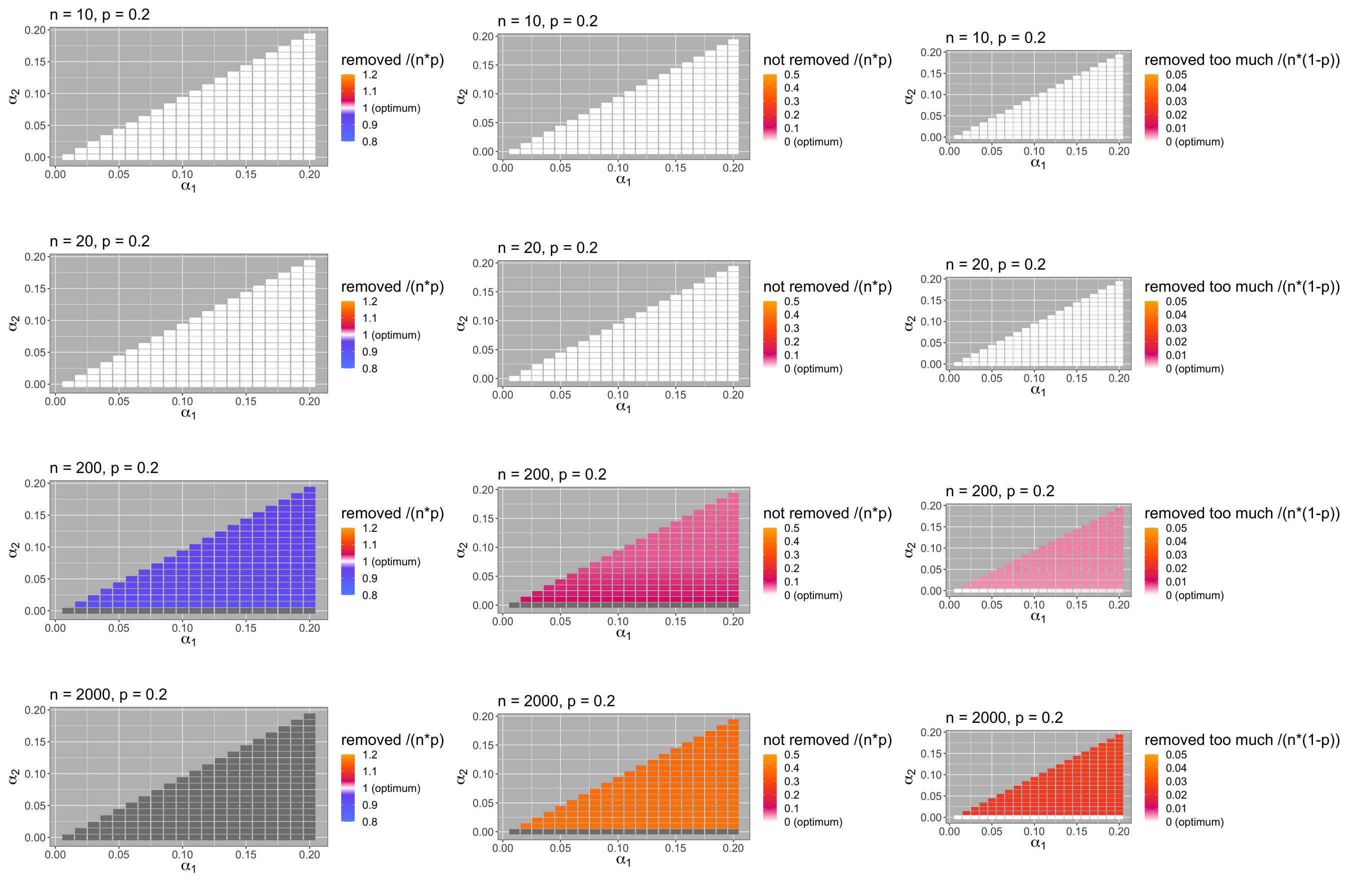

**Fig. 26.** *PhyloThin* is optimized for $\alpha_1 \in (0, 0.2]$ and $\alpha_2 \in [0, \alpha_1)$ by looking at the number of removed tips, the number of false negatives and false positives for simulation results ($N = 100\,000$, $n \in \{10, 20, 200, 2000\}$, $p = 0.2$). The simulation setting is the same as described in the caption of Fig. 23.
For optimal $\alpha_1$ and $\alpha_2$ the following should hold: $\frac{\#\text{removed}}{np} \approx 1$, $\frac{\#\text{not removed}}{np} \approx 0$, $\frac{\#\text{removed too much}}{n(1-p)} \approx 0$ for $p > 0$. The color scheme (given in the legend on the right hand side of each panel) is the same for each column. The optimal value (0 or 1) is indicated in white color. Gray colored areas correspond to higher values than given in the legend.

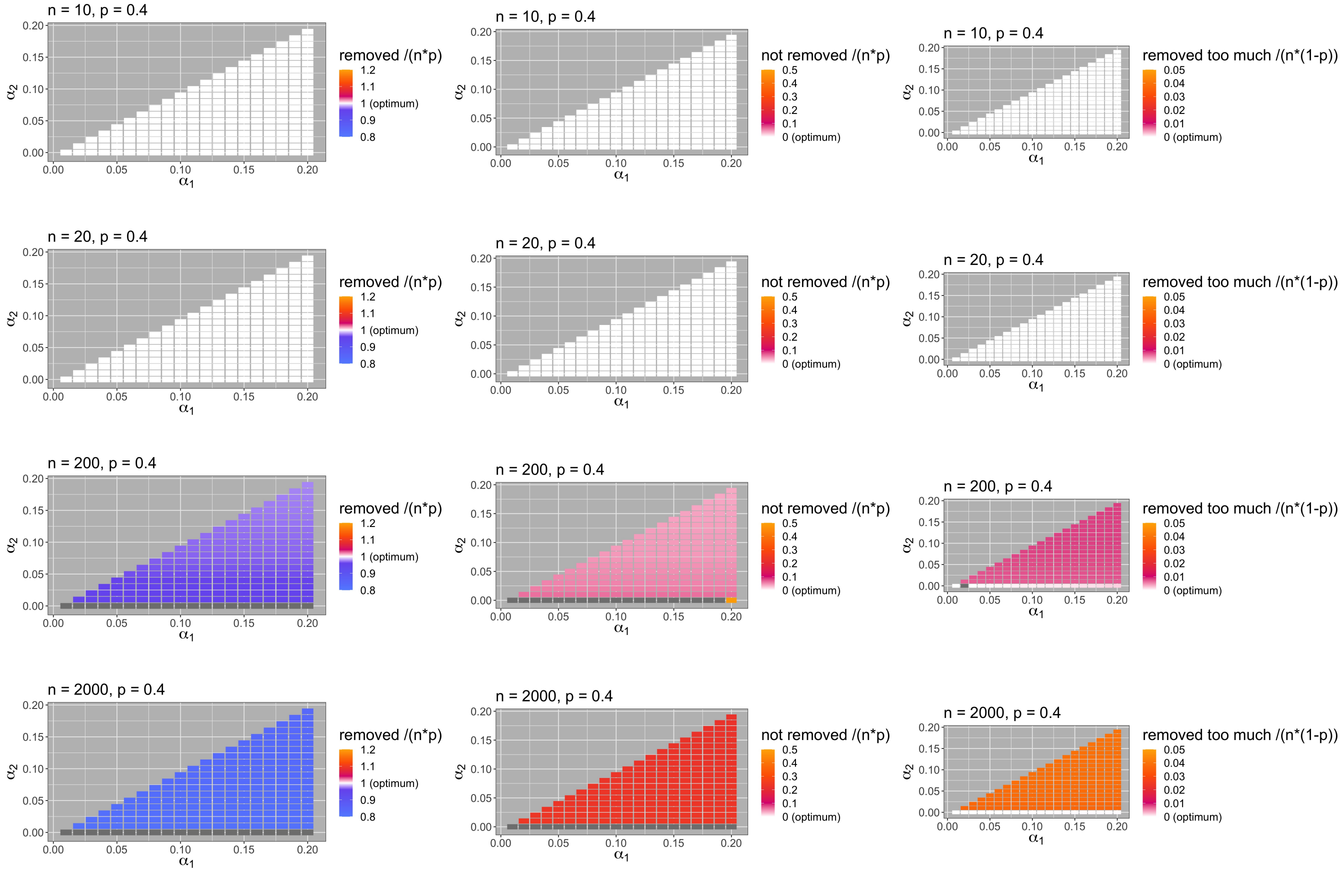


**Fig. 27.** *PhyloThin* is optimized for $\alpha_1 \in (0, 0.2]$ and $\alpha_2 \in [0, \alpha_1)$ by looking at the number of removed tips, the number of false negatives and false positives for simulation results ($N = 100\,000$, $n \in \{10, 20, 200, 2000\}$, $p = 0.4$). The simulation setting is the same as described in the caption of Fig. 23.

For optimal $\alpha_1$ and $\alpha_2$ the following should hold: $\frac{\#\text{removed}}{np} \approx 1$, $\frac{\#\text{not removed}}{np} \approx 0$, $\frac{\#\text{removed too much}}{n(1-p)} \approx 0$ for $p > 0$. The color scheme (given in the legend on the right hand side of each panel) is the same for each column. The optimal value ($0$ or $1$) is indicated in white color. Gray colored areas correspond to higher values than given in the legend.

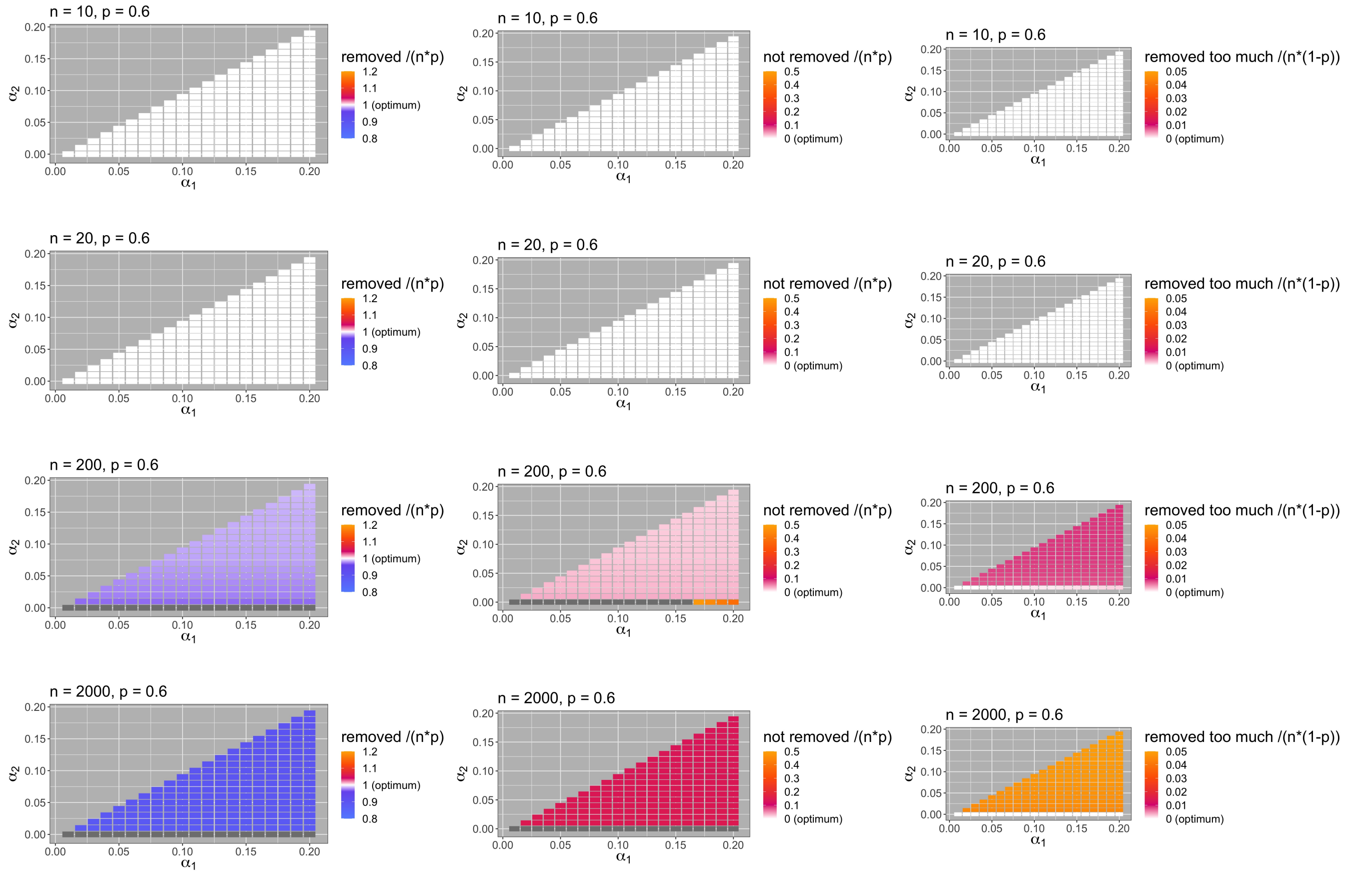


**Fig. 28.** *PhyloThin* is optimized for $\alpha_1 \in (0, 0.2]$ and $\alpha_2 \in [0, \alpha_1)$ by looking at the number of removed tips, the number of false negatives and false positives for simulation results ($N = 100\,000$, $n \in \{10, 20, 200, 2000\}$, $p = 0.6$). The simulation setting is the same as described in the caption of Fig. 23.
For optimal $\alpha_1$ and $\alpha_2$ the following should hold: $\frac{\#\text{removed}}{np} \approx 1$, $\frac{\#\text{not removed}}{np} \approx 0$, $\frac{\#\text{removed too much}}{n(1-p)} \approx 0$ for $p > 0$. The color scheme (given in the legend on the right hand side of each panel) is the same for each column. The optimal value ($0$ or $1$) is indicated in white color. Gray colored areas correspond to higher values than given in the legend.

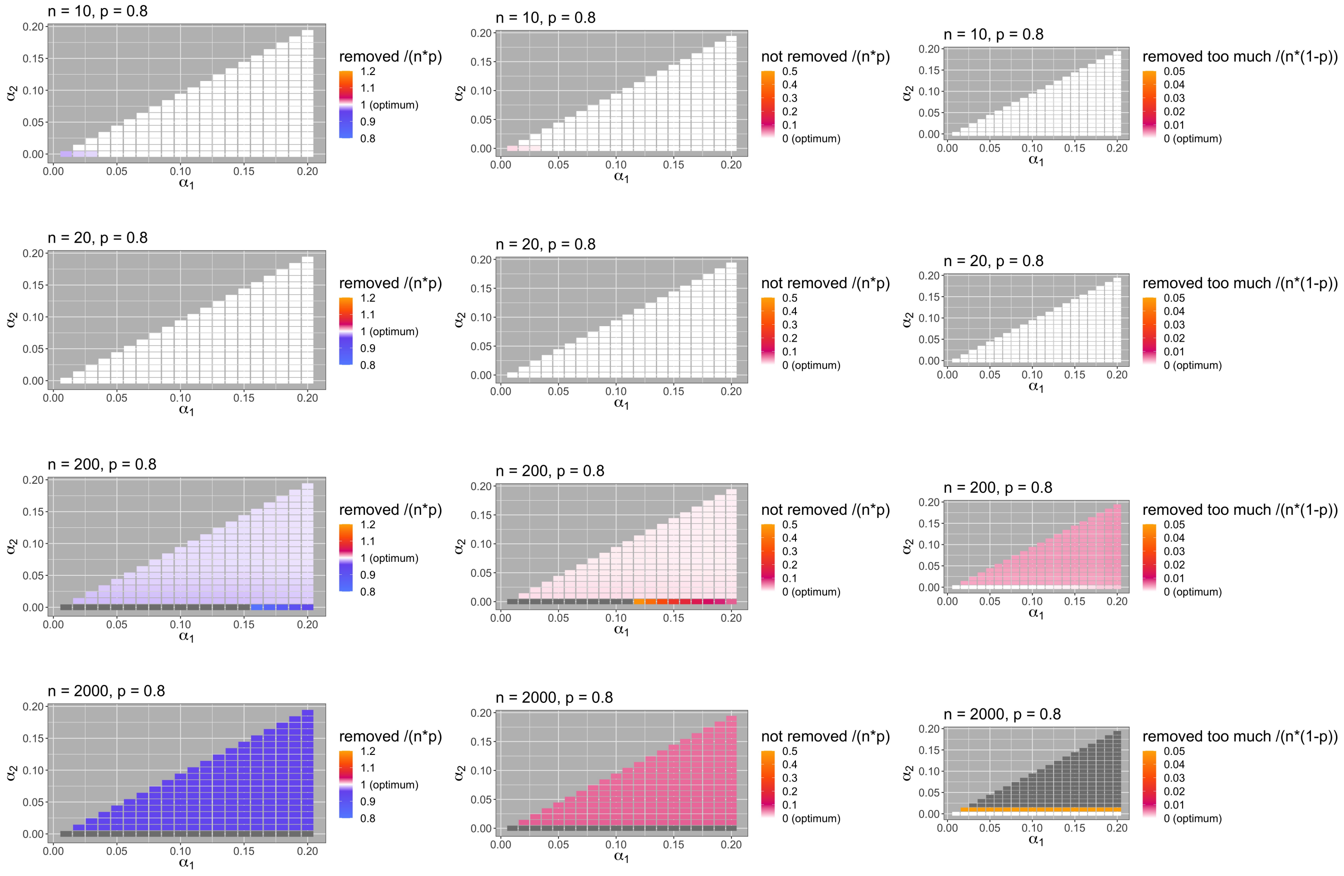


**Fig. 29.** *PhyloThin* is optimized for $\alpha_1 \in (0, 0.2]$ and $\alpha_2 \in [0, \alpha_1)$ by looking at the number of removed tips, the number of false negatives and false positives for simulation results ($N = 100\,000$, $n \in \{10, 20, 200, 2000\}$, $p = 0.8$). The simulation setting is the same as described in the caption of Fig. 23.
For optimal $\alpha_1$ and $\alpha_2$ the following should hold: $\frac{\#\text{removed}}{np} \approx 1$, $\frac{\#\text{not removed}}{np} \approx 0$, $\frac{\#\text{removed too much}}{n(1-p)} \approx 0$ for $p > 0$. The color scheme (given in the legend on the right hand side of each panel) is the same for each column. The optimal value (0 or 1) is indicated in white color. Gray colored areas correspond to higher values than given in the legend.

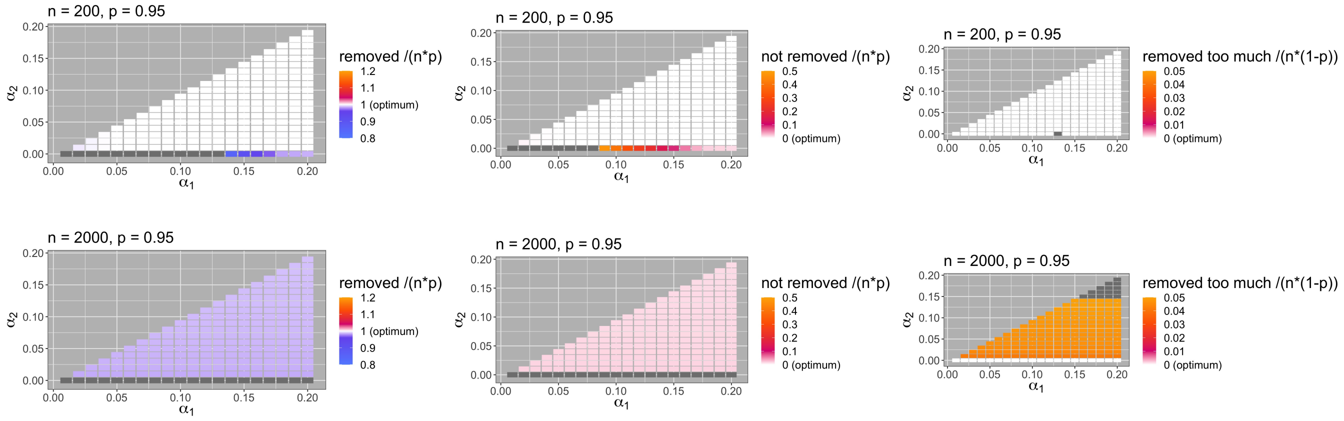


**Fig. 30.** *PhyloThin* is optimized for $\alpha_1 \in (0, 0.2]$ and $\alpha_2 \in [0, \alpha_1)$ by looking at the number of removed tips, the number of false negatives and false positives for simulation results ($N = 100\,000$, $n \in \{200, 2000\}$, $p = 0.95$). The simulation setting is the same as described in the caption of Fig. 23.
For optimal $\alpha_1$ and $\alpha_2$ the following should hold: $\frac{\#\text{removed}}{np} \approx 1$, $\frac{\#\text{not removed}}{np} \approx 0$, $\frac{\#\text{removed too much}}{n(1-p)} \approx 0$ for $p > 0$. The color scheme (given in the legend on the right hand side of each panel) is the same for each column. The optimal value (0 or 1) is indicated in white color. Gray colored areas correspond to higher values than given in the legend.

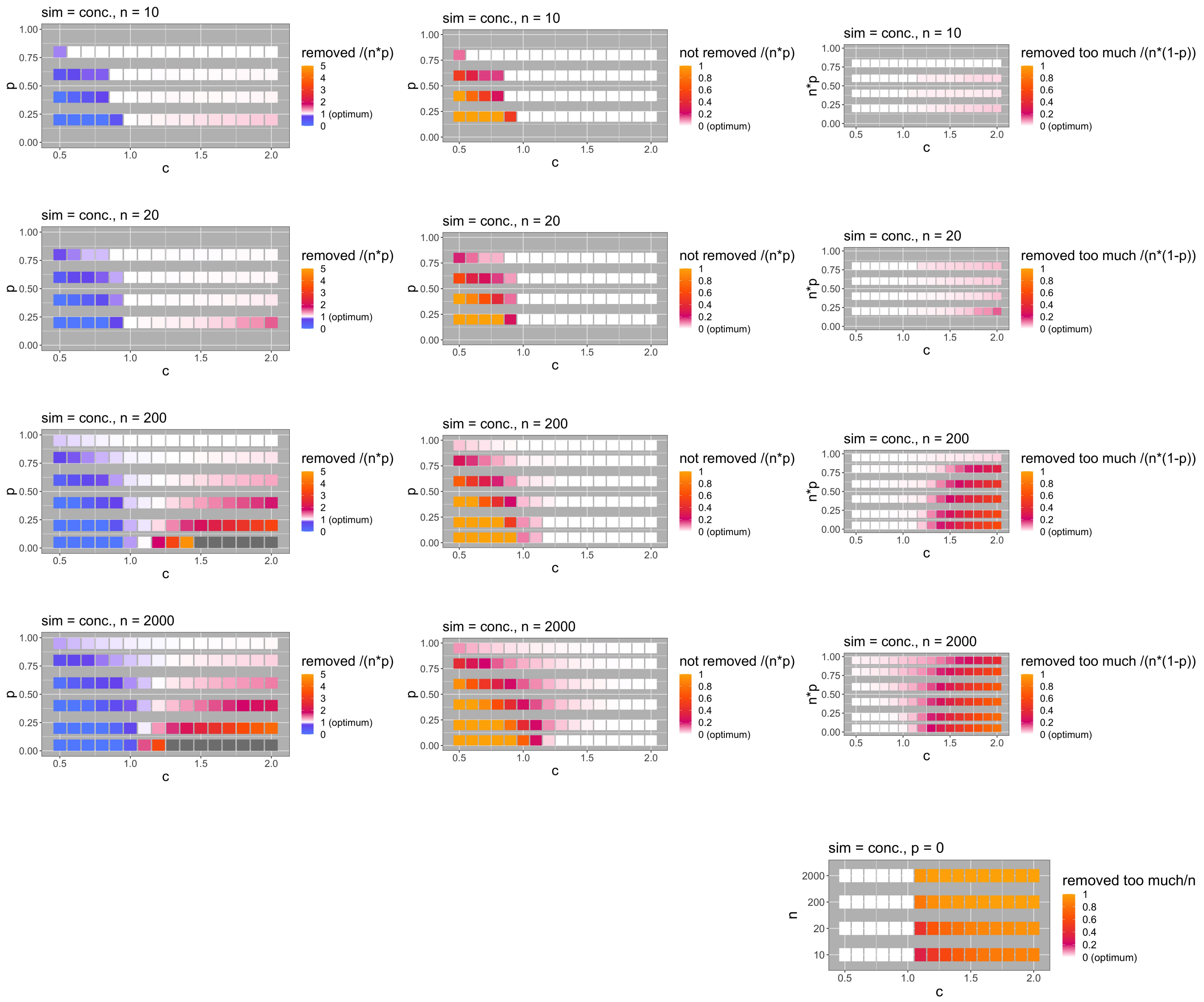


**Fig. 31.** The robustness of the scaling step of *PhyloThin* is examined on simulation results with concentrated (conc.) sampling bias (where the true tree lengths instead of mutations are used) by using $I = \{2, \cdots, n(1-p)\}$ as the index set for scaling but multiply the obtained scaling factor $s$, Eq. (3), by $c$, where $c \in \{0.5, 0.6, 0.7, \cdots, 2\}$. The simulation setting is the same as described in the caption of Fig. 23.

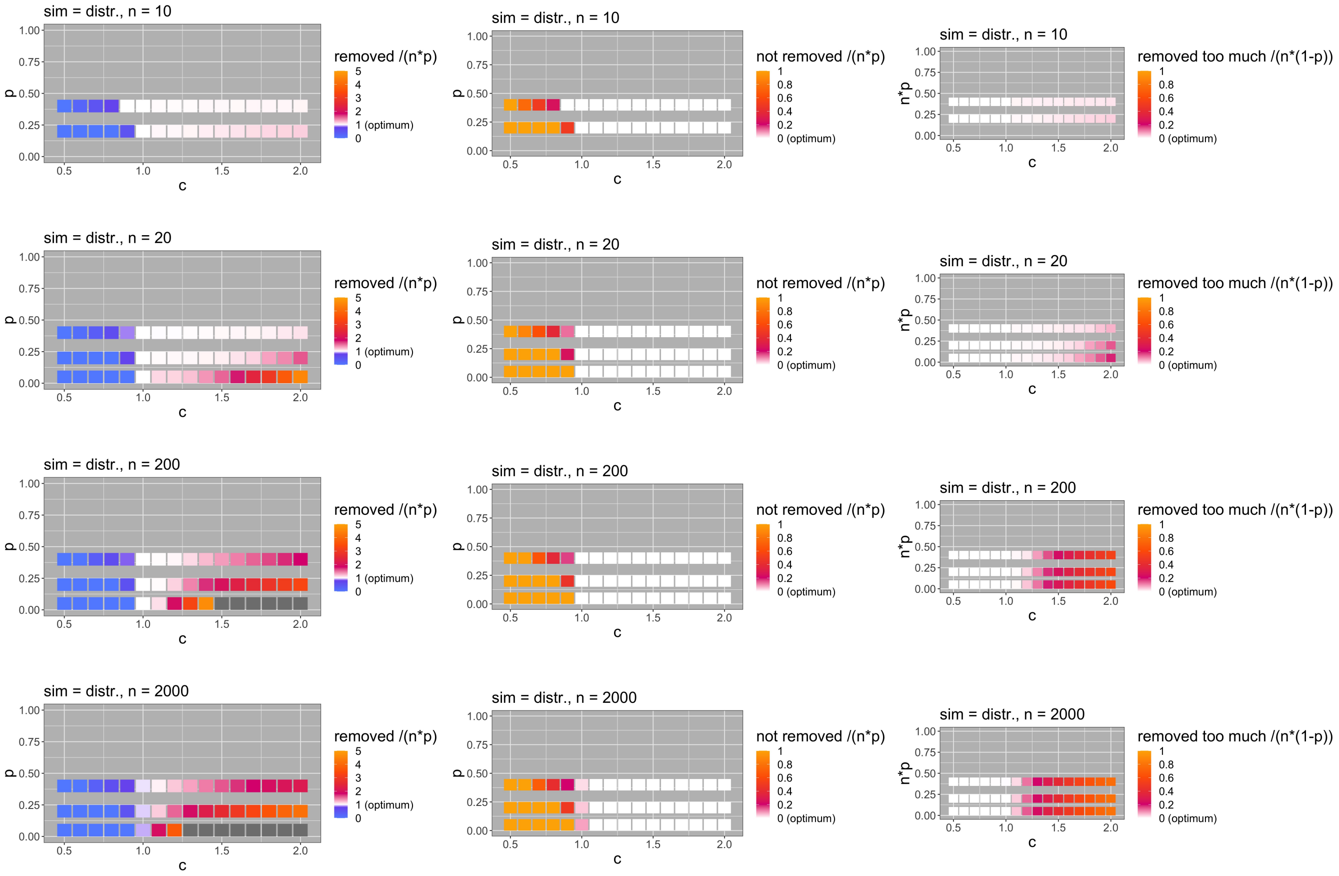


**Fig. 32.** The robustness of the scaling step of *PhyloThin* is examined on simulation results with distributed (distr.) sampling bias (where the true tree lengths instead of mutations are used) by using $I = \{2, \cdots, n(1-p)\}$ as the index set for scaling but multiply the obtained scaling factor $s$, Eq. (3), by $c$, where $c \in \{0.5, 0.6, 0.7, \cdots, 2\}$. The simulation setting is the same as described in the caption of Fig. 23.

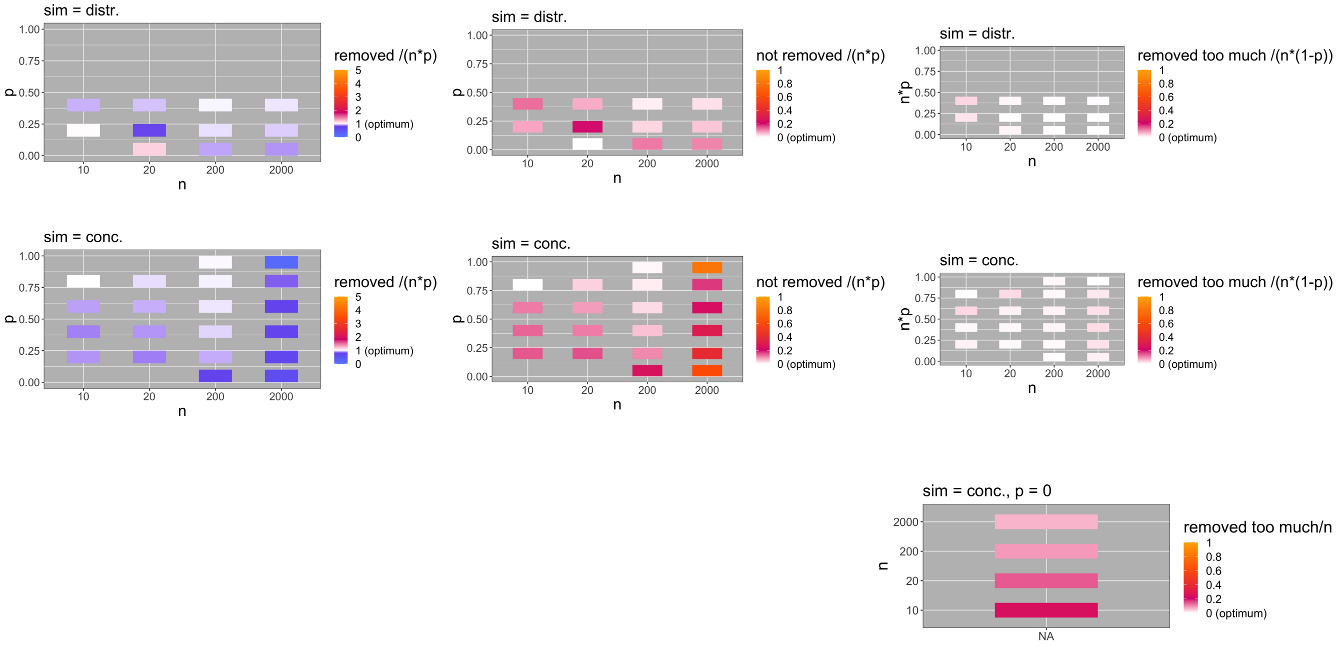


**Fig. 33.** The performance of *PhyloThin* is examined on simulation results with concentrated (conc.) and distributed (distr.) sampling bias (where the true tree lengths instead of mutations are used, $\Theta \to \infty$). The simulation setting is the same as described in the caption of Fig. 23.

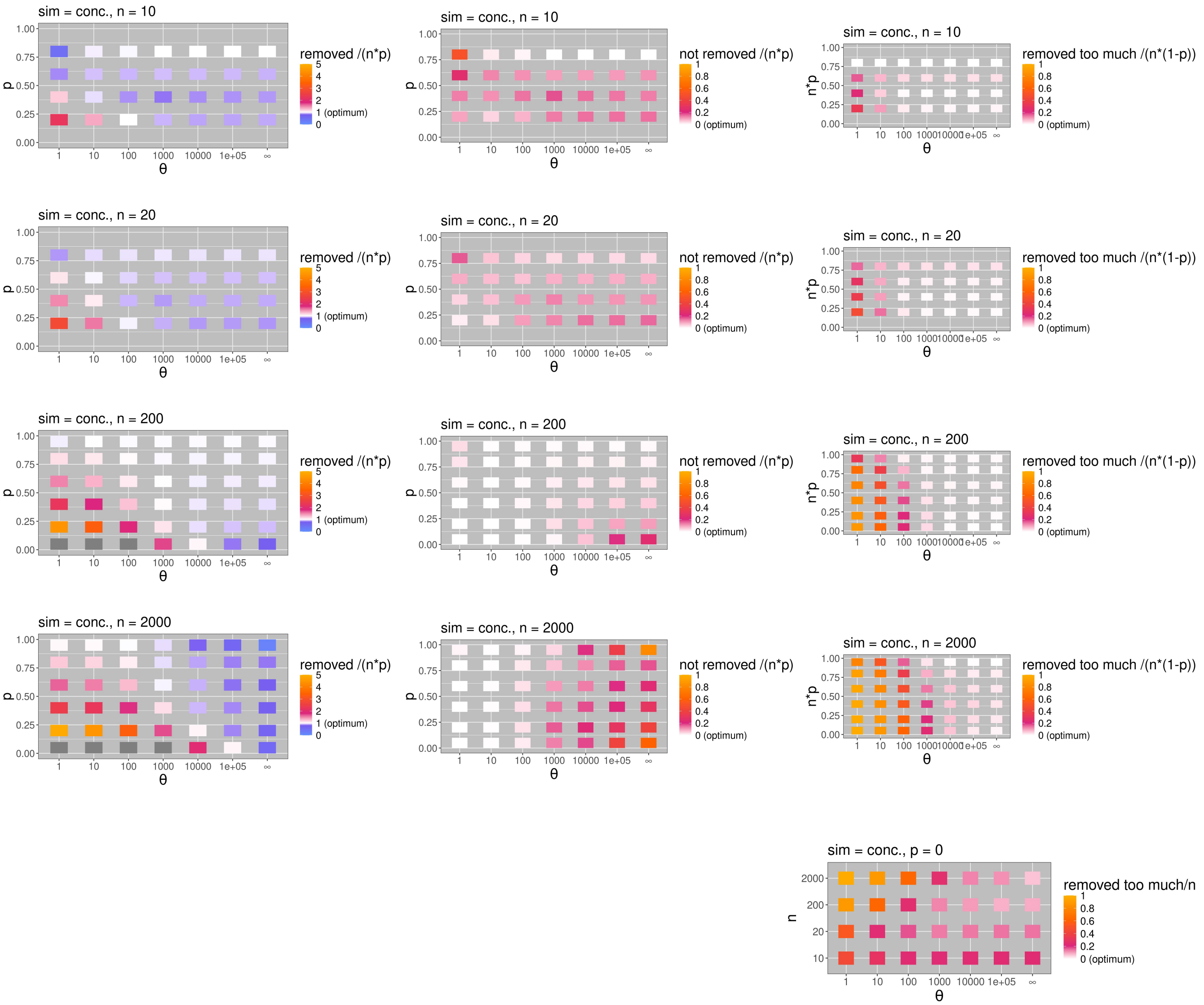


**Fig. 34.** The performance of *PhyloThin* is examined on simulation results with concentrated (conc.) sampling bias by varying the mutation rate $\Theta \in \{1, 10, 100, 1.000, 10.000, 100.000\}$ ($N = 100000$, $n \in \{10, 20, 200, 2000\}$, $p \in \{0, 0.05, 0.2, 0.4, 0.6, 0.8, 0.95\}$) and comparing to simulation results where the true tree lengths instead of mutations are used ($\Theta \to \infty$). The simulation setting is the same as described in the caption of Fig. 23.

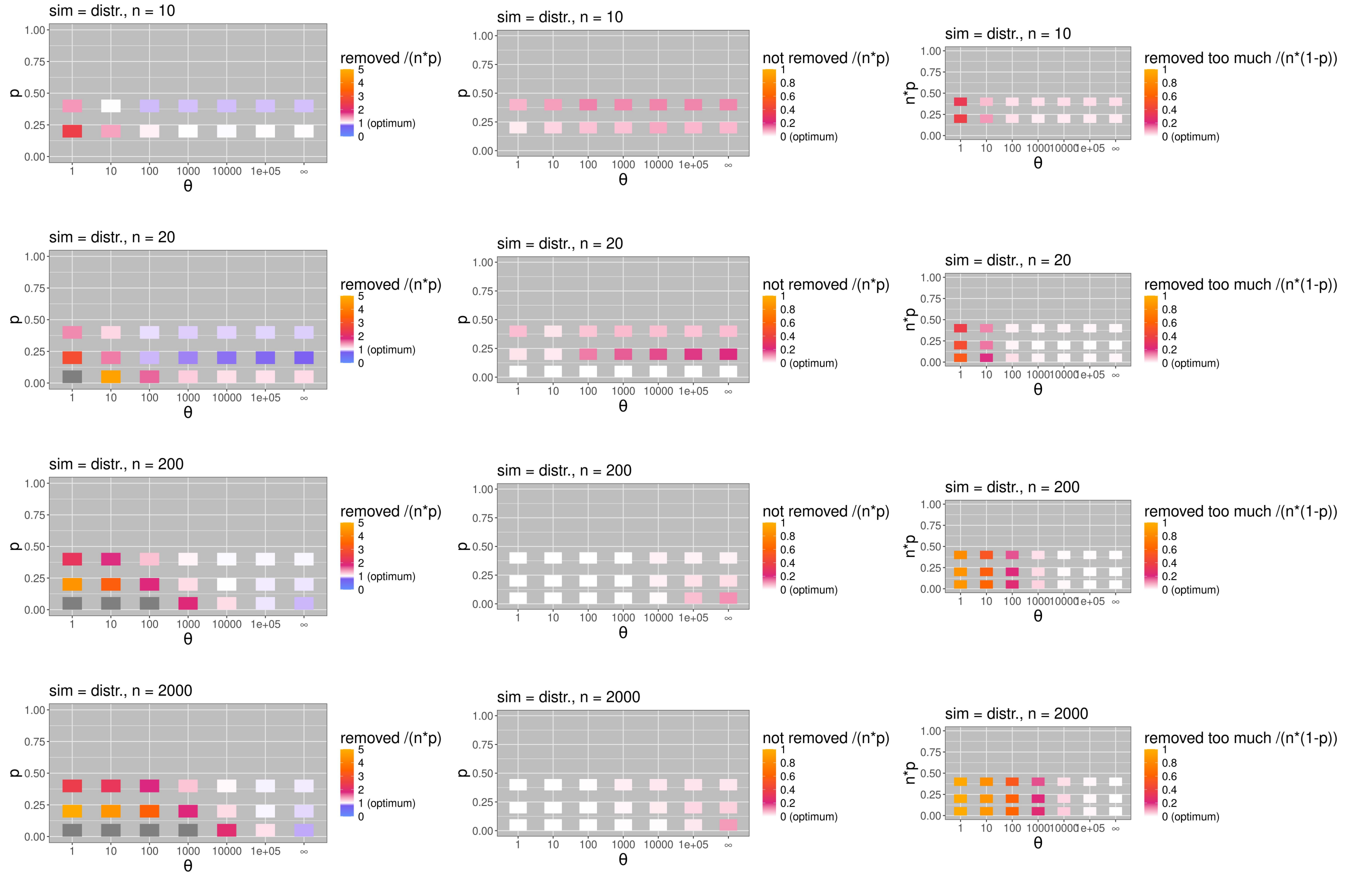

**Fig. 35.** The performance of *PhyloThin* is examined on simulation results with distributed (distr.) sampling bias by varying the mutation rate $\Theta \in \{1, 10, 100, 1.000, 10.000, 100.000\}$ ($N = 10000$, $n \in \{10, 20, 200, 2000\}$, $p \in \{0.05, 0.2, 0.4\}$) and comparing to simulation results where the true tree lengths instead of mutations are used ($\Theta \to \infty$). The simulation setting is the same as described in the caption of Fig. 23.
